# Serosolver: an open source tool to infer epidemiological and immunological dynamics from serological data

**DOI:** 10.1101/730069

**Authors:** James A. Hay, Amanda Minter, Kylie Ainslie, Justin Lessler, Adam J. Kucharski, Steven Riley

## Abstract

We present a flexible, open source R package designed to obtain additional biological and epidemiological insights from commonly available serological datasets. Analysis of serological responses against pathogens with multiple strains such as influenza pose a specific statistical challenge because observed antibody responses measured in serological assays depend both on unobserved prior infections and the resulting cross-reactive antibody dynamics that these infections generate. We provide a general modelling framework to jointly infer these two typically confounded biological processes using antibody titres against current and historical strains. We do this by linking latent infection dynamics with a mechanistic model of antibody dynamics that generates expected antibody titres over time. This makes it possible to use observations of antibodies in serological assays to infer an individual’s infection history as well as the parameters of the antibody process model. Our aim is to provide a flexible inference package that can be applied to a range of datasets studying different viruses over different timescales. We present two case studies to illustrate how our model can infer key immunological parameters, such as antibody titre boosting, waning and cross-reaction, and well as latent epidemiological processes such as attack rates and age-stratified infection risk.

## Introduction

Serological assays measure the interaction of a virus with the antibody repertoire of an individual host [1]. Originally developed in the mid-20th Century, assays based on haemagglutination inhibition (HI) and viral neutralization (VN) are still widely used and are highly repeatable within the same lab [2]. These assays can be setup relatively easily when viral culture systems are in place and require no specialist kits. Usually, serum is diluted in 2- or 4-fold steps. Limiting dilutions with higher titres indicate a stronger antibody response, whereas titres below the limit of detection indicate the absence of a significant response. In influenza, ‘lower than 1:10’ is often the minimum reading and dilutions of 1:1024 or higher indicate strong antibody responses. The longevity of antibodies such as IgG make serological assays a key tool in epidemiological surveillance, particularly where virological assays are not possible and symptoms are non-specific or non-existent [3–6]. When only a single sample is available for an individual, a threshold titre is often used as evidence of prior exposure or protection or both, for example the commonly used threshold of 1:40 for influenza [7, 8].

Paired blood samples and serological assays using known circulating strains can be used to estimate exposure within a specific period of time. Samples taken before and after an influenza season for which the main circulating strain is known can therefore be used to infer attack rates [9–11]. Samples are usually processed as a pair to limit the impact of between batch variability in testing. A *≥* 4-fold rise in titre against the circulating strain (homologous titre) between the pre- and post-exposure samples is typically assumed to be evidence of influenza infection. Because there is a degree of subjectivity in the characterization of a sample being a limiting dilution, a *≥* 4-fold difference, within a 2-fold dilution scheme, is deemed to be more robust against human error than a *≥* 2-fold difference in assessing the presence of haemaglutination (for HI) or cell death (for VN) in each well of the assay plate [12, 13]. However, a Bayesian analysis of titre rise data has suggested that the somewhat arbitrary fourfold rise hides a substantial number of lesser true titre rises that may represent missed infections [14]. Individual-level differences in age, infection history, time between exposure and measurement, and virus-specific effects likely all play a role in generating sub-fourfold titre rises [15–18].

Cross-reactivity complicates the interpretation of serological results when an individual may have been exposed to two or more antigenically related viruses. Two pathogens are considered antigenically related if exposure to one generates a cross-reactive antibody response to the other in a serological assay. For example, antibody responses against one dengue virus serotype can cross-react with another [19], as well as other flaviviruses such as Zika virus [20, 21]. Moreover, sequential lineages of individual influenza A subtypes cross-react with their precursors and progeny [22]. One popular use of HI assays is to assess the cross-reactivity between current influenza A sub-types. Naive ferrets are inoculated with one of a panel of current reference strains to produce virus-specific serum. HI titres are then measured for potentially novel viruses using stocks of these reference ‘antisera’ [23].

Recently, there have been a number of initiatives to refine the analyses of commonly available serological data. Antigenic cartography was developed to reduce complex tables of HI readings for novel viruses and reference antisera to two dimensional space, visualised as an ‘antigenic map’ [23–25]. An individual’s entire antibody repertoire against an antigenically variable pathogen can be then projected as a surface over these antigenic maps, with the height of the surface at any specific point indicating the expected titre for that individual against a strain at that location in the map [26]. These ‘antibody landscapes’ can be used to generate biological insight by investigating how antibody profiles develop over an individual’s life [27]. Further, compartmental transmission models can be defined with explicit strata for each serological assay result and used to test hypothesis about the interplay of social mixing and pre-existing immunity [28]. These approaches retain much of the information present in the magnitude of an assay measurement that may be lost when using seroconversion and seropositivity thresholds.

Here, we present the R package `serosolver`, which is the latest version of a code base developed specifically to increase the epidemiological insight available from serological assays [27, 29]. `Serosolver` takes assay results from one or more serum samples for an individual, which may have been tested against one or more related viral strains, and infers a history of infections for that individual that is consistent with the observed titres. It can jointly estimate the process parameters for the antibody kinetic process by simultaneously inferring infection histories for many people. We use a Bayesian approach and obtain correlated samples from the posterior densities for infection histories and process parameters. The required assumptions for some priors are straightforward and may incorporate previously observed immunological phenomena. Prior assumptions for infection histories and the process that generates them can also be incorporated, but can require additional justification, as we shall discuss.

The basic inference challenge can be summarised as follows. For a given set of serological data (***Y***, which may include assay measurements against one or more strains), we wish to obtain the joint posterior distribution of the process parameters (*θ*), individual infection histories (***Z***) and temporal probability of infection in the population (*ϕ*). This posterior is proportional to three components: (i) the observation and antibody process models *f* (***Y***_***i***_*|****Z***_***i***_, *θ*), which give the likelihood of observing a set of titres ***Y***_***i***_ for each individual *i* at serum sampling times (*t*_*i*_), given infection history ***Z***_***i***_ and process parameters *θ*; (ii) the transmission level *P* (*Z*_*i,j*_|*ϕ*_*j*_), which gives the probability of individual *i* having an infection with the strain circulating in time period *j*, given population infection probability *ϕ*_*j*_; and (iii) the prior level, giving the prior probability for the process parameters, *P* (*θ*) and the prior probability of any infection at each time period *j, P* (*ϕ*_*j*_). This results in the following expression:

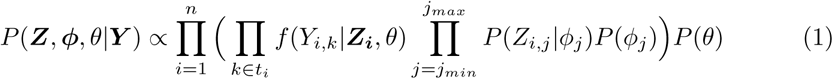

First we outline how this expression is flexibly implemented in the `serosolver` package, then we show how the package can be applied to cross-sectional and longitudinal influenza data from China and Hong Kong to infer key epidemiological and immunological values.

## Design and Implementation

### Antibody process model

For a given individual infection history and set of biological parameters, the antibody process model generates a set of expected log titres for that individual against all possible test strains. Following previous work [27], the expected log titre individual *i* has against the strain that circulated at time *j* when observed at time *k* is defined as a linear combination of the contribution of antibody responses from each prior infection:

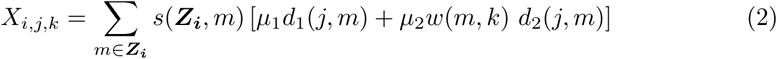

The model components are defined by:

1. Long-term boosting. This is defined by a parameter *µ*_1_, equivalent to the expected persistent rise in titre against a homologous strain following primary infection.
2. Short-term boosting. The transient component of the antibody dynamics is defined by *µ*_2_*w*(*m, k*) = *µ*_2_ max{0, 1 *− ωt*_*m*_}, where *µ*_2_ is the boost in titre, *ω* is a waning parameter to be fitted, and *t*_*m*_ = *k − m* is the time since infection with strain *m*.
3. Long-term cross-reactive antibody response from related strains. We assume the level of cross-reaction between a test strain *j* and infecting strain *m ∈* ***Z***_***i***_ 1 decreases linearly with antigenic distance (see Data section below for definition). The cross-reaction function is *d*_1_(*j, m*) = max{0, 1 *− σ*_1_*δ*_*m,j*_}, where *δ*_*m,j*_ is antigenic distance between strains *j* and *m*, and *σ*_1_ is a fitted parameter.
4. Short-term cross-reactive antibody response. Similar to the long-term response, except this can wane over time. Cross-reactivity between a test strain *j* and infecting strain *m* is defined as *d*_2_(*j, m*) = max{0, 1 *− σ*_2_*δ*_*m,j*_}
5. Antigenic seniority by suppression. This results in lower titres from later infections in comparison to earlier ones. In the model, this works by scaling the titre contribution by a factor *s*(***Z***_***i***_, *m*) = max{0, 1 *− τ* (*N*_*m*_ *−* 1)}, where *N*_*m*_ is the 1 infection number (i.e., primary infection is 1, secondary is 2) and *τ* is a fitted parameter.

The antibody process model can be reduced to simpler models by setting certain parameter values equal to 0. For example, a model without antigenic seniority can be created by setting *τ* = 0 or a model with only waning responses by setting *µ*_1_ = 0.

In addition, `serosolver` can been extended to include more complex antibody kinetics, as described in Supplementary Material 2. We note that the additional immunological phenomena described in Supplementary Material 2 are not exhaustive, and additional mechanisms may easily be implemented by making minor modifications to the package code.

### Antigenic distance

The antibody process model described in Equation 2 includes parameters which describe short- and long-term cross-reactive antibody processes. These processes depend on a metric of antigenic distance between each pair of strains [23]. In the model, the antigenic distance *δ*_*m,j*_ between strains *m* and *j* is therefore defined by a matrix of pairwise distances. `Serosolver` can accommodate antigenically varying strains (all *δ*_*m,j*_ are specified) or a single homologous strain (all *δ*_*m,j*_ = 0). The extent to which strains are antigenically distinct or similar can be described using the distance matrix.

### Observation model

The expected titre *X*_*i,j,k*_ defined in Equation 2 feeds into the observation model, which converts the continuously valued model predicted titre into a discrete observed titre. The distribution of the observed titre consists of a normally distributed random variable *g*(*s*) with mean *X*_*i,j,k*_ and variance *ε*, which is then censored to account for integer-valued log titres in the assay. Hence the probability of observing an empirical titre at time *k* within the limits of a particular assay *Y*_*i,j,k*_ *∈* {0, *…, Y*_*max*_} given expected titre *X*_*i,j,k*_ is,

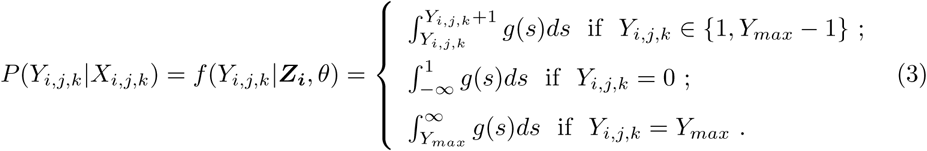

Serosolver includes an additional option to include strain-specific measurement bias, which may arise through strain-specific differences in assay reactivity [26, 30–32]. Specifically, an additional observation error is added to the predicted log antibody titres; this measurement error can be different for each individuals strain or can be specified for a group (or cluster) of strains. The predicted titre 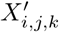 taking into account strain-specific measurement bias is given as:

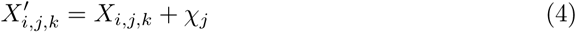

Where *χ*_*j*_ is the measurement offset for strain *j*. The hierarchical form of the measurement bias term may also be specified by the user: *χ*_*j*_ may be estimated as an independent parameter for each *j*; may be assumed to come from a hierarchical distribution 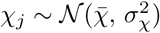; and may be fixed for particular strains/groups e.g., fixing 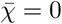 or 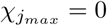.

### Infection history model

`Serosolver` tracks each individual’s infection history as a binary vector of latent states indicating the presence (1) or absence (0) of infection, where each element of the vector represents a time period during which individuals could be infected. The set of infection histories for the sample population are therefore described by a binary matrix, ***Z***, where each row represents an individual, *i*, and each column represents a time, *j*, at which an individual could be infected once. The probability of the infection history matrix, *P* (***Z***) is given by,

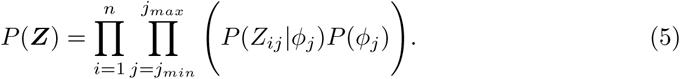

Each infection event (*Z*_*i,j*_) is the outcome of a single Bernoulli trial, with probability 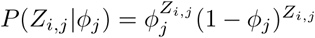. The choice of the prior distribution for the probability of infection, *P* (*ϕ* _*j*_), is discussed below and in further detail in Supplementary Material 1. The time resolution of infection times may be set by the user depending on the data; frequent sampling times affords greater time resolutions (e.g., months), whereas less frequent sampling may be better suited to cruder time resolutions (e.g., years).

The infection history posterior can be used to calculate a key epidemiological measure of interest: the population attack rate over time. Attack rates can be inferred through combining inferred infection histories post-hoc to estimate the proportion of at risk individuals (those that were alive and in the sample) that were infected in a given time period. Summing the columns of the infection history matrix gives the total number of infections for a given time period, whereas summing the rows give the total number of lifetime infections for an individual. To ensure biological plausibility, individual infection histories are constrained to prevent infections before an individual is born and after the last time at which a serum sample was taken. A key feature of the package is that the user is given control over the prior assumptions for the infection history and the probability of infection in each time unit (months, years etc).

### Application to influenza A/H3N2

The initial development of `serosolver` focused on influenza A/H3N2, which has circulated in human populations since 1968 and has undergone substantial antigenic evolution over this time [23, 32–34]. Figure 1 illustrates how our analytical approach applies to influenza A/H3N2. In this case, we make the assumption that the antigenic distance between strains can be described by a two-dimensional distance, with strains moving through the space over time. The expected log antibody titre for a given individual against a specific strain at a specific time can therefore be predicted using this antigenic distance map, the antibody process model described by Equation 2, and the individual level infection histories. Finally, the observed log antibody titres can be used to infer individual level infection histories and antibody process parameters based on time of sampling and the observation model.

**Fig 1.**
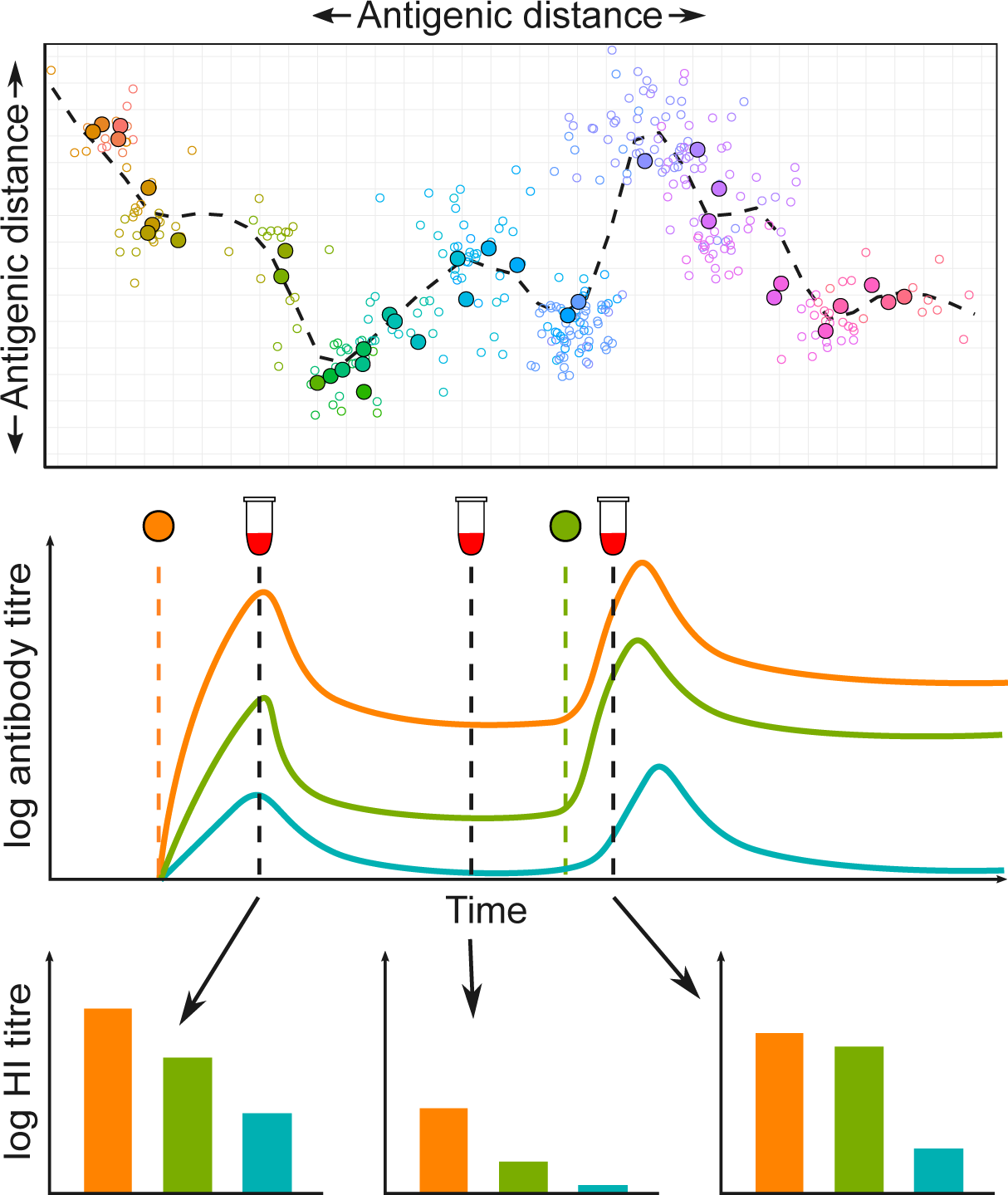
Conceptual overview of the analytical approach used in serosolver, as applied to influenza A/H3N2. Top panel: antigenic map for influenza A/H3N2 using coordinates from [23], with different viruses coloured by year of isolation. Solid points show centroids across all strains isolated in a given year, hollow points show individual strains. Dashed line shows an antigenic summary path, generated by fitting a smoothing spline through the observed isolates. Points further apart in space are less cross-reactive. **Middle panel:** conceptual illustration of the antibody kinetics model. An individual is infected with the orange virus, which results in boosting and waning of homologous antibody titres. In parallel, antibodies that cross react with viruses at different points in antigenic space also boost and wane (green and blue viruses). The individual is later infected by the green virus, which leads to further boosting and waning of antibodies. **Bottom panel:** HI titres measured from serum samples taken at different times capture different parts of the homologous and cross reactive antibody kinetics. Different sampling strategies will represent different subsets of these measurements e.g., a cross-sectional study might inform a single subplot, whereas a longitudinal study might inform just the orange bars from each of the three subplots. Clearly a sampling strategy with multiple serum samples and many viruses tested per sample will provide the most information.

## Data

The `serosolver` package requires two datasets as inputs. The first is an antigenic map, which defines the two-dimensional location of viruses that circulated at each time point during the period of interest, and hence can be used to calculate the pairwise antigenic distance between any two viruses (i.e., *δ*_*m,j*_ in the antibody process model, for strains *m* 1 and *j*). The model automatically sets the potential period during which individuals could have been infected based on the earliest and latest circulating strains in the antigenic map.

The second dataset consists of individual-level log titres against one or more viruses defined in the antigenic map. Each titre measurement is accompanied by a sampling time *k* (i.e., when the serum sample was collected) and strain circulation time *j* (i.e., when the strain was originally isolated).

## Inference

### Prior assumptions

Inference in `serosolver` is fully Bayesian, which means priors must defined for all model parameters and infection histories. The priors on the antibody process parameters are uniform by default, but users may create their own prior function, which may be based on previous analyses. For example, constrained estimates for the short term antibody waning parameters may be used to specify strong beta or Gaussian priors on some of the antibody kinetics parameters for analyses where serum samples may be poorly suited to inform such short term effects.

Priors on the infection histories require more consideration, as the prior also captures any assumptions regarding the infection generating process. Because the number of potential infection times and strains can be vast, the contribution of the infection history prior must be well characterised to avoid any unforeseen bias during inference. The prior assumption on the functional form of *ϕ*, whether individual infection risks are independent at a given time *j*, and whether an individual’s risk of infection depends on infection outcomes at previous times can have important implications for the prior on key infection history summary metrics, such as the attack rate in a given time period and the lifetime number of infections for an individual.

Although the literature for Bayesian variable selection presents a number of potential options, infection states are influenced by epidemiological and immunological structures that are not well characterised by standard prior assumptions (i.e., highly dispersed attack rates and variation in individual-level susceptibility) [35]. We therefore provide the user with flexibility in the assumed infection history and attack rate priors, with different prior assumptions each bringing their own biases and rationale.

`Serosolver` includes four infection history prior options. We summarise these priors in the main text, though an extensive discussion is provided in Supplementary Material 1.

### Hyper-prior on the probability of infection over time, version 1

Under this prior, the probability of infection is given by *ϕ*_*j*_. The infection generating process is:

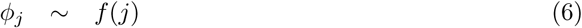

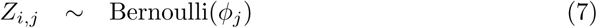

where *f* is a user specified function describing the prior distribution on ***ϕ***, *P* (*ϕ*_*j*_). By default, *f* is the uniform distribution, *ϕ*_*j*_ *∼ unif* (0, 1), though it may be set to incorporate information related to transmission such as seasonality or changes in social behaviour.

### Beta prior on the probability of infection over time, version 2

As in prior 1, this prior assumes that individuals are under a common infection process during a given window of time. However, by placing a beta prior with parameters *α* and *β* and integrating over values for *ϕ*, each *ϕ* need not be estimated explicitly. We have found that this improves convergence of the model fitting framework. The infection generating process is:

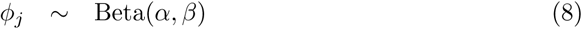

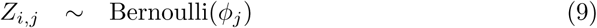

The probability of infection in a given time period is independent of other timperiods, but dependent on the infection status of other individuals in the population at that time. The prior on the per-capita attack rate is therefore a beta distribution, and the prior on the lifetime number of infections for any individual follows a binomial distribution.

### Beta-binomial prior on the total number of infections during an individual’s life, version 3

Unlike priors 1 and 2, this prior assumes that an individual’s risk of infection at a given time is independent of all other individuals. Rather, a prior is placed on the total number of infections that an individual is expected to experience over the course of their life. This is the prior used in our previous work [27]. The infection generating process is assumed to be:

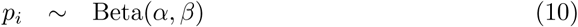

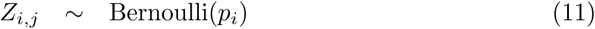

The prior on the per-capita attack rate across all individuals therefore follows binomial distribution, and the prior on the lifetime number of infections for any individual follows a beta-binomial distribution, with parameters *α* and *β* that can be set by the user.

### Beta prior on the probability of any infection, version 4

In the final prior version, infection states are assumed to be independently and identically distributed with respect to both time and individual under the following infection generating process:

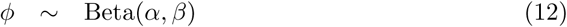

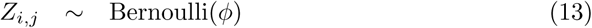

This assumption places a beta-binomial prior on both the number of infections at a given time *j* (the attack rate) and the number of lifetime infections experienced by individual *i*.

### Markov Chain Monte Carlo

`Serosolver` uses a custom, adaptive Markov Chain Monte Carlo (MCMC) framework to sample from the joint posterior distribution of *θ* and ***Z*** conditional on the antibody titre data (Equation 1). The package jointly estimates *θ* and ***Z*** using a Metropolis-Hastings algorithm, alternating between sampling values for *θ* and ***Z***. The MCMC framework automatically tunes the proposal step size for *θ*, and changes the number of individuals sampled for ***Z*** to achieve a specified acceptance rate. Given that MCMC sampling of binary variables is a challenging problem [35, 36], `serosolver` includes additional custom proposal steps included for *Z* to improve chain mixing. The full sampling algorithm for ***Z*** is described in Supplementary Material 1. Briefly, the algorithm uses a random-scan Metropolis-within-Gibbs proposal on infection histories to either propose new infection states or swap the times of existing infection states. These steps were developed to improve MCMC mixing when the infection states in adjacent time periods may be highly correlated. Where automated tuning is insufficient to achieve good mixing, all of the parameters controlling the proposal algorithm are exposed to the user to be changed manually from their default values.

### MCMC diagnostics

To ensure reliable MCMC model fitting, thorough convergence diagnostics must be calculated to ensure that separate MCMC chains have converged on the same distribution, are not trapped in local modes and provide estimates of the posterior distribution with sufficient sample size. `Serosolver` includes functions to test these criteria in two broad categories: (i) visual assessment of convergence and goodness of fit; metrics of convergence checking between-chain agreement, auto-correlation and effective sample size. Alongside existing tools in the `coda` and `bayestools` packages [37, 38], these functions include: MCMC trace and density plots for antibody kinetics parameters; MCMC trace and density plots for inferred attack rates over time; MCMC trace and density plots for inferred infection histories; model predicted titres plotted against observed titres; and inferred attack rates over time. MCMC chain outputs are written to disk during the fitting procedure, and the chain outputs are compatible with the `coda` and `bayesplot` R packages. The full posterior distribution of infection states as augmented data is therefore easily recoverable for further analysis, for example regression analysis of numbers of infections during some period of time.

### Implementation

In `serosolver`, model inputs and assumptions may be changed depending on the serological data and hypotheses under consideration. For example, in some cases the user may be most interested in short-term, fine-scale (e.g., weekly or monthly) dynamics of infection; in other situations, long-term annual dynamics may be of interest. Furthermore, although much of the development of this package came from analysis of influenza A/H3N2 dynamics, these concepts and inputs are easily adaptable to antigenically stable pathogens by specifying the input antigenic map.

The package work flow is divided into a number of distinct stages, which handle the data and parameter inputs, simulation, inference, posterior diagnostics, and analysis (Fig 2) We developed the package to rely on only a few function calls for each of these stages, but with ample room for customisation and flexibility at each stage.

**Fig 2.**
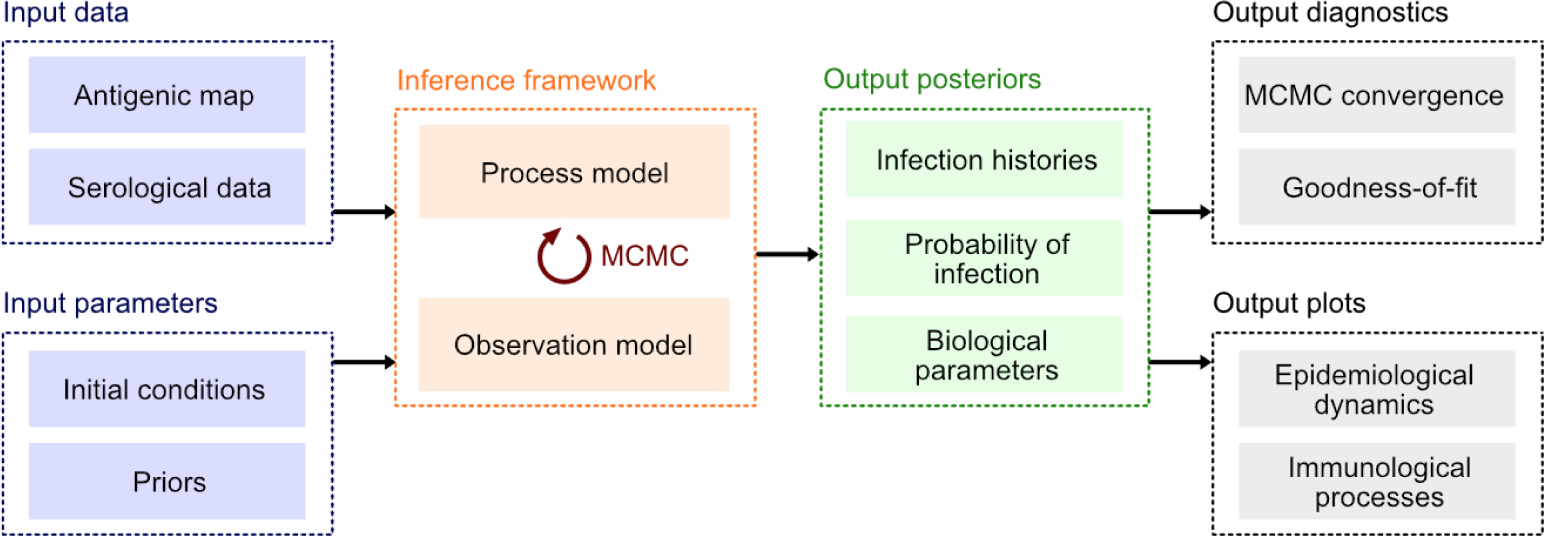
Inputs and outputs for the serosolver R package. The package has two sets of inputs to define data and parameters. These feed into the process model that can either be used to simulate data by itself, or combined with observed data and MCMC to obtain three posterior outputs: individual-level infection histories, population probability of infection, and biological parameters. Once these posteriors have been obtained, `serosolver` can run MCMC diagnostics and plot key immunological and epidemiological processes

To set up the model, users only need to provide: a data frame describing the model parameters (they can also change a flag to fix or estimate any of the parameters); a data frame with the antibody titre data in long format; and an antigenic map describing the antigenic relationship between each strain. Examples of a typical data cleaning workflow are provided in Supplementary Material 4.

`Serosolver` allows users to create their own likelihood and prior functions on top of those provided by default, requiring only that they return a vector of likelihoods (one per individual), and accept arguments for a vector of parameters (matching those defined in the general `serosolver` model) and the infection history matrix. Users can specify which prior assumption about infection histories is used, as specified above. In addition to the range of inbuilt options, the modular workflow of `serosolver` means that custom extensions tailored to particular problems should be readily achievable with only minor modifications to the code. In particular, alternative antibody kinetics models that capture pathogen-specific immunology and alternative assumptions about the infection history generating process.

It is essential to run multiple chains to assess mixing properties and potential bias in any MCMC analysis. Furthermore, model comparison and sensitivity analyses are a common output of model fitting analysis. It is simple to use `serosolver` with a parallel back-end, either through a computing cluster or locally with packages such as `doParallel` [39] to generate multiple chains in parallel. The accompanying vignettes (Supplementary Material 3 and Supplementary Material 4) demonstrate how multiple chains may be run in parallel locally, but we note that much of our own work with `serosolver` is done using a high performance cluster.

## Results

We present two case studies to highlight the range of insights that `serosolver` can generate from serological samples. These cover two types of study designs commonly used to examine epidemiological and immunological dynamics using serological data, which can be thought of as subsets of the observations shown in Fig 1, bottom panel. The first is a serological survey testing individuals against a single homologous strain, which can reveal short-term epidemic dynamics, analogous to observing each of the bars of a single colour from Fig 1. We use data from a longitudinal study conducted in Hong Kong between 2009 and 2011 to estimate short-term antibody kinetics parameters against A/H1N1pdm09 in a population with no prior immunity. The second type of study design involves testing samples against a panel of previously circulating strains, which can provide insights into historical patterns of infection, analogous to observing all of the bars within a single serum sample from Fig 1. To illustrate this application, we apply the package to cross-sectional samples tested against a panel of historical A/H3N2 influenza strains to infer infection histories and antibody kinetics.

### Case Study 1

The first case study uses data from a cohort study in Hong Kong during and after the 2009 A/H1N1pdm09 outbreak [40]. With repeat serological samples tested against a given virus, `serosolver` can reconstruct the unobserved infection dynamics from measured titres collected several months apart. It is also possible to examine these infection dynamics stratified by available demographic variables, such as vaccination status (Fig 3A) and age (Fig 3B). Finally, we can estimate biological parameters shaping the short-term antibody response (Fig 3C).

**Fig 3.**
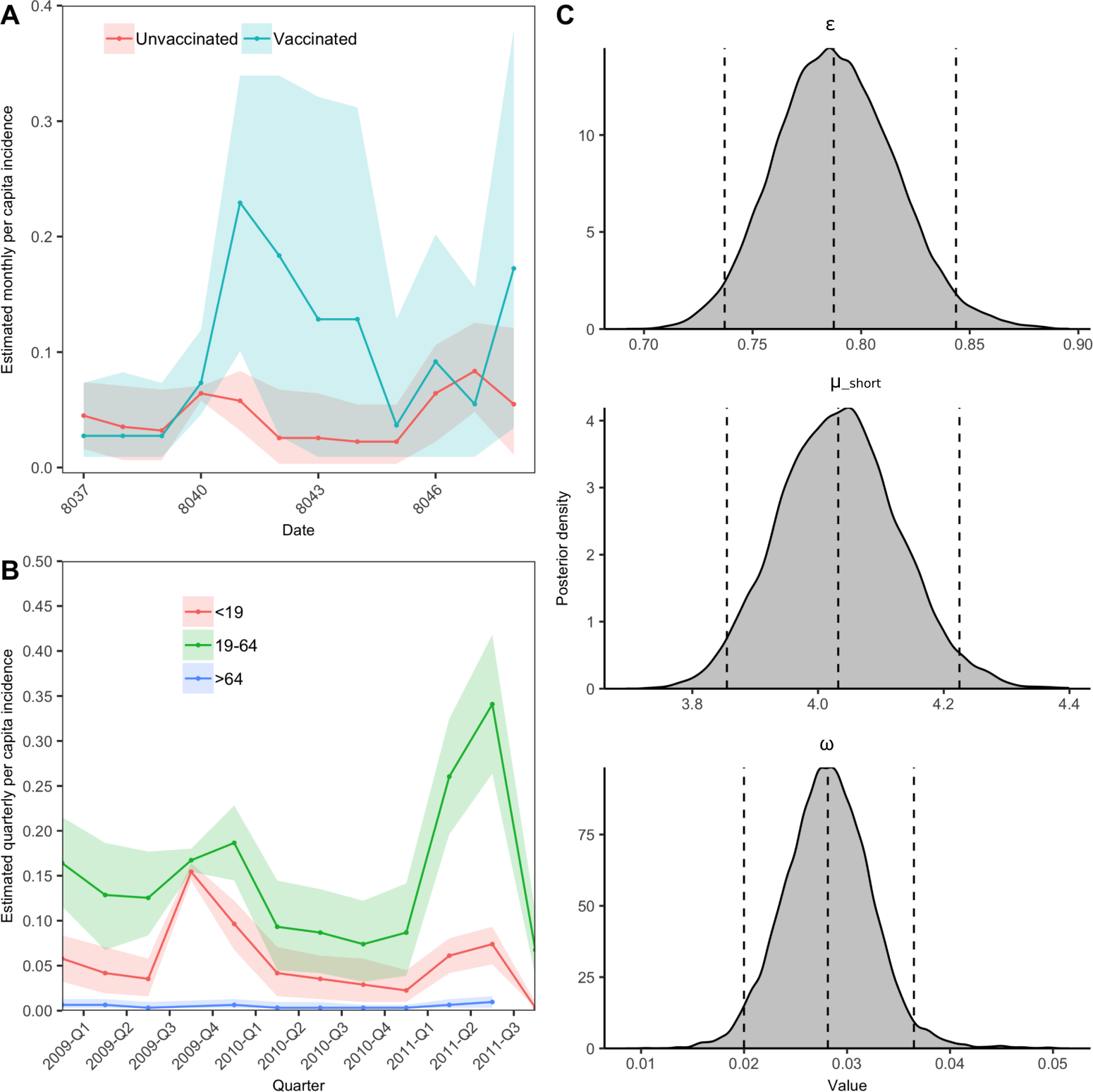
Influenza A/H1N1pdm09 infection dynamics in Hong Kong cohort. A: Exposure rates in unvaccinated individuals. Red line shows median estimate from `serosolver`, with 95% credible intervals (CI); black line shows reported A/H1N1pdm09 isolates. B: Age-specific infection rates in unvaccinated individuals. Lines show median estimates from serosolver for each age group (red: *<*19, green: 19-64, blue: *>*64) with 95% CI. C: Posterior densities of process parameter estimates. Dashed vertical lines represent 2.5th, 50th, and 97.5th percentiles.

We were able to estimate quarterly exposure rates, which could include either infection or vaccination. The inferred peaks in exposure rates are consistent with the observed two waves of the 2009 pandemic. We investigated the impact of vaccination status and age on inferred exposure rates. We found differences in exposure rates in vaccinated individuals compared to unvaccinated individuals, with higher overall exposure rates in vaccinated individuals. Intuitively, we would expect infection rates to 3be lower in vaccinated individuals; however, the converse suggests that vaccination causes boosts in antibody titres that are being inferred as infections. Thus, an individual’s vaccination status is an important consideration when using serological data to infer infection history. Additionally, we observed clear differences in age-stratified exposure rates with exposure rates highest among adults and children, and lowest among the elderly, confirming previous findings of age-stratified exposure rates during the 2009 pandemic [41]. Finally, we aimed to characterise the short-term immune response following infection by estimating short-term antibody kinetics parameters. We found that there is a strong short term boost (16-fold rise) in antibody titre following infection which wanes by 3% every 4 months.

To assess whether data contain enough information to reliably estimate the infection histories and biological process parameters, `serosolver` can be used to run a simulation recovery study. For example, if data of the same structure as the A/H1N1pdm09 outbreak in Hong Kong are generated using plausible parameter values [27], it is possible to re-infer these parameters (Fig 4B) alongside the individual-level infection histories (Fig 4C) and overall probabilities of infection (Fig 4A). However, depending on the sampling frequency, number of tested strains and number of repeat measurements, there are varying levels of information to estimate these quantities. When antibody titre data is sparse, the priors placed on either the antibody parameters, infection histories or probability of infection parameters will have a greater effect on the estimation performance. We therefore recommend routine implementation of simulation recovery on new data to ensure that the most suitable model is being applied to the data available.

**Fig 4.**
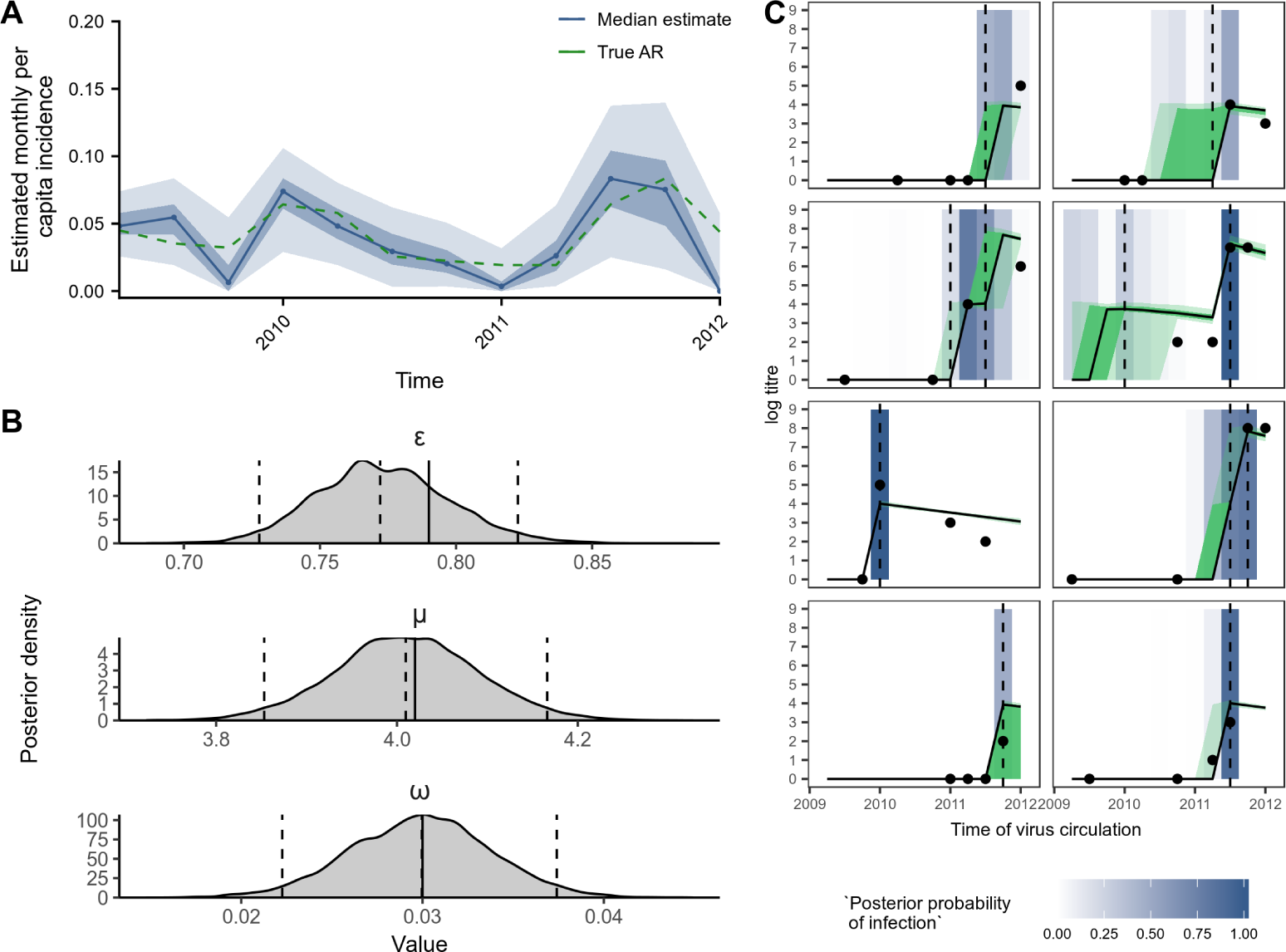
Simulation-recovery of parameter and infection estimates using simulated single strain longitudinal data in same format as the Hong Kong dataset. A: Model estimated attack rates versus ‘true’ attack rates. Solid line shows estimated attack rate with 50% and 95% credible intervals (CI); green dashed line shows true attack rates. B: ‘True’ process parameters used for simulation compared to estimated posterior densities. Black solid vertical lines indicate true parameter values; dashed vertical lines represent 2.5th, 50th, and 97.5th percentiles. C: Model predicted titres and inferred infections compared to observed titres and known infections. Black points indicate observed titres; black lines indicate posterior median model predicted titres; green shading shows 50% and 95% CI on model predicted latent titres; dashed vertical lines indicate the timings of true infections; blue shading indicates posterior probability of infection.

### Case Study 2

The second case study considers cross-sectional serological samples collected in southern China in 2009, which were tested against nine historical influenza A/H3N2 strains that circulated between 1968 and 2008 [29, 42]. `Serosolver` can be used to reconstruct several features of the epidemiological and immunological dynamics. First, Fig 5A shows substantial variation in the inferred historical attack rates of A/H3N2, with clear periods of high incidence interspersed by periods of very low incidence (range of posterior medians: 3.63% to 95.2%). In these analysis, we used a weakly informative prior on the annual attack rate with a mode of 15% with prior version 2. Our posterior estimates were very similar to this, with a median inferred attack rate of 14.6%, suggesting either agreement between the data and prior or a lack of information in the data. We also identified clear age-specific patterns of infection. Fig 5D shows the median number of infections per 10 years alive stratified by age at the time of exposure. These results agree with previous analyses that individuals are infected, or at least experience antibody boosting, less frequently as they get older [27]. Fig 5E shows the proportion of individuals infected at least once by a virus from each of the 14 antigenic clusters considered here stratified by age at the time of exposure. Inference of long-term biological parameters suggested that individuals experience a long-term antibody boost *mu*_1_ of 2.24 log units (posterior median, 95% CI: 1.95-2.51), corresponding to approximately a 4-fold boost to long term homologous titres that wanes with antigenic distance (long term cross reaction *σ*_1_ = 0.105 posterior median, 95% CI: 0.0962-0.113) and decreases with each successive exposure (antigenic seniority parameter, *τ* = 0.0310 posterior median, 95% CI: 0.0210-0.0415).

**Fig 5.**
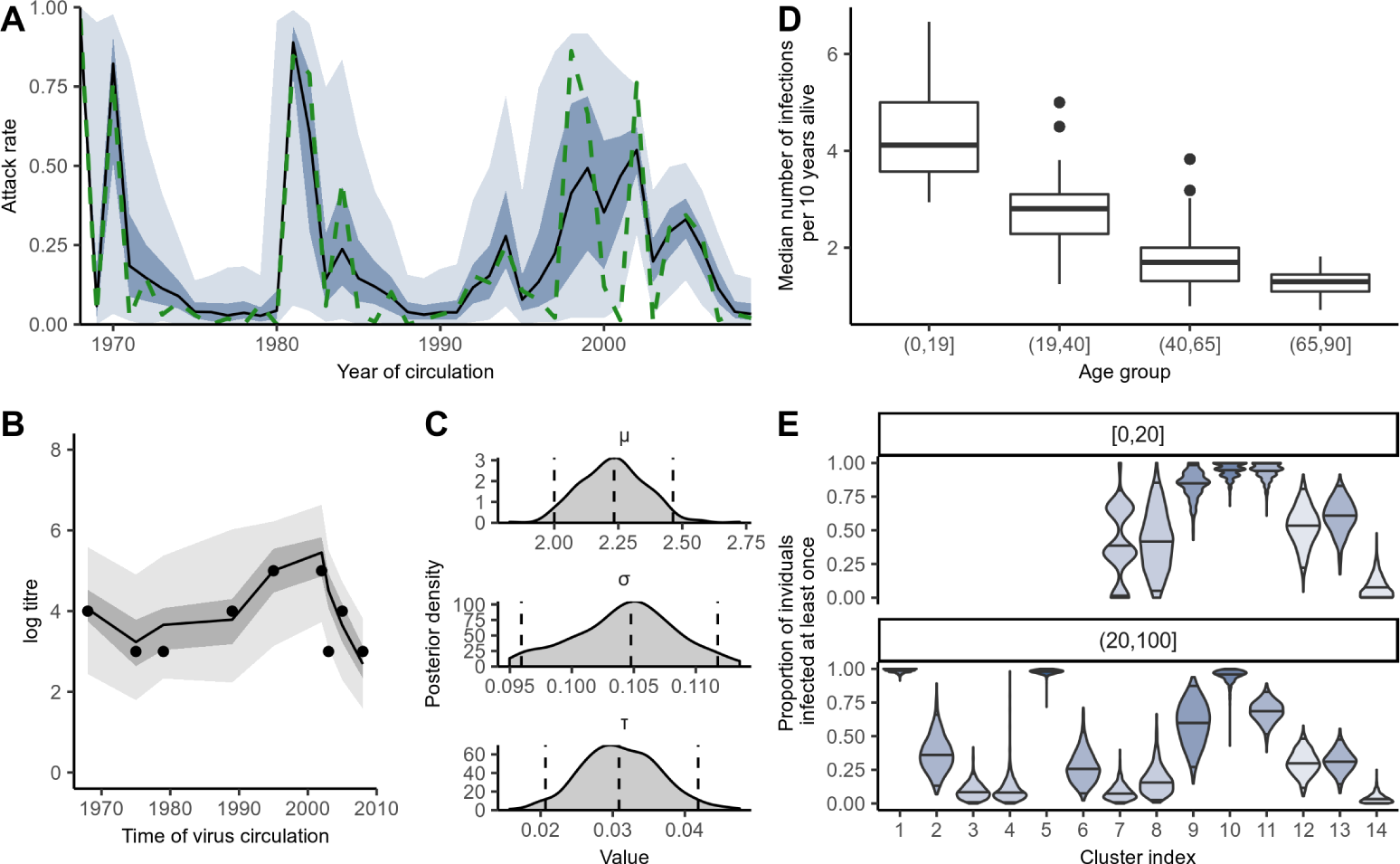
Influenza A/H3N2 dynamics in southern China. A: Inferred historical attack rates. Shaded regions show 50% and 95% credible intervals(CI), black line shows posterior median, dashed green line shows maximum posterior probability estimate; B: Example latent titre trajectory (dark grey region, light grey region and black line show 50% CI, 95% CI and posterior median estimates respectively) against observed titres (black dots) of inferred or one individual. D: Frequency of infection by age group. C: Posterior densities for the inferred antibody kinetics parameters. 95% CI and posterior medians shown as dashed lines. E: Per cluster attack rates in *<*20 and *≥*20 age groups. Clusters with darker shading circulated for longer before succession.

As with the first case study, simulation recovery was used to validate the ability of `serosolver` to correctly infer underlying processes from a given dataset (discussed in 3 detail in Supplementary Material 4).

### Computational performance

`Serosolver` uses a C++ back-end with substantial optimisation to scale the model to large data sets and high infection time resolutions with reasonable run times. Table 1 displays the mean run time of 5 MCMC chains fitting the `serosolver` model to serological data of different dimensions. In the most complex scenario, which involves fitting the model to 164,000 antibody titre measurements and inferring the infection state of 1000 individuals at 164 different time points (164,000 infection states), effective sample sizes *>*200 are achievable for both the antibody process parameters and attack rate estimates in *<*12 hours. For smaller scale analysis (e.g., 100 individuals, *<*5000 titres), high effective sample sizes and well-mixed chains are easily generated within 30 minutes.

**Table 1.**
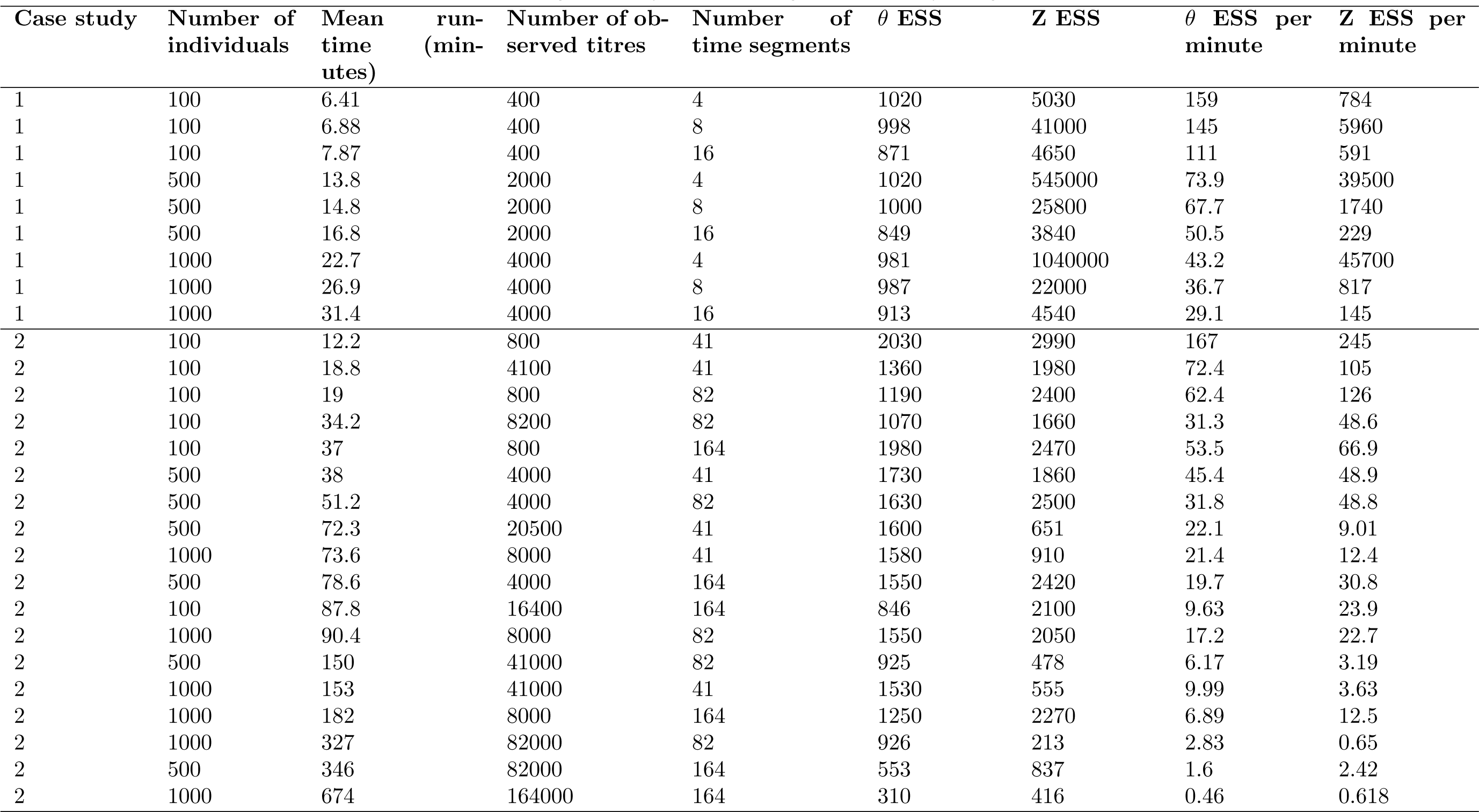
Comparison of run time and posterior sampling efficiency across a range of serosurvey designs.

### Availability and Future Directions

`Serosolver` provides a general inference framework to estimate epidemiological and immunological dynamics from serological data. The open source package is available from GitHub (https://github.com/seroanalytics/serosolver), with detailed accompanying vignettes covering the main implementation and case studies we describe here. The aim of this package is to provide an open source, modifiable framework to fit antibody kinetics models that also require inference of unobserved infections. Disparate serosurveys measuring antibody titres over time are often underpinned by comparable dynamics, and we therefore felt that a unifying tool to enable quick reproduction and direct comparison of analyses across different datasets would be a useful addition to the literature.

As well as the stand-alone applications we have illustrated in the case studies above, `serosolver` could easily link with traditional epidemiological analysis. The results presented here are not intended to be exhaustive analyses, but rather to demonstrate the utility and range of insights that can be generated from serological data. In particular, the posterior latent individual-level infection histories and titre trajectories could act as inputs into regression models. For example, `serosolver` outputs could be combined with syndromic or lab-confirmation data to examine the relationship between susceptibility and titre at time of infection [43]. These methods could also apply to other pathogens; a similar model structure has recently been used to examine latent titres for dengue [44].

Moreover, `serosolver` can incorporate prior knowledge on time of exposure either from surveillance data or, if relevant, temporal climate variables. In the case studies presented, we used relatively simple priors for the probability of infection. However, more complex temporal priors could be imposed by having a different prior distribution for the probability of at each time point (i.e., different value of *α* and *β*) to account for seasonality in transmission dynamics. In the future, we hope to extend `serosolver` to include non-linear feedback between past exposures and future risk, by embedding an epidemic model as well as the probability of infection [45]. In theory, this package could be used to generate an ongoing database of inferred immunological parameters, allowing estimates to be updated and combined between to better estimate attack rates and infection histories in less data-rich cohorts.

`Serosolver` could also be used to inform the design of serological sample collection and testing. Given potential logistical or budgetary restrictions on analysis of stored sera or collection of new samples, `serosolver` could be used to simulate different study designs and show how accurately these designs could recover the main parameters of interest.

At present, `serosolver` focuses on inference for a single exposure type. However, for viruses like influenza and dengue, individuals may be exposed to multiple subtypes or serotypes in the same season. Exposure to one antigen may cross react with another antigen providing protection against antigens an individual has not been directly exposed to. For example, infection with influenza A/H1N1 may provide cross-reactive protection against other group 1 viruses, and A/H3N2 against group 2 viruses [46]. Additionally, the incorporation of multiple exposures can facilitate the inclusion of vaccine exposure. In influenza, where vaccination is recommended annually, exposure to vaccination is an important piece of the immunological life course puzzle of an individual [47]. In its current form, `serosolver` can estimate differences between exposures by being fit independently to different subtypes. It can also fit models separately to vaccinated or unvaccinated populations to estimate how serological dynamics vary between these groups. Although this is a useful first approximation, future versions of `serosolver` will include potential for multiple exposure types during the same season so that any interactions can be modelled explicitly.

There is increasing evidence that serological titre data contain substantial additional information about infection and immunity dynamics, which are not captured by simple four-fold rise metrics [14, 44, 48, 49] Furthermore, in multi-strain pathogen systems, evidence is mounting that individual-level heterogeneity in unobserved exposure histories is a key driver of susceptibility to infection and disease [26, 47, 50, 51].

`Serosolver` provides a generic framework to extract this information from commonly collected data. As serological data become increasingly available, it will be important to develop modern analytical methods and tools that account for known biological and epidemiological processes that may confound or bias inference [49, 52–54].

## Supporting information

Supplementary Material 1. Full description and discussion of the infection history priors and their implications for inference.

Supplementary Material 2. Additional immunological mechanisms and how to modify code to incorporate alternative antibody kinetics.

Supplementary Material 3. Case study 1 vignette with all code required for model fitting, figure generation and simulation recovery.

Supplementary Material 4. Case study 2 vignette with all code required for model fitting, figure generation and simulation recovery.

## References

1. Hirst GK. The quantitative determination of influenza virus and antibodies by means of red cell agglutination. J Exp Med. 1942;75(1):49–64.

2. Laurie KL, Engelhardt OG, Wood J, Heath A, Katz JM, Peiris M, et al. An international laboratory comparison of influenza microneutralisation assay protocols for A (H1N1) pdm09, A (H3N2) and A (H5N1) influenza A viruses by CONSISE. Clin Vaccine Immunol. 2015; p. CVI–00278.

3. Cutts FT, Hanson M. Seroepidemiology: an underused tool for designing and monitoring vaccination programmes in low- and middle-income countries. Tropical Medicine & International Health. 2016;21(9):1086–1098. doi:10.1111/tmi.12737

4. Amanna IJ, Carlson NE, Slifka MK. Duration of humoral immunity to common viral and vaccine antigens. The New England journal of medicine. 2007;357(19):1903–1915. doi:10.1542/peds.2008-2139LLLL

5. Salje H, Cauchemez S, Alera MT, Rodriguez-Barraquer I, Thaisomboonsuk B, Srikiatkhachorn A, et al. Reconstruction of 60 Years of Chikungunya Epidemiology in the Philippines Demonstrates Episodic and Focal Transmission. Journal of Infectious Diseases. 2016;213(4):604–610. doi:10.1093/infdis/jiv470

6. Cardoso CWC, Kikuti M, Prates AAPPB, Paploski IIAD, Tauro LBL, Silva MMO, et al. Unrecognized Emergence of Chikungunya Virus during a Zika Virus Outbreak in Salvador, Brazil. PLOS Neglected Tropical Diseases. 2017;11(1):e0005334. doi:10.1371/journal.pntd.0005334

7. Hobson D, Curry RL, Beare AS, Ward-Gardner A. The role of serum haemagglutination-inhibiting antibody in protection against challenge infection with influenza A2 and B viruses. J Hyg. 1972;70(4):767–777.

8. Coudeville L, Bailleux F, Riche B, Megas F, Andre P, Ecochard R. Relationship between haemagglutination-inhibiting antibody titres and clinical protection against influenza: development and application of a bayesian random-effects model. BMC medical research methodology. 2010;10:18. doi:10.1186/1471-2288-10-18

9. Wu JT, Ho A, Ma ESK, Lee CK, Chu DKW, Ho PL, et al. Estimating Infection Attack Rates and Severity in Real Time during an Influenza Pandemic: Analysis of Serial Cross-Sectional Serologic Surveillance Data. PLoS Medicine. 2011;8(10):e1001103. doi:10.1371/journal.pmed.1001103

10. Wu JT, Leung K, Perera RAPM, Chu DKW, Lee CK, Hung IFN, et al. Inferring Influenza Infection Attack Rate from Seroprevalence Data. PLoS Pathogens. 2014;10(4):e1004054. doi:10.1371/journal.ppat.1004054

11. Zhao X, Siegel K, Chen MIC, Cook AR. Rethinking thresholds for serological evidence of influenza virus infection. Influenza and Other Respiratory Viruses. 2017;11(3):202–210. doi:10.1111/irv.12452

12. Katz JM, Hancock K, Xu X. Serologic assays for influenza surveillance, diagnosis and vaccine evaluation. Expert Review of Anti-infective Therapy. 2011;9(6):669–683. doi:10.1586/eri.11.51

13. Wood JM, Gaines-Das RE, Taylor J, Chakraverty P. Comparison of influenza serological techniques by international collaborative study. Vaccine. 1994;12(2):167–74.

14. Cauchemez S, Horby P, Fox A, Mai LQ, Thanh LT, Thai PQ, et al. Influenza infection rates, measurement errors and the interpretation of paired serology. PLoS Pathog. 2012;8(12):e1003061.

15. Miller E, Hoschler K, Hardelid P, Stanford E, Andrews N, Zambon M. Incidence of 2009 pandemic influenza A H1N1 infection in England: a cross-sectional serological study. Lancet. 2010;375(9720):1100–1108.doi:10.1016/S0140-6736(09)62126-7

16. Freeman G, Pereram RAPM, Ngan E, Fang VJ, Cauchemez S, Ip DKM, et al. Quantifying homologous and heterologous antibody titre rises after influenza virus infection. Epidemiology and Infection. 2016;144(11):2306–2316. doi:10.1017/S0950268816000583

17. Barr IG, Russell C, Besselaar TG, Cox NJ, Daniels RS, Donis R, et al. WHO recommendations for the viruses used in the 2013â€“2014 Northern Hemisphere influenza vaccine: Epidemiology, antigenic and genetic characteristics of influenza A(H1N1)pdm09, A(H3N2) and B influenza viruses collected from October 2012 to January 2013. Vaccine. 2014;32(37):4713–4725. doi:10.1016/J.VACCINE.2014.02.014

18. Huang QS, Bandaranayake D, Wood T, Newbern EC, Seeds R, Ralston J, et al. Risk Factors and Attack Rates of Seasonal Influenza Infection: Results of the Southern Hemisphere Influenza and Vaccine Effectiveness Research and Surveillance (SHIVERS) Seroepidemiologic Cohort Study. The Journal of Infectious Diseases. 2019;219(3):347–357. doi:10.1093/infdis/jiy443

19. Katzelnick L, Fonville J, Gromowski G, Arriaga J, Green A, et al. Dengue viruses cluster antigenically but not as discrete serotypes. Science. 2015;349(6254):1338–1343.

20. Priyamvada L, Quicke KM, Hudson WH, Onlamoon N, Sewatanon J, Edupuganti S, et al. Human antibody responses after dengue virus infection are highly cross-reactive to Zika virus. Proceedings of the National Academy of Sciences. 2016;113(28):7852–7857. doi:10.1073/pnas.1607931113

21. Montoya M, Collins M, Dejnirattisai W, Katzelnick LC, Puerta-Guardo H, Jadi R, et al. Longitudinal Analysis of Antibody Cross-neutralization Following Zika Virus and Dengue Virus Infection in Asia and the Americas. The Journal of Infectious Diseases. 2018;218(4):536–545. doi:10.1093/infdis/jiy164

22. Nachbagauer R, Choi A, Hirsh A, Margine I, Iida S, Barrera A, et al. Defining the antibody cross-reactome directed against the influenza virus surface glycoproteins. Nature Immunology. 2017;18(4):464–473. doi:10.1038/ni.3684

23. Smith DJ, Lapedes AS, de Jong JC, Bestebroer TM, Rimmelzwaan GF, Osterhaus ADME, et al. Mapping the antigenic and genetic evolution of influenza virus. Science. 2004;305(5682):371–376.

24. Lapedes A, Farber R. The Geometry of Shape Space: Application to Influenza. Journal of Theoretical Biology. 2001;212(1):57–69. doi:10.1006/jtbi.2001.2347

25. Cai Z, Zhang T, Wan XF, Dhillon I, Butler E. A Computational Framework for Influenza Antigenic Cartography. PLoS Computational Biology. 2010;6(10):e1000949. doi:10.1371/journal.pcbi.1000949

26. Fonville JM, Wilks SH, James SL, Fox A, Ventresca M, Aban M, et al. Antibody landscapes after influenza virus infection or vaccination. Science. 2014;346(6212):996–1000.

27. Kucharski AJ, Lessler J, Cummings DAT, Riley S. Timescales of influenza A/H3N2 antibody dynamics. PLoS Biol. 2018;16(8):e2004974.

28. Yuan HY, Baguelin M, Kwok KO, Arinaminpathy N, van Leeuwen E, Riley S. The impact of stratified immunity on the transmission dynamics of influenza. Epidemics. 2017;.

29. Kucharski AJ, Lessler J, Read JM, Zhu H, Jiang CQ, Guan Y, et al. Estimating the life course of influenza A(H3N2) antibody responses from cross-sectional data. PLoS Biol. 2015;13(3):e1002082.

30. Hensley SE, Das SR, Bailey AL, Schmidt LM, Hickman HD, Jayaraman A, et al. Hemagglutinin receptor binding avidity drives influenza A virus antigenic drift. Science (New York, NY). 2009;326(5953):734–6. doi:10.1126/science.1178258

31. Li Y, Bostick DL, Sullivan CB, Myers JL, Griesemer SB, Stgeorge K, et al. Single hemagglutinin mutations that alter both antigenicity and receptor binding avidity influence influenza virus antigenic clustering. Journal of virology. 2013;87(17):9904–10. doi:10.1128/JVI.01023-13

32. Bedford T, Suchard MA, Lemey P, Dudas G, Gregory V, Hay AJ, et al. Integrating influenza antigenic dynamics with molecular evolution. eLife. 2014;2014(3):e01914. doi:10.7554/eLife.01914

33. Fitch WM, Bush RM, Bender CA, Cox NJ. Long term trends in the evolution of H(3) HA1 human influenza type A. Proceedings of the National Academy of Sciences of the United States of America. 1997;94(15):7712–8. doi:10.1073/PNAS.94.15.7712

34. Russell CA, Jones TC, Barr IG, Cox NJ, Garten RJ, Gregory V, et al. The Global Circulation of Seasonal Influenza A (H3N2) Viruses. Science. 2008;320(5874).

35. O’Hara RB, Sillanpää MJ. A review of Bayesian variable selection methods: what, how and which. Bayesian Analysis. 2009;4(1):85–117. doi:10.1214/09-BA403

36. George EI, McCulloch RE. Approaches for Bayesian Variable Selection. Statistica Sinica. 1997;7:339–373. doi:10.2307/24306083

37. Plummer M, Best N, Cowles K, Vines K. CODA: Convergence Diagnosis and Output Analysis for MCMC. R News. 2006;6(1):7–11.

38. Gabry J. bayesplot: Plotting for Bayesian models; 2017. Available from: http://mc-stan.org/.

39. Foreach Parallel Adaptor for the ‘parallel’ Package [R package doParallel version 1.0.14];.

40. Kwok KO, Riley S, Perera RAPM, Wei VWI, Wu P, Wei L, et al. Relative incidence and individual-level severity of seasonal influenza A H3N2 compared with 2009 pandemic H1N1. BMC Infectious Diseases. 2017;17(1):337. doi:10.1186/s12879-017-2432-7

41. Riley S, Kwok KO, Wu KM, Ning DY, Cowling BJ, Wu JT, et al. Epidemiological characteristics of 2009 (H1N1) pandemic influenza based on paired sera from a longitudinal community cohort study. PLOS Medicine. 2011;8(6):e1000442. doi:10.1371/journal.pmed.1000442

42. Lessler J, Riley S, Read JM, Wang S, Zhu H, Smith GJD, et al. Evidence for antigenic seniority in influenza A (H3N2) antibody responses in southern China. PLoS Pathogens. 2012;8(7):e1002802. doi:10.1371/journal.ppat.1002802

43. Lau MSY, Cowling BJ, Cook AR, Riley S. Inferring influenza dynamics and control in households. Proc Natl Acad Sci U S A. 2015;112(29):9094–9099.

44. Salje H, Cummings DA, Rodriguez-Barraquer I, Katzelnick LC, Lessler J, Klungthong C, et al. Reconstruction of antibody dynamics and infection histories to evaluate dengue risk. Nature. 2018;557(7707):719.

45. Ranjeva S, Subramanian R, Fang VJ, Leung GM, Ip DKM, Perera RAPM, et al. Age-specific differences in the dynamics of protective immunity to influenza. Nature Communications. 2019;10(1):1660. doi:10.1038/s41467-019-09652-6

46. Gostic KM, Ambrose M, Worobey M, Lloyd-Smith JO. Potent protection against H5N1 and H7N9 influenza via childhood hemagglutinin imprinting. Science. 2016;354(6313):722–726.

47. Cobey S, Hensley SE. Immune history and influenza virus susceptibility. Current opinion in virology. 2017;22:105–111.

48. Pepin KM, Kay SL, Golas BD, Shriner SS, Gilbert AT, Miller RS, et al. Inferring infection hazard in wildlife populations by linking data across individual and population scales. Ecology Letters. 2017;doi:10.1111/ele.12732

49. White MT, Griffin JT, Akpogheneta O, Conway DJ, Koram KA, Riley EM, et al. Dynamics of the antibody response to Plasmodium falciparum infection in African children. J Infect Dis. 2014;210(7):1115–1122.

50. Guthmiller JJ, Wilson PC. Harnessing immune history to combat influenza viruses. Current Opinion in Immunology. 2018;53:187–195. doi:10.1016/J.COI.2018.05.010

51. Andrews SF, Huang Y, Kaur K, Popova LI, Ho IY, Pauli NT, et al. Immune history profoundly affects broadly protective B cell responses to influenza. Sci Transl Med. 2015;7(316):316ra192–316ra192. doi:10.1126/scitranslmed.aad0522

52. Cauchemez S, Hoze N, Cousien A, Nikolay B, Ten bosch Q. How Modelling Can Enhance the Analysis of Imperfect Epidemic Data. Trends in Parasitology. 2019;35(5):369–379. doi:10.1016/J.PT.2019.01.009

53. de Lusignan S, Borrow R, Tripathy M, Linley E, Zambon M, Hoschler K, et al. Serological surveillance of influenza in an English sentinel network: pilot study protocol. BMJ open. 2019;9(3):e024285. doi:10.1136/bmjopen-2018-024285

54. Metcalf CJE, Farrar J, Cutts FT, Basta NE, Graham AL, Lessler J, et al. Use of serological surveys to generate key insights into the changing global landscape of infectious disease. The Lancet. 2016;388(10045):728–730. doi:10.1016/S0140-6736(16)30164-7

