## Supplementary Material 1 for "Serosolver: an open source tool to infer epidemiological and immunological dynamics from serological data"

Supplementary Material 1: Infection History Prior

#### 1 Motivation

Here we provide more details about the assumptions and priors required to perform Bayesian inference of infection histories with a model that considers past infection outcomes of a population as a matrix of latent features. It also serves as a reference guide to understand the impact of the different priors on inferred infection histories and attack rates in the accompanying `serosolver` package.

Each infection event, $Z_{i,j}$, is the outcome of a single Bernoulli trial, where $Z_{i,j} = 1$ indicates that individual $i$ was infected at time $j$, and $Z_{i,j} = 0$ indicates that they were not. Infection events are not observed directly. Rather, infections lead to the production of antibodies that undergo longitudinal and cross-reactive kinetics. If we consider a system with $n$ individuals who may be infected once in each of $m$ distinct time periods, then there are $nm$ possible infection events. We are interested in jointly inferring the posterior distributions outcomes of each of these $nm$ infection events and a set of antibody kinetics parameters using serological data. That is, what is the posterior probability of the $nm$ infection outcomes in $Z$ given the serological data $y$, defined as:

$$P(\boldsymbol{Z}, \theta | \boldsymbol{Y}) \propto \prod_{i=1}^{n} \Big( \prod_{k \in t_i} f(Y_{i,k} | \boldsymbol{Z_i}, \theta) \prod_{j=j_{min}}^{j_{max}} P(Z_{i,j}) \Big) P(\theta) \tag{1}$$

where $\theta$ is the vector of antibody kinetics parameters that describe the link between $\boldsymbol{Z}$ and $\boldsymbol{Y}$. $P(\boldsymbol{Z})$ and $P(\theta)$ are independent. Crucially, only $\boldsymbol{Y}$ is known, so we must infer (or augment) the values of $\boldsymbol{Z}$ as latent features. The function $f(\boldsymbol{Y_i} | \boldsymbol{Z_i}, \theta)$ that relates infection histories to the likelihood of observed antibody titres is complex due to unobserved immunological phenomena, distinguishing the model from more standard Bayesian regression models (ie. we are not interested in inferring a regression coefficient $\beta_{i,j}$ for each $Z_{i,j}$). $f(\boldsymbol{Y_i} | \boldsymbol{Z_i}, \theta)$ is the antibody kinetics model and is described elsewhere and $P(\theta)$ represents a standard Bayesian prior. However, the choice of structure for $P(\boldsymbol{Z})$ is not immediately obvious in the context of infection events.

This problem falls within the remit of Bayesian binary variable selection: a well described area of research in the context of regression models and a notoriously challenging problem as the number of binary variables and resulting model space grows large [1, 2, 3]. Methods such as Stochastic search variable selection and Reversible Jump Markov chain Monte Carlo are well established for model selection tasks, but not sufficient to describe the present problem, where binary outcomes are the result of complex, unobserved epidemiological processes that must be taken into consideration. In particular, we must carefully consider that not all infection events are independent. For example, individuals that are infectious at a given time exert a force of infection on other individuals in the same population, and *a priori* we do not know if an individual experienced many or few infections over their lifetime. Here, we describe a number of priors that can take these processes into account and discuss their implications on inferring individual infection histories and attack rates with the `serosolver` package.

#### 2 Intuitive prior

Intuitively, an uninformative prior on an indicator random variable, $Z_{i,j}$, might be that $Z_{i,j} = 1$ occurs with fixed $p = 0.5$ and $Z_{i,j} = 0$ occurs with fixed $q = (1 - p) = 0.5$. More generally:

$$P(Z_{i,j}) = p_{i,j}^{Z_{i,j}}(1 - p_{i,j})^{1-Z_{i,j}} \tag{2}$$

$$P(\boldsymbol{Z_i}) = \prod_{j=1}^{m} p_{i,j}^{Z_{i,j}}(1 - p_{i,j})^{1-Z_{i,j}} \tag{3}$$

$$P(\boldsymbol{Z}) = \prod_{i=1}^{n}\prod_{j=1}^{m} p_{i,j}^{Z_{i,j}}(1 - p_{i,j})^{1-Z_{i,j}} \tag{4}$$

If we consider an individual's infection history $Z_i$ to be a sequence of binary variables, $\boldsymbol{Z_i} = [Z_{i,1}, Z_{i,2}, \ldots, Z_{i,m}])$, then this prior implicitly assumes that the total number of infections are binomially distributed with mean $m$, where $p = 0.5$ (with the binomial coefficient $\binom{m}{k}$, as there are more combinations of infections that lead to $k \approx mp$ than $k = 0$ and $k = m$). Similarly, the total number of infections per individual $i$ and per time $j$ are both also binomially distributed. Setting $p = 0.5$ is equivalent to assuming that all infection histories are equally likely: an infection history with all infections is as likely as one with no infections, and as likely as any other sequence of 1s and 0s.

Although intuitive, this prior structure also places a strong prior on the total number of infections where $pm$ infections are more likely than an infection history with 0 or $m$ total infections. In the situation where there is little contribution from the data ($P(\boldsymbol{Y}|\boldsymbol{Z}, \theta)$ contributes little) to the posterior, $P(\boldsymbol{Z})$ would bias the inferred infection histories towards $pm$ infections per individual and $pn$ infections per unit time. The posterior probability of an infection history with few infections and large amounts of antibody boosting per infection would be far lower than of an infection history with $0.5m$ infections and low antibody boosting.

In reality, influenza infections likely happen less frequently than every other year, though do occur infrequently for some individuals and frequently for others. Similarly, although the total number of infections in a given influenza season are well described by a binomial distribution, the distribution of infections across multiple outbreaks is likely over-dispersed relative to the binomial distribution due to between-outbreak variation in severity. Clearly a prior that allows us to capture these assumptions is desirable.

### 3 `serosolver` priors

`Serosolver` provides 4 options for different infection probability priors. Each option follows a different set of assumptions and definitions, leading to different implications for infection history inference. Table 1 summarises each of these priors and the situations when each one is advised, and the remainder of this section describes their derivations in more detail. The implementation in `serosolver` for each of these priors is described in the Appendix.

#### 3.1 Hyper-prior on per-time infection probability

Individuals may be from the same population, which implies that their risks of infection are correlated. Under this prior we assume that individuals are infected by the same infection process during a given time period (ie. there is a force of infection on the population), and that infection processes are independent across time periods. There is an intuition to this approach which is revealed by the following question: given a sample of $n$ individuals for which we know all $n$ infection states for time $j$, what is the prior predictive probability that individual $n+1$ was also infected during time $j$? If we know that 80% of the population were infected at time $j$, then we should have some prior belief that individual $n+1$ was also infected.

Under this prior, the probability of success (infection) is given as $p = \phi_j$ and probability of failure (no infection) is given as $q = 1 - \phi_j$. $\phi$ is related to the attack rate and therefore the probability of an individual becoming infected. Under this prior, the infection generating process is:

$$\phi_j \sim f(j) \tag{5}$$

$$Z_{i,j} \sim \text{Bernoulli}(\phi_j) \tag{6}$$

Where $f$ is a user specified function describing the prior distribution of $\phi$. The probability mass function for an individual infection event in time period $j$ is therefore given by:

$$P(Z_{i,j}|\phi_j) = \phi_j^{Z_{i,j}}(1-\phi_j)^{(1-Z_{i,j})} \tag{7}$$

Thus, the likelihood of observing a particular combination of infections at time $j$ is given by a Bernoulli trials process:

$$\begin{aligned} P([Z_{1,j}, Z_{2,j}, \ldots, Z_{n,j}]|\phi_j) &= \textstyle\prod_{i=1}^n \phi_j^{Z_{i,j}}(1-\phi_j)^{(1-Z_{i,j})} \\ &= \phi_j^{k_j}(1-\phi_j)^{(m_j-k_j)} \end{aligned}$$

Retaining the correlation between individuals and adding $m$ infection times to the system, the likelihood of $Z$ conditional on $\phi$ becomes:

$$P(\boldsymbol{Z}|[\phi_1, \phi_2, \ldots, \phi_m]) = \prod_{j=1}^{m} \phi_j^{k_j}(1-\phi_j)^{(n_j-k_j)}$$

Where $\boldsymbol{Z}$ is an $n$ by $m$ matrix representing the outcome of $m$ possible infection events for $n$ individuals. The first example in Section 2 above makes the strong assumption that $\phi_j$ is fixed at 0.5 for all times $j$. As discussed above, this implicitly assumes that the total number of infections in a given time period and over an individual's life are both binomially distributed with means $pn_j$ and $pa_i$, where $n$ is the total number of individuals alive during time period $j$ and $a_i$ is the number of years that individual $i$ is alive.

To avoid this strong assumption, we can assume that all $\phi$ are unknown parameters to be estimated by correctly defining a prior on $\phi$. Under Bayes theorem and using a beta prior (the conjugate prior for the Bernoulli process) on $\phi$, we derive the following:

$$\begin{aligned}P(\boldsymbol{\phi}|\boldsymbol{Z})P(\boldsymbol{Z}) &= P(\boldsymbol{Z}|\boldsymbol{\phi})P(\boldsymbol{\phi}) \\ &= \prod_j \phi_j^{k_j}(1-\phi_j)^{(m-k_j)}\frac{\phi_j^{\alpha-1}(1-\phi_j)^{\beta-1}}{B(\alpha,\beta)} \\ &= \prod_j \phi_j^{\alpha+k_j-1}(1-\phi_j)^{\beta+m-k_j-1}\frac{1}{B(\alpha,\beta)}\end{aligned}$$

With appropriate choice of values for $\alpha$ and $\beta$ (either $\alpha = \beta = 1$ for uniform, $\alpha = \beta = 0.5$ for Jeffrey's prior and $\alpha = \beta = \frac{1}{3}$ for a neutral prior) [4], we recover the posterior distribution for $\phi$ conditional on the infection history matrix $Z$. In `serosolver`, $P(\boldsymbol{\phi})$ may be any proper prior distribution and need not be the Beta distribution.

Under this prior Equation 1 is modified to give:

$$\begin{aligned}P(\boldsymbol{Z},\theta,\boldsymbol{\phi}|\boldsymbol{Y}) &= \frac{P(\boldsymbol{Z},\boldsymbol{\phi},\theta,\boldsymbol{Y})}{P(\boldsymbol{Y})} && (8)\\ &= \frac{P(\boldsymbol{Y}|\boldsymbol{Z},\boldsymbol{\phi},\theta)P(\boldsymbol{Z},\boldsymbol{\phi},\theta)}{P(\boldsymbol{Y})} && (9)\\ &= \frac{P(\boldsymbol{Y}|\boldsymbol{Z},\boldsymbol{\phi},\theta)P(\boldsymbol{Z}|\boldsymbol{\phi},\theta)P(\boldsymbol{\phi},\theta)}{P(\boldsymbol{Y})} && (10)\\ &= \frac{P(\boldsymbol{Y}|\boldsymbol{Z},\phi)P(\boldsymbol{Z}|\boldsymbol{\phi})P(\boldsymbol{\phi})P(\theta)}{P(\boldsymbol{Y})} && (11)\end{aligned}$$

$$P(\boldsymbol{Z},\boldsymbol{\phi},\theta|\boldsymbol{Y}) \propto \prod_{i=1}^{n}\Big(\prod_{k\in t_i} f(Y_{i,k}|\boldsymbol{Z_i},\theta)\prod_{j=j_{min}}^{j_{max}} P(Z_{i,j}|\phi_j)P(\phi_j)\Big)P(\theta) \quad (12)$$

This structure opens up a number of useful possibilities. For example: $\phi$ may be defined as a function rather than a variable; different priors may be placed on different times $j$; $\phi$ may be inferred explicitly.

##### 3.1.1 Prior on number of lifetime infections

Assuming that all $P(\phi_j)$ are equal, the total number of lifetime infections for an individual follows a binomial distribution with $p = \mathbb{E}(\phi)$. Using a beta prior for $P(\phi)$ with parameters $\alpha$ and $\beta$ and assuming that $\alpha$ and $\beta$ are equal for all $j$, then $P(\sum_j Z_{i,j} = Z_{i,1} + Z_{i,2} + \cdots + Z_{i,m})$ follows a binomial distribution with mean $\frac{\alpha}{\alpha+\beta}$ and $N = m_i$, where $m_i$ is the number of time periods that individual $i$ could be infected. Although a binomial prior is relatively informative, it is also intuitive: if the expectation of the attack rates for all times $j$ is $p = 0.5$, then we would assume *a priori* that individuals are infected in every other time period.

##### 3.2 Beta prior on per-time infection probability

The above prior allows for explicit control over the form of $P(\phi_j)$, but also results in a large number of additional nuisance parameters that must be estimated (each $\phi_j$). It is possible to calculate the marginal distribution $P(\boldsymbol{Z})$ under the above prior by integrating out $\phi$. In terms of MCMC mixing, integrating over possible all possible $\phi$ for each $j$ reduces the number of free parameters to be estimated rather than needing to estimate each $\phi$. This is particularly useful because inferring posterior distributions for $\phi$ and $Z$ simultaneously based on Equation 12 is practically difficult due to their clear correlation, particularly when $m$ is large. The reader is referred to related work on the Indian Buffet Process: a stochastic process defining a probability distribution over sparse binary matrices with finite rows and infinite columns [5]. Our problem is the related case where binary matrices are not necessarily sparse, and the number of columns can be considered finite. However, there is a potential avenue for infection history inference where the number of infection periods (the number of columns) is not necessarily fixed and finite, and we therefore refer to this work here.

Similar to prior version 1, we define a beta-Bernoulli process for the generation of $Z$ as:

$$\phi_j \sim \text{Beta}(\alpha, \beta) \tag{13}$$

$$Z_{i,j} \sim \text{Bernoulli}(\phi_j) \tag{14}$$

The prior probability of $\phi_j$ is defined as:

$$P(\phi_j) = \frac{\phi_j^{\alpha-1}(1-\phi_j)^{\beta-1}}{B(\alpha,\beta)} \tag{15}$$

where $B(\alpha, \beta)$ is the Beta function:

$$B(\alpha,\beta) = \int_0^1 \phi_j^{\alpha-1}(1-\phi_j)^{\beta-1} d\phi_j \tag{16}$$

$$= \frac{\Gamma(\alpha)\Gamma(\beta)}{\Gamma(\alpha+\beta)} \tag{17}$$

$Z_{i,j}$ is independent of all other entries in $\boldsymbol{Z}$, conditional on $\phi_j$ which are also assumed to be independent of all other
*bmphi*. $P(Z_{i,j})$ can then be calculated directly by integrating over all $\boldsymbol{\phi}$, giving the marginal likelihood of the entire infection history matrix $\boldsymbol{Z}$ as:

$$P(\boldsymbol{Z}) = \prod_{j=1}^{m} \int_0^1 \Big(\prod_{i=1}^{n} P(Z_{i,j}|\phi_j)\Big) P(\phi_j) d\phi_j \tag{18}$$

$$= \prod_{j=1}^{m} \frac{B(k_j+\alpha, \beta+n_j-k_j)}{B(\alpha,\beta)} \tag{19}$$

In the MCMC framework, we may propose new values for each $Z_{i,j}$ directly from this prior as:

$$P(Z_{i,j}=1|\boldsymbol{Z_{-i,j}},\alpha,\beta) = \int_0^1 P(Z_{i,j}|\phi_j)P(\phi_j|\boldsymbol{Z_{-i,j}})d\phi_j \tag{20}$$

$$= \frac{k_{-i,j}+\alpha}{n_{-i,j}+\alpha+\beta} \tag{21}$$

Giving the proposal probability of proposing $Z_{i,j} = 1$ and proposing $Z_{i,j} = 0$ otherwise, where $k_{-i,j}$ is the number of infections during time $j$ less $Z_{i,j}$, and $n_{-i,j}$ is the number of individuals alive during time $j$ less individual $i$. The acceptance probability then just becomes the ratio of likelihoods in the Metropolis step.

We may choose values for $\alpha$ and $\beta$ to give a prior on the attack rate with known properties: $\mathbb{E}(k_j) = n\frac{\alpha}{\alpha+\beta}$ and $\mathrm{Var}(k_j) = n\frac{\alpha\beta}{(\alpha+\beta)^2}[1+(n-1)\frac{1}{\alpha+\beta+1}]$. When $\alpha = \beta$, the attack rate prior has an expectation of $0.5n$, and the variance may be decreased by increasing $\alpha$ and $\beta$. `Serosolver` includes functions to calculate values of $\alpha$ and $\beta$ that have a desired mode and certainty by solving the following:

$$\alpha = Mo(c-2) + 1 \tag{22}$$

$$\beta = (1 - Mo)(c-2) + 1 \tag{23}$$

Where $Mo$ is the desired mode, and $c$ is analogous to the number of prior observations (ie. $c = 2$ corresponds to having seen two prior outcomes). `Serosolver` also includes a function to find values for $\alpha$ and $\beta$ that have a particular mean with the largest possible variance by solving:

$$\alpha = \bar{\phi}^2 \frac{1 - \bar{\phi}}{\sigma_\phi - \frac{1}{\bar{\phi}}} \tag{24}$$

$$\beta = \alpha \frac{1}{\bar{\phi} - 1} \tag{25}$$

where $\bar{\phi}$ is the desired mean attack rate, and $\sigma_\phi$ is the maximum variance for $\phi$ that results in a uni-modal distribution of $\phi$. Note that values of $\alpha$ and $\beta$ may be set that lead to a multi-modal of $\phi$ eg. $\alpha = \beta = \frac{1}{2}$.

###### 3.2.1 Prior on number of lifetime infections

The implicit prior on an individual's number of lifetime infections is the same as in Section 3.1: the beta prior on $P(\phi)$ with parameters $\alpha$ and $\beta$ and results in a binomial distribution on the total number of lifetime infections with mean $\frac{\alpha}{\alpha+\beta}$ and $N = m_i$, where $m_i$ is the number of time periods that individual $i$ could be infected.

##### 3.3 Beta prior on per-individual infection probabilities

Under this prior, each individual's prior probability of infection is drawn from a Bernoulli distribution with independent $p_i$ for all $i$, but the same $p_i$ for individual $i$ across all times $j$. This prior captures the idea that individuals may have a tendency to get infected more or less frequently, but the probability of an individual becoming infected is independent of all other individuals. Similar to prior version 2, we model this with a beta-Bernoulli process where the probability of infection as a random variable $p_i$. We defined $Z_{i,j} \sim Ber(p_i)$ with probability $p_i \sim B(\alpha, \beta)$, where $Ber$ is the Bernoulli distribution and $B$ is the beta distribution (the conjugate prior to the Bernoulli distribution). This places a beta prior on the per-time probability of infection, assuming that each individual has a unique $p_i$. The prior probability of a particular infection history for individual $i$, $P(\boldsymbol{Z_i})$, is therefore given by the standard beta-Bernoulli distribution where $p_i$ is treated as a random variable (but does not need to be explicitly estimated, as in prior version 2).

Under this prior, the infection generating process is assumed to be:

$$p_i \sim \text{Beta}(\alpha, \beta) \tag{26}$$
$$Z_{i,j} \sim \text{Bernoulli}(p_i) \tag{27}$$

If the prior on $p_i$ is:

$$P(p_i) = \frac{1}{B(\alpha,\beta)} p_i^{\alpha-1}(1-p_i)^{\beta-1} \tag{28}$$

and the conditional probability of $\boldsymbol{Z_i}$ given $p_i$ is:

$$P(\boldsymbol{Z_i}|p_i) = p_i^k(1-p_i)^{m-k} \tag{29}$$

then the marginal distribution of $P(\boldsymbol{Z_i})$ is:

$$\begin{aligned} P(\boldsymbol{Z_i}) &= \mathbb{E}[P(\boldsymbol{Z_i}|p_i)] && (30)\\ &= \int_0^1 p_i^k(1-p_i)^{m-k}P(p_i)dp_i && (31)\\ &= \frac{B(\alpha+k, \beta+m-k)}{B(\alpha,\beta)} && (32)\\ &= \frac{\alpha^{[k]}\beta^{[m-k]}}{(\alpha+\beta)^{[m]}} && (33) \end{aligned}$$

where $\boldsymbol{Z_i} = (Z_{i,1}, Z_{i,2}, \ldots, Z_{i,m})$, $k$ is the total number of infections experienced by individual $i$ ($\sum_{j=1}^m Z_{i,j}$), $m$ is the number of time periods that individual $i$ could be infected in, and $r^{[x]}$ denotes the ascending power $r(r+1)\ldots[r+(x-1)]$. The probability mass function for the total number of infections $k_i$ is therefore given by:

$$P(k_i, m_i|\alpha,\beta) = \binom{m_i}{k_i}\frac{B(\alpha+k_i, \beta+m_i-k_i)}{B(\alpha,\beta)} \tag{34}$$

which is the beta-binomial distribution. This prior makes the following assumptions:

1. Each $p_i$ comes from a single draw from the same beta distribution
2. All $p_i$ are equal for a given $i$ ie. all $Z_{i,j}$ are drawn from the same Bernoulli distribution
3. All $p_j$ are independent for a given $j$ ie. each $p_j$ is drawn from a different distribution for each $j$.

Formulating the prior in this way allows an explicit prior to be defined through $\alpha$ and $\beta$ on a particular infection history $Z_i$, with $\mathbb{E}(k_i) = m\frac{\alpha}{\alpha+\beta}$ and $\text{Var}(k_i) = m\frac{\alpha\beta}{(\alpha+\beta)^2}[1+(m-1)\frac{1}{\alpha+\beta+1}]$. An intuitive uniform prior on an infection history would therefore be that any total number of lifetime infections is equally likely, which is the case where $\alpha=\beta=1$. In addition, as $\lim \alpha = \beta \to \infty$, $P(k,m|\alpha,\beta) \to \text{Binom}(k,m)$, where any infection history is equally likely. A more informative prior on $\boldsymbol{Z_i}$ is also possible by choosing values for $\alpha$ and $\beta$ that give a desired mean and variance on the total number of infections per individual. I assume that $\alpha$ and $\beta$ are the same for all individuals.

###### 3.3.1 Prior on attack rates

The assumption of independent individuals and a beta-Bernoulli prior on the total number of lifetime infections places a binomial prior on the attack rate within a given time period $j$ across $n$ individuals. The marginal likelihood of infection in an individual's infection history is the same across all individuals, such that:

$$
\begin{aligned}
P(Z_{i,j} = 1|P_{i,j} = p_{i,j}) &= p_{i,j} && (35)\\
P(P_{i,j} = p_{i,j}) &= \frac{p_{i,j}^{\alpha-1}(1-p_{i,j})^{\alpha-1}}{B(\alpha,\beta)} && (36)\\
P(Z_{i,j} = 1) &= \int_0^1 P(Z_{i,j} = 1|P_{i,j} = p_{i,j})P(P_{i,j} = p_{i,j})dp_{i,j} && (37)\\
&= \int_0^1 p_{i,j}\frac{p_{i,j}^{\alpha-1}(1-p_{i,j})^{\alpha-1}}{B(\alpha,\beta)}dp_{i,j} && (38)\\
&= \mathbb{E}(p_{i,j}) && (39)\\
&= \frac{\alpha}{\alpha+\beta} && (40)
\end{aligned}
$$

Given our assumption that individuals are independent and therefore all $p_{i,j}$ are independent across $j$, and that $\alpha$ and $\beta$ are the same for all individuals $i$, then it follows that $P(\sum \boldsymbol{Z_j} = Z_{1,j} + Z_{2,j} + \dots + Z_{n,j})$ is binomially distributed with probability $\frac{\alpha}{\alpha+\beta}$ and $N = n$, the number of individuals. Importantly, $P(\sum \boldsymbol{Z_j})$ is binomially distributed with mean $0.5n$ for all $\alpha = \beta$, even in the case where $\alpha = \beta = 1$.

This prior would suggest that the infection status of individual $n+1$ during time $j$ follows the Bernoulli distribution with $p_j = \frac{\alpha}{\alpha+\beta}$, and the overall number of infections $k_j$ follows the binomial distribution with the same $p_j$ and $N = n_j$. This does fulfil a number of desirable properties: (i) if we know that a proportion $\frac{\alpha}{\alpha+\beta}$ of the population were infected, then the expectation of the attack rate prior would also be $\frac{\alpha}{\alpha+\beta}$; (ii) if $x$ individuals from our sample of $n$ were infected, then we should have less confidence in our prior belief that a random individual in the same population was infected than the case with $100x$ infections from a sample of $100x$, but with the same expectation $\frac{x}{n}$. However, with large $n$ and this binomial prior, the majority of probability density for the attack rate is in a relatively small region of parameter space, resulting in a prior that strongly influences the posterior, and might therefore swamp the likelihood. Furthermore, given that $\mathbb{E}(p_j) = \frac{\alpha}{\alpha+\beta}$, then necessarily $p_1 = p_2 = \dots = p_m$ for all $m$.

###### 3.3.2 Further considerations and Gibbs sampling

In practice, assuming that all $j$ are exchangeable, it is possible to sample $Z_{i,j}$ from the prior directly in a Gibbs-like fashion, which leads to far more efficient proposals. Rather than either moving to a proposed location or staying in the previous location, we can think about the proposal step as offering the algorithm two choices:

1. For an individual $i$, choose a random location, $j$, from the infection history vector, $\boldsymbol{Z_i}$

2. Remove element $j$ to give $Z_{i,-j}$

3. There are now two potential moves to get back to a vector with the same dimensions as $\boldsymbol{Z_i}$. Set $Z_{i,j} = 1$ or $Z_{i,j} = 0$.

Let $\boldsymbol{Z'_i}$ be the case where $Z_{i,j} = 0$ and $Z_i$ be the case where $Z_{i,j} = 1$. More generally, the proposals can be drawn from:

$$P(\text{propose } \boldsymbol{Z_i}) = \frac{g(\boldsymbol{Z_i}|Z_{i,-j})}{g(\boldsymbol{Z_i}|Z_{i,-j}) + g(\boldsymbol{Z'_i}|Z_{i,-j})} \tag{41}$$

In the case of the binomial prior on $k$ (where $P(Z_{i,j} = 1) = 0.5$ when $\alpha = \beta = \infty$), we would have a proposal such that $g(\boldsymbol{Z_i}|\boldsymbol{Z'_i}) = g(\boldsymbol{Z'_i}|\boldsymbol{Z_i}) = g(\boldsymbol{Z'_i}|\boldsymbol{Z'_i}) = g(\boldsymbol{Z_i}|\boldsymbol{Z_i})$. In this case, the probability of proposing $\boldsymbol{Z'_i}$ is the same as the probability of proposing $\boldsymbol{Z_i}$ (ie. 50/50). However, if we explicitly define $g(\boldsymbol{Z_i}|\boldsymbol{Z_{i,-j}})$ and $g(\boldsymbol{Z'_i}|\boldsymbol{Z_{i,-j}})$ then we can control the proposal distribution and therefore the implicit prior:

$$g(\boldsymbol{Z_i}|\boldsymbol{Z_{i,-j}}, \alpha, \beta) = f(Z_{i,j} = 1|\boldsymbol{Z_{i,-j}}, \alpha, \beta) \tag{42}$$
$$g(\boldsymbol{Z'_i}|\boldsymbol{Z_{i,-j}}, \alpha, \beta) = f(Z_{i,j} = 0|\boldsymbol{Z_{i,-j}}, \alpha, \beta) \tag{43}$$

$$f(Z_{i,j} = 1|\boldsymbol{Z_{i,-j}}, \alpha, \beta) = \frac{P(Z_i)}{P(\boldsymbol{Z_{i,-j}})} \tag{44}$$
$$= \frac{\alpha^{[k+1]}\beta^{[m-k]}}{(\alpha+\beta)^{[m+1]}}\frac{(\alpha+\beta)^{[m]}}{\alpha^{[k]}\beta^{[m-k]}} \tag{45}$$
$$= \frac{\alpha+k}{\alpha+\beta+m-1} \tag{46}$$

$$f(Z_{i,j} = 0|\boldsymbol{Z_{i,-j}}) = 1 - f(Z_{i,j} = 1|\boldsymbol{Z_{i,-j}}, \alpha, \beta) \tag{47}$$
$$= \frac{\beta+m-k-1}{\alpha+\beta+m-1} \tag{48}$$

where $k = \sum Z_{i,-j}$, and $\alpha$ and $\beta$ are the left and right parameters of the Beta distribution. Following this proposal (which is equivalent to sampling from the prior for $Z_{i,j}$, the proposal is accepted based on the Metropolis acceptance probability:

$$A(\boldsymbol{Z_{new}}, \boldsymbol{Z_{old}}) = \min(1, \frac{P(\boldsymbol{Y}|\boldsymbol{Z_{new}}, \theta)}{P(\boldsymbol{Y}|\boldsymbol{Z_{old}}, \theta)}) \tag{49}$$

#### 3.4 Beta prior on the probability of any infection event

The final and perhaps most truly "uninformative" prior comes from the assumption that **all** infections are independent and identically distributed events; belonging to a common group or time period is considered irrelevant and the order does not matter. In this case:

$$\phi \sim \text{Beta}(\alpha, \beta) \tag{50}$$
$$Z_{i,j} \sim \text{Bernoulli}(\phi) \tag{51}$$

Such that

$$P(\phi) = \frac{\phi^{\alpha-1}(1-\phi)^{\beta-1}}{B(\alpha,\beta)} \tag{52}$$

$$P(Z_{i,j}|\phi) = \phi^{Z_{i,j}}(1-\phi)^{1-Z_{i,j}} \tag{53}$$

and the marginal likelihood of $\boldsymbol{Z}$ is

$$P(\boldsymbol{Z}) = \int_0^1 \Big(\prod_{j=1}^{m}\prod_{i=1}^{n} P(Z_{i,j}|\phi)\Big)P(\phi)d\phi \tag{54}$$

$$= \frac{B(k+\alpha, \beta+nm-k)}{B(\alpha,\beta)} \tag{55}$$

where $k$ is the total number of infections across all years and individuals and $nm$ is the total number of possible infection events. This gives the conditional probability that a given individual was infected at a given time, and also a proposal distribution:

$$P(Z_{i,j}=1|\boldsymbol{Z_{-i,j}},\alpha,\beta) = \int_0^1 P(Z_{i,j}|\phi)P(\phi|\boldsymbol{Z_{-i,j}})d\phi \tag{56}$$

$$= \frac{k_{-i}+\alpha}{nm_{-i}+\alpha+\beta} \tag{57}$$

This assumption has the desirable property of placing a beta prior on both the total number of infections over a lifetime for a given individual, and on the total number of infections during a given time. However, these desirable properties are traded off against the strong and potentially unrealistic assumption that infection events are conditionally independent across both times and individuals.

**Table 1.** Summary of infection history priors

| Description | Version | Assumption | Summary | Prior on lifetime infections | Prior on augmented attack rates | Use cases |
|---|---|---|---|---|---|---|
| Hyper-prior placed on the probability of infection terms, $\phi$ | 1 | Infection status in a given time correlated with infection status of population at that time, but independent of other times. | The prior probability of any individual $i$ becoming infected during time $j$ is $p_{i,j} = \phi_j$, and all time periods are independent such that each time period $j$ has a unique parameter parameter $\phi_j$ as in Equation 12. These parameters are estimated explicitly and are independent of one another. | Binomial | Beta-binomial | Where $\phi_j$ is of interest and a distinct user-specified prior is desired for each $j$, $P(\phi_j)$. Appropriate when the number of time periods under consideration $j$ is small. Can otherwise lead to poor convergence when $j$ is large. Allows the user full control over the form of $P(\phi_j)$, which is not possible with other versions. |
| Beta prior on per-time probability of infection | 2 | Infection status in a given time correlated with infection status of population at that time, but independent of other times. | Mathematically equivalent to version 1, but $\phi$ is integrated out by placing a conjugate Beta prior on the probability of infection terms. All infection times $j$ are independent, and users may specify the Beta parameters to define the prior distribution. | Binomial, $p = \frac{\alpha}{\alpha+\beta}$ | Beta-binomial with parameters $\alpha$ and $\beta$. | Unbiased attack rate inference. Reason to assume that individuals are under the same probability of infection process eg. same location, but require better MCMC mixing over inferring all $\phi_j$ independently as in version 1. Appropriate when there are a large number of individuals in the sample, but not necessarily a large amount of antibody data per individual. |
| Beta prior on per-individual probability of infection | 3 | Infection status in a given time is independent of infection status of population, but correlated with infection status of that individual at other times. | A Beta prior is placed on the probability of a given individual becoming infected in any time period. However, whereas the above priors assumed independence between times but not individuals, this prior assumes independence between individuals but not between times. Infection probabilities are drawn from a single Beta distribution. | Beta-Binomial with parameters $\alpha$ and $\beta$ | Binomial, $p = \frac{\alpha}{\alpha+\beta}$ | Unbiased per-individual infection history inference. Reason to assume that individuals are under different infection processes, but share antibody kinetics parameters eg. different locations or populations. Appropriate with a relatively small number of individuals and large amount of antibody data per individual. |
| Beta prior on overall probability of infection | 4 | Infection status correlated to all other infection events ie. frequent infection in other individuals and in the past suggests more likely to be infected in the future. | A Beta prior is placed on the probability of any infection, assuming that infection events are independent both across individuals and time periods. | Beta-binomial with parameters $\alpha$ and $\beta$. | Beta-binomial with parameters $\alpha$ and $\beta$. | Weakly informative priors on both attack rates and lifetime infections are desired, over-dispersed relative to the binomial distribution on all summaries. Appropriate with a small number of individuals and relatively small amount of antibody data per individual, as convergence is slower than under other versions. |

### 4 Other considerations

#### 4.1 All infection events independent but not identically distributed

It is possible to also relax the assumption that individual infection probabilities are correlated within years or individuals by considering a unique force of infection term for each $j$ and $i$ ie. placing a beta prior on each $\phi_{i,j}$. However, this would be a redundant model, as each $\phi$ influences only one $Z$ and vice versa. One could instead directly place a Bernoulli prior on each $Z$ with $p = \frac{\alpha}{\alpha+\beta}$ (ie. by integrating out $p$). In this case, assuming that all $Z_{i,j}$ are independent and drawn from different distributions would mean that the total number of infections across all times and individuals is binomially distributed (as this would be the sum of $nm$ independent Bernoulli random variables), which as discussed above is problematic for our inference.

#### 4.2 Prior on total number of infections

The analytical form of the prior $P(\sum_i \sum_j Z_{i,j} = k_{i,j})$ is not well defined, and is therefore not derived here. This can be thought of as either the sum of independent beta-binomial or binomial variables (with different $p$). In the case of summing $n$ binomial variables each of size $m$ with $p = p_j$, the result is also binomially distributed with $p = p_j$ and $N = nm$ when all $p_j$ are equal. However, when all $p_j$ are non-identical, the result becomes more complicated and must be either approximated or calculated numerically [6, 7]. Other approaches to defining this quantity might involve considering a Poisson-binomial random variable with a beta distribution on each group of $p$ (depending on whether the prior is on attack rates or total lifetime infections), or approximating each binomial with a Gaussian and summing. However, given that the total number of infections in $nm$ is not a quantity of interest relative to the per time or per individual number of infections, this is left for future work.

#### 4.3 Comparison with simulation results

Figures 1 and 2 compare infection histories simulated from these priors compared to the analytical priors for summations across different dimensions. These results are intended to support the derivations given above and to highlight the impact of choosing different values for the hyper parameters $\alpha$ and $\beta$ on the inferred attack rate, total number of lifetime infections and overall number of infections. In all simulations, $n = 100$ and $m$=60. Note that in the rightmost panel in all these plots, the distribution for the total across $nm$ is not defined, but resembles an over-dispersed binomial distribution.

Figures 3 and 4 show the prior on an individual's cumulative number of lifetime infections by running the full MCMC framework under each prior with the contribution of the likelihood set to 0 (ie. sampling from the priors). In Figure 3, a uniform beta prior is used with $\alpha = \beta = 1$, though we see that this leads to a strong binomial prior on the total number of lifetime infections in the top two panels, despite a uniform prior on the probability of infection in any given year as in Figure 1. Conversely, in the bottom two panels with the same beta parameters, the prior on the total number

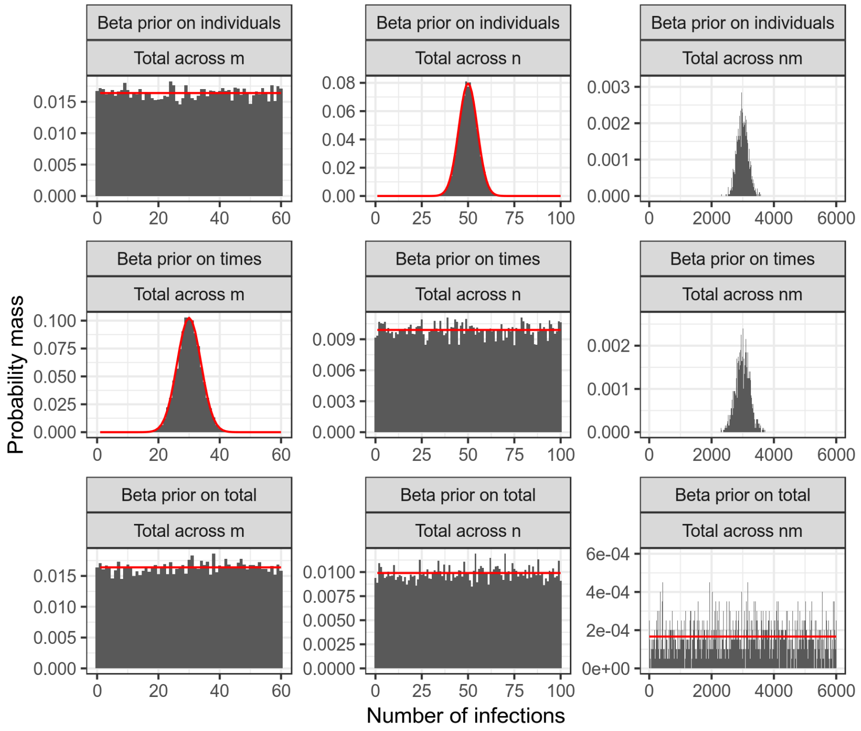


**Fig 1. Simulated vs. known prior when $\alpha = \beta = 1$.** Bars show density histogram of infections from 20000 simulated infection histories. Red line shows known probability mass function. Top row refers to the assumption that a beta prior is placed on the probability of an infection within an individual's infection history, but that individuals are independent. Middle row refers to the assumption that a beta prior is placed on the attack rate within a given time period, and that time periods are independent. Bottom row refers to the assumption that a beta prior is placed on the probability of an infection in any time period for any individual. Leftmost column (total across $m$) refers to the distribution of lifetime infections for one individual. Middle column refers to the distribution on attack rate for a single time period (total across $n$). Rightmost column refers to the distribution of the total number of infections across all times and individuals (total across $nm$).

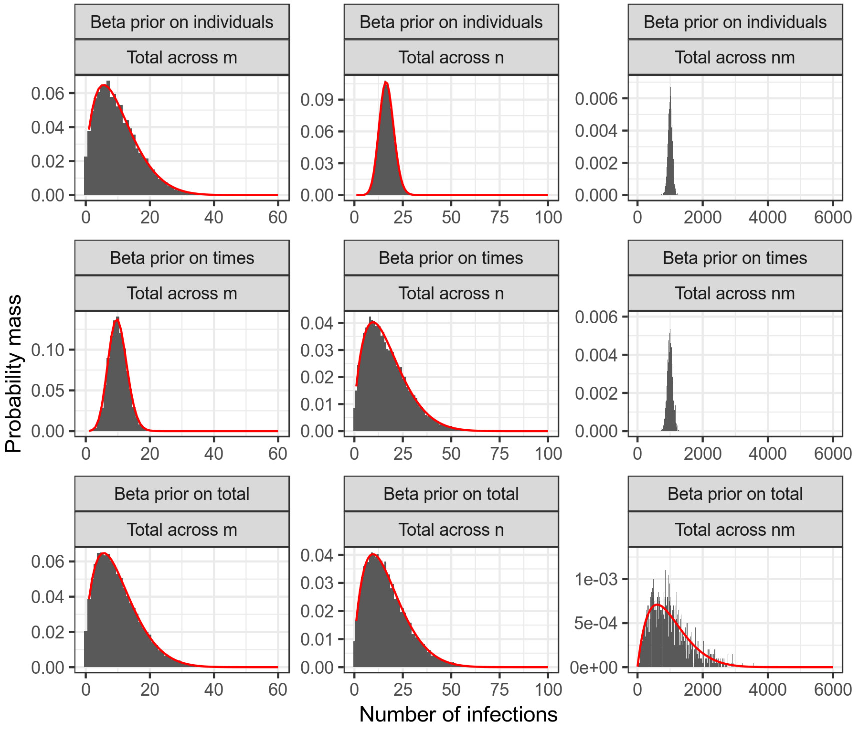


**Fig 2. Simulated vs. known prior when $\alpha = 2$ and $\beta = 10$.** Bars show density histogram of infections from 20000 simulated infection histories. Red line shows known probability mass function. Top row refers to the assumption that a beta prior is placed on the probability of an infection within an individual's infection history, but that individuals are independent. Middle row refers to the assumption that a beta prior is placed on the attack rate within a given time period, and that time periods are independent. Bottom row refers to the assumption that a beta prior is placed on the probability of an infection in any time period for any individual. Leftmost column refers to the distribution of lifetime infections for one individual. Middle column refers to the distribution on attack rate for a single time period. Rightmost column refers to the distribution of the total number of infections across all times and individuals.

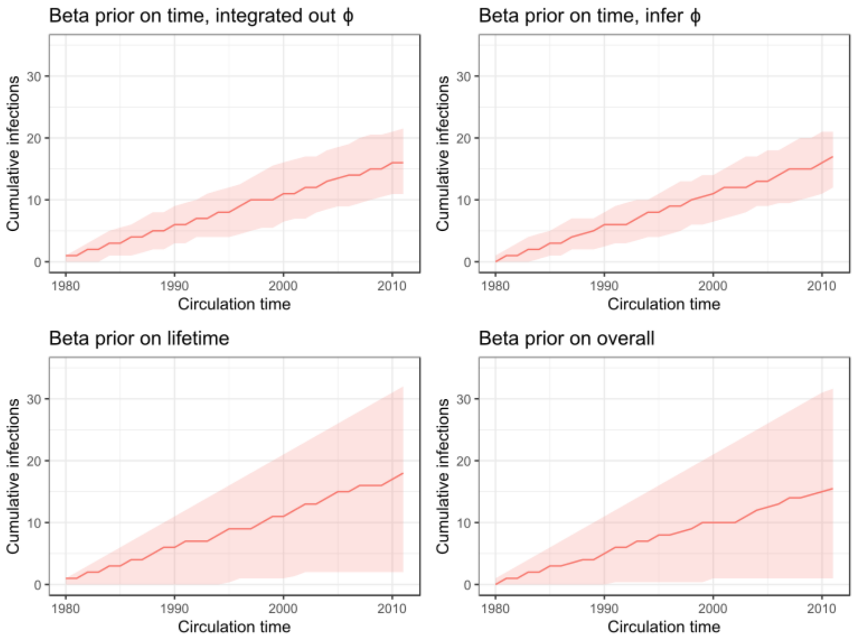


**Fig 3. Priors on cumulative number of lifetime infections when $\alpha = \beta = 1$.** Subplot titles show assumed prior corresponding to the main text. Red line shows posterior median. Red shaded regions show posterior 95% credible intervals.

of lifetime infections follows the beta-binomial distribution. The different shapes and widths of these distributions demonstrates how different prior forms may lead to strong assumptions about an individual's infection history. Figure 4 makes the same comparison, but choosing informative beta parameters of $\alpha = 2$ and $\beta = 10$.

#### 4.4 Choice of prior

If the dimensions of $m$ (few infection times) and $n$ (few individuals) are small and the contribution of the likelihood is large (ie. large amounts of titre data), then the assumptions of these priors have relatively little impact on the inferred infection histories. However, if the amount of data and therefore contribution of the likelihood is small, then the contribution of the prior to inferred infection histories becomes large. In the beta-Bernoulli prior case, a large number of individuals (large $n$) places a very strong binomial prior on the attack rate (total number of infections per time period), whereas a high time resolution (large $m$) (eg. months) has little impact on the a priori total number of lifetime infections per individual. Conversely, in the beta prior case, large $n$ has no impact on the attack rate prior, but does have an impact on the prior for the total number of lifetime infections per individual. The choice of which prior to use therefore depends on the data structure, prior knowledge and scientific aims. For example, inferring antibody kinetics that describe individual titres well across a sample population is readily achieved using the Beta-Bernoulli prior (because data purely informs the infection history that best describes the antibody data), whereas inferring accurate historical attack rates is better suited to the beta prior (because information on infection state is

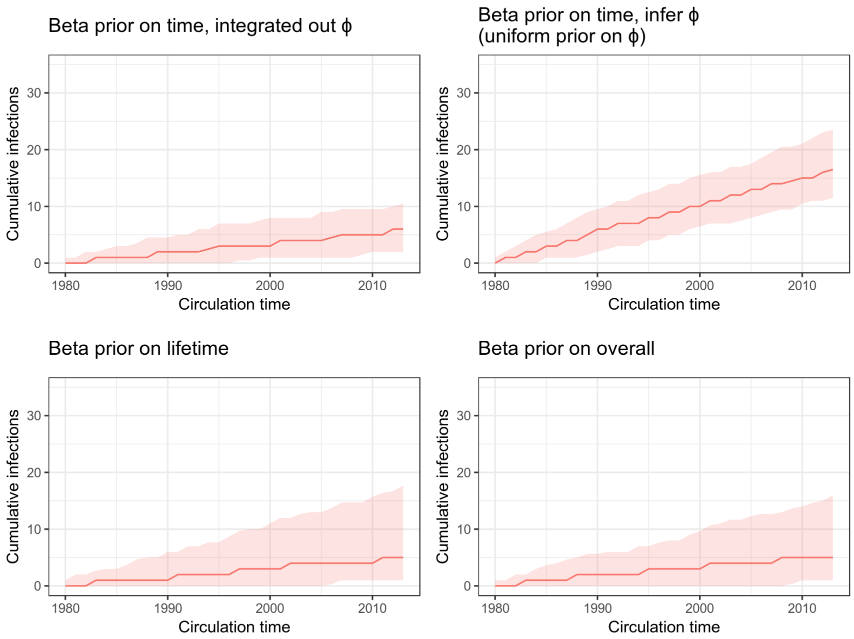


**Fig 4. Priors on cumulative number of lifetime infections when $\alpha = 2$ and $\beta = 10$.** Subplot titles show assumed prior corresponding to the main text. Red line shows posterior median. Red shaded regions show posterior 95% credible intervals. Note that a uniform prior is assumed for $\phi$, and this prior therefore matches that in Figure 3.

shared between individuals).

To illustrate the impact of different prior assumptions on infection history and antibody kinetics parameter inference, we ran simulation-recovery experiments for each for the latter 3 priors with varying amounts of titre data and Beta prior parameters. We did not run simulation recovery for prior version 1, as this is equivalent to prior version 2. The simulated serosurvey design was similar to that of case-study 2 in the main text: 200 individuals of varying ages between 10 and 75 years old; potential annual infections from 1968 to 2009; estimating only long-term antibody kinetics parameters $\mu$ (long term boosting), $\sigma_l$ (long term cross reactivity), $\tau$ (antigenic seniority) and $\epsilon$ (measurement error). For each prior version, we considered three data scenarios: 1) sparse data, only one blood sample taken and titres against 9 viruses taken (as in the real data); 2) full data, one blood sample taken and titres against each of the 41 viruses (one per year); 3) additional data, 5 blood samples taken at random intervals between 2000 and 2009, with 41 viruses tested from each blood sample. These three data scenarios represent a range of low data contribution to the posterior up to extremely high data contribution to the posterior. For each data scenario, we tested 4 prior assumptions: 1) neutral prior with $\alpha = \beta = \frac{1}{3}$; 2) uniform prior with $\alpha = \beta = 1$; 3) weakly informative prior with prior probability of infection mode of 0.15 and high variance, with $\alpha = 1.3$ and $\beta = 2.7$, conceptually similar to a prior informed by 4 prior observations; 4) strongly informative prior with prior probability of infection mode of 0.15 and low variance, with $\alpha = 15.7$ and $\beta = 84.3$, conceptually similar to a prior informed by 100 prior observations. We then ran `serosolver` to generate 5 chains each of 1200000 iterations for these scenarios,discarding the first 200000 iterations as burn in. Note that the same data are used for all prior scenarios.

Figure 5 is particularly revealing: prior versions 2 and 4 recover unbiased estimates of the long-term boosting parameter $\mu$ for all data and prior scenarios, whereas prior version 3 is only unbiased with a large amount of data or strong prior information. Under this survey design, using prior version 2 or 4 would be recommended for estimating long-term dynamics and attack rates. This is supported by Figure 6 and 12, where recovering the true attack rates is shows little bias at all but the strongest prior assumption with very little data. In Figure 7, 8, 13 and 14, attack rate estimation becomes increasingly accurate, and only the scenario with very strong prior information biases the inferred attack rates. Conversely, for prior version 3, only the more substantial data scenarios allow recovery of the general trends of higher and lower attack rates, and even at the highest data contribution these remain biased.

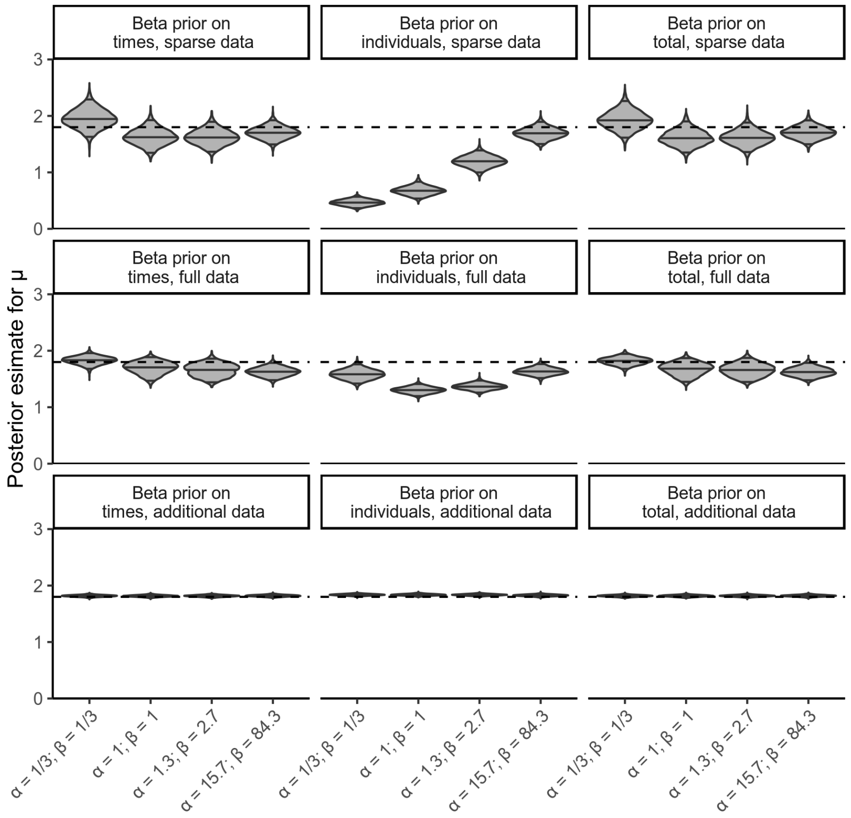


**Fig 5. Posterior distribution of long-term boosting parameters $\mu$ under different prior and data scenarios.** Horizontal dashed line shows true parameter value $\mu = 1.8$, shaded regions show inferred posterior distribution with medians and 95% credible intervals shown as solid horizontal lines. X-axis shows assumed beta prior parameters. Left-hand column shows results under prior version 2; middle column shows results under prior version 3; right-hand column shows results under prior version 4.

Figures 15-23 show the ability of these different data and prior scenarios to infer the same individual's infection history. Prior versions 2 and 4 are able to accurately recover the timing of that individual's infections even under the neutral and uniform priors, with increasing certainty in the more data rich scenarios (Figures 17&23). With sparse data, prior version 3 does not recover constrained posterior estimates for the cumulative infection history under all but the strongest prior (Figure 18). However, under the more data rich scenarios, prior version 3 is able to recover unbiased estimates of the true cumulative infection history, despite clear bias in the inferred attack rates (Figure 23&14). These results highlight that these different priors have their uses depending on the distribution of titre data, the resolution of the model and the particular question under consideration: data sets may be rich in different dimensions (eg. number of individuals vs. number of viruses tested), which leads to different levels of inferential accuracy for different quantities. For example, a data set with very few individuals but a large number of tested titres per individual may

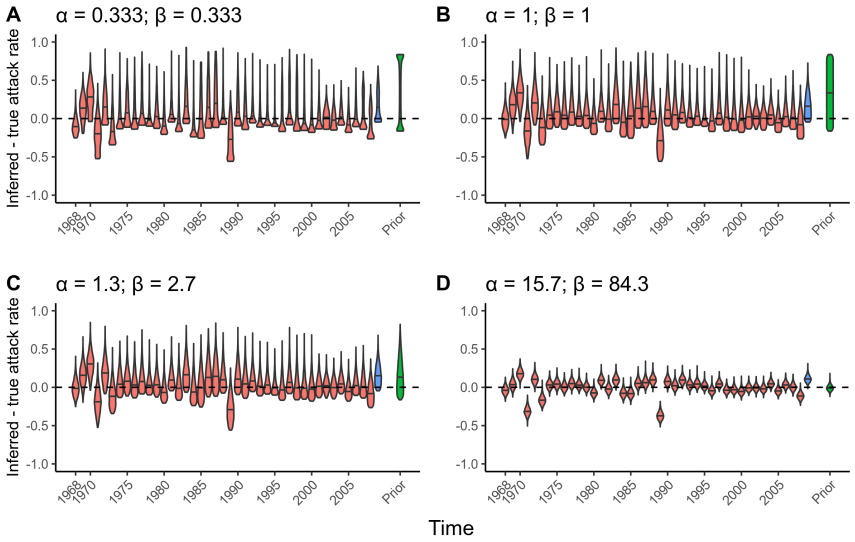


**Fig 6. Simulation-recovery of attack rates using prior version 2 with various strength priors, sparse data.** Simulation with 200 individuals, 9 viruses tested for each individual, one blood sample taken. Y-axis shows inferred attack rate minus the true attack rate. Red violin plot show the inferred posterior distribution of these attack rate residuals in years where no blood sample was taken, whereas blue violin plots show the attack rate residuals in years where a blood sample was taken. Green violin plot shows the empirical prior minus the mean true attack rate across all times. Plot subtitles show assumed beta prior parameters.

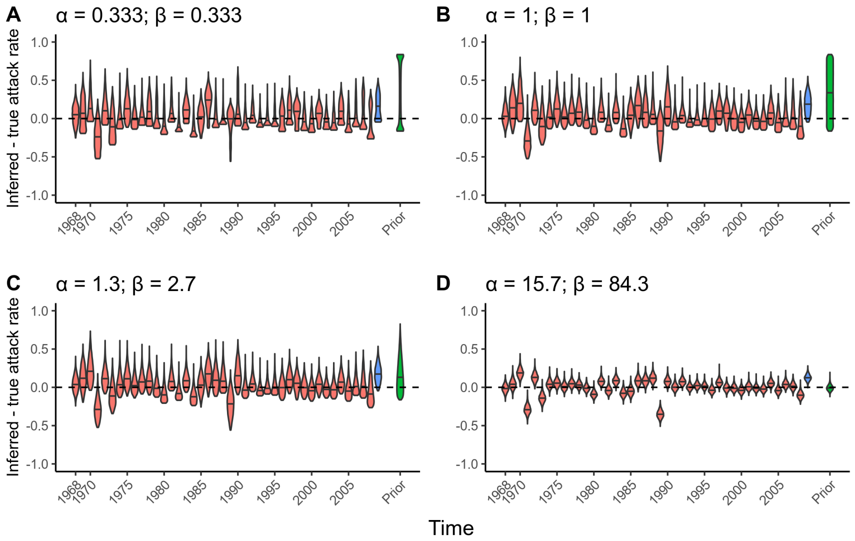


**Fig 7. Simulation-recovery of attack rates using prior version 2 with various strength priors, full data.** Simulation with 200 individuals, 41 viruses tested for each individual, one blood sample taken. Y-axis shows inferred attack rate minus the true attack rate. Red violin plot show the inferred posterior distribution of these attack rate residuals in years where no blood sample was taken, whereas blue violin plots show the attack rate residuals in years where a blood sample was taken. Green violin plot shows the empirical prior minus the mean true attack rate across all times. Plot subtitles show assumed beta prior parameters.

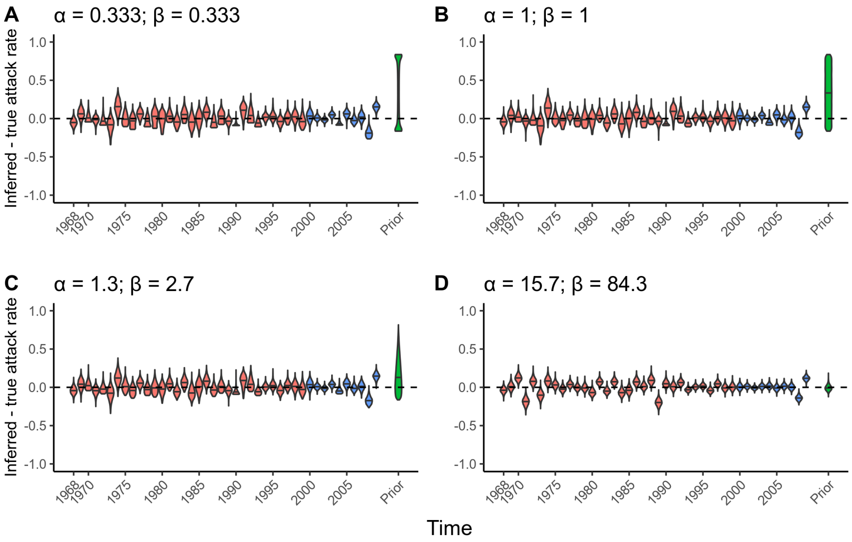


**Fig 8. Simulation-recovery of attack rates using prior version 2 with various strength priors, rich data.** Simulation with 200 individuals, 41 viruses tested for each individual, 5 blood samples taken. Y-axis shows inferred attack rate minus the true attack rate. Red violin plot show the inferred posterior distribution of these attack rate residuals in years where no blood sample was taken, whereas blue violin plots show the attack rate residuals in years where a blood sample was taken. Green violin plot shows the empirical prior minus the mean true attack rate across all times. Plot subtitles show assumed beta prior parameters.

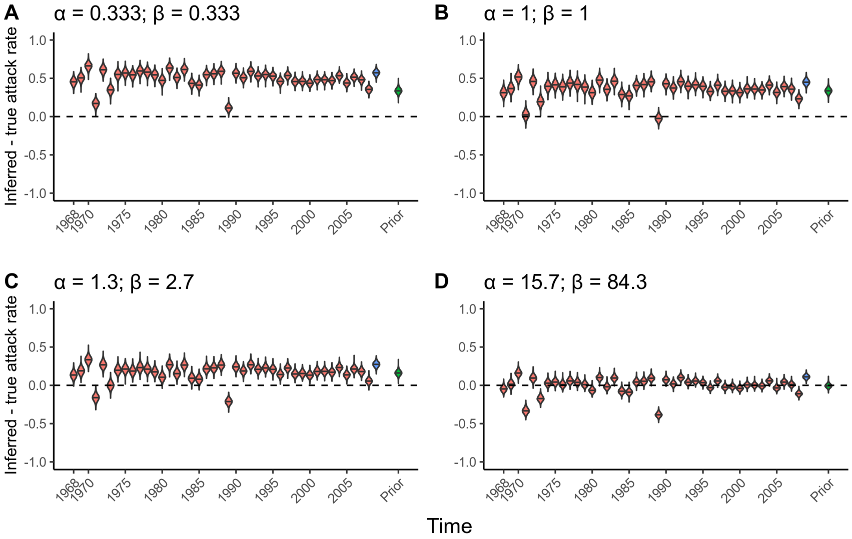


**Fig 9. Simulation-recovery of attack rates using prior version 3 with various strength priors, sparse data.** Simulation with 200 individuals, 9 viruses tested for each individual, one blood sample taken. Y-axis shows inferred attack rate minus the true attack rate. Red violin plot show the inferred posterior distribution of these attack rate residuals in years where no blood sample was taken, whereas blue violin plots show the attack rate residuals in years where a blood sample was taken. Green violin plot shows the empirical prior minus the mean true attack rate across all times. Plot subtitles show assumed beta prior parameters.

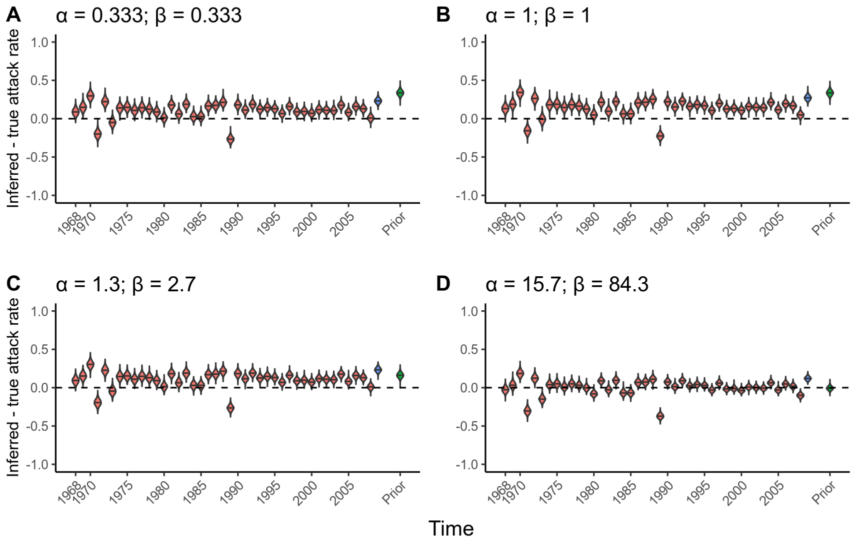


**Fig 10. Simulation-recovery of attack rates using prior version 3 with various strength priors, full data.** Simulation with 200 individuals, 41 viruses tested for each individual, one blood sample taken. Y-axis shows inferred attack rate minus the true attack rate. Red violin plot show the inferred posterior distribution of these attack rate residuals in years where no blood sample was taken, whereas blue violin plots show the attack rate residuals in years where a blood sample was taken. Green violin plot shows the empirical prior minus the mean true attack rate across all times. Plot subtitles show assumed beta prior parameters.

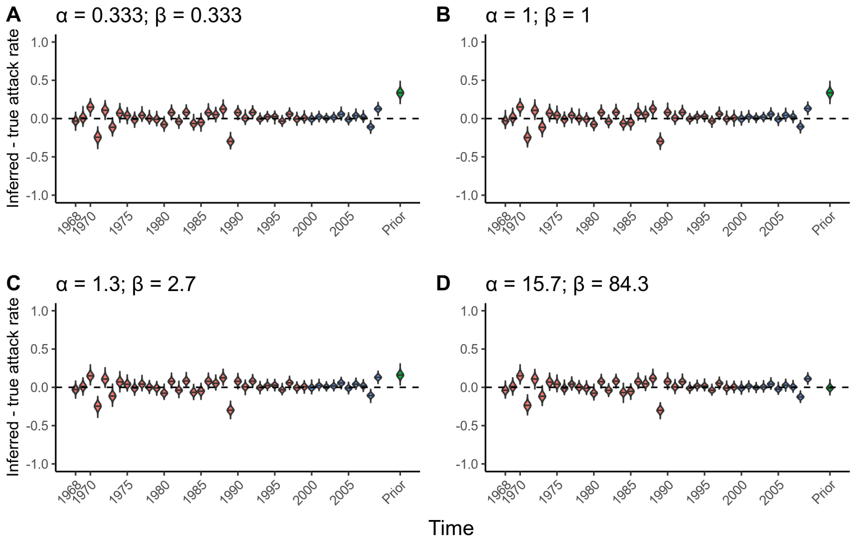


**Fig 11. Simulation-recovery of attack rates using prior version 3 with various strength priors, rich data.** Simulation with 200 individuals, 41 viruses tested for each individual, 5 blood samples taken. Y-axis shows inferred attack rate minus the true attack rate. Red violin plot show the inferred posterior distribution of these attack rate residuals in years where no blood sample was taken, whereas blue violin plots show the attack rate residuals in years where a blood sample was taken. Green violin plot shows the empirical prior minus the mean true attack rate across all times. Plot subtitles show assumed beta prior parameters.

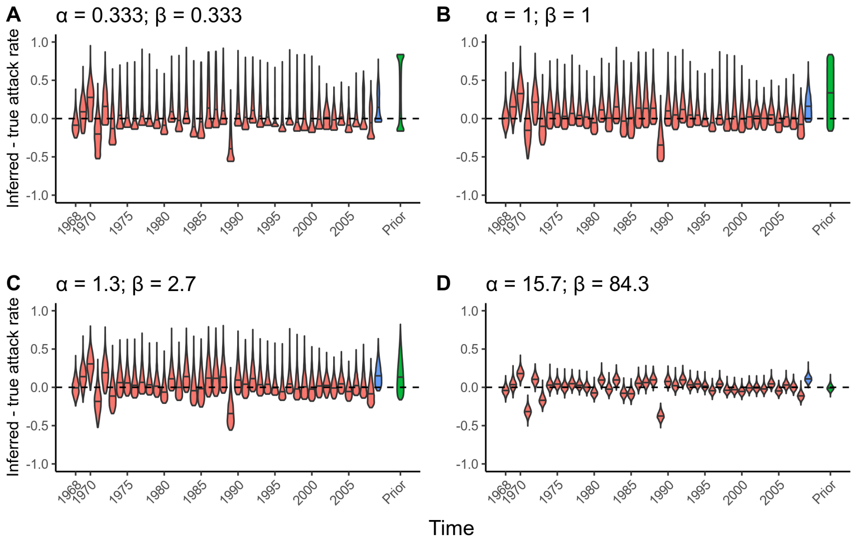


**Fig 12. Simulation-recovery of attack rates using prior version 4 with various strength priors, sparse data.** Simulation with 200 individuals, 9 viruses tested for each individual, one blood sample taken. Y-axis shows inferred attack rate minus the true attack rate. Red violin plot show the inferred posterior distribution of these attack rate residuals in years where no blood sample was taken, whereas blue violin plots show the attack rate residuals in years where a blood sample was taken. Green violin plot shows the empirical prior minus the mean true attack rate across all times. Plot subtitles show assumed beta prior parameters.

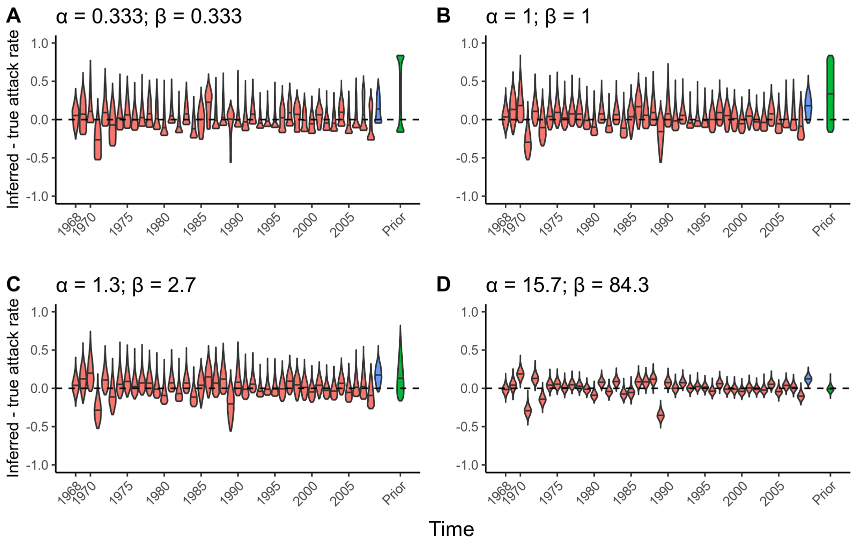


**Fig 13. Simulation-recovery of attack rates using prior version 4 with various strength priors, full data.** Simulation with 200 individuals, 41 viruses tested for each individual, one blood sample taken. Y-axis shows inferred attack rate minus the true attack rate. Red violin plot show the inferred posterior distribution of these attack rate residuals in years where no blood sample was taken, whereas blue violin plots show the attack rate residuals in years where a blood sample was taken. Green violin plot shows the empirical prior minus the mean true attack rate across all times. Plot subtitles show assumed beta prior parameters.

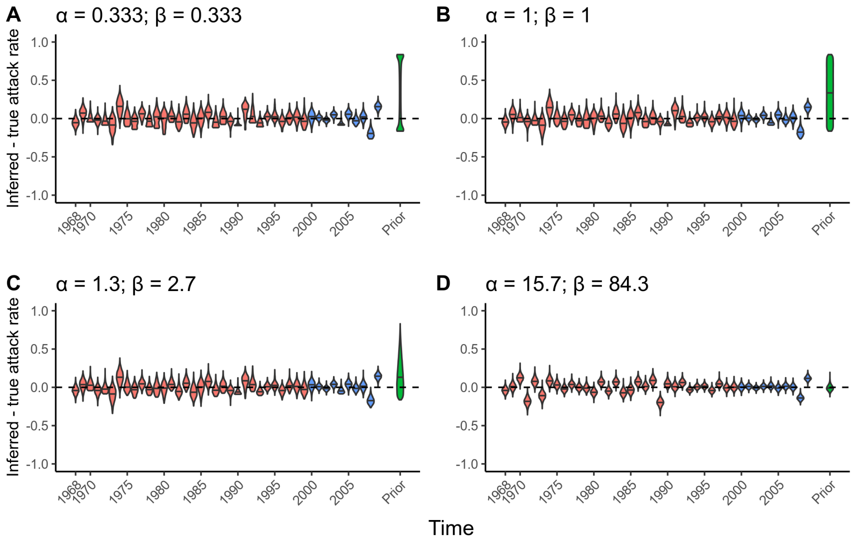


**Fig 14. Simulation-recovery of attack rates using prior version 4 with various strength priors, rich data.** Simulation with 200 individuals, 41 viruses tested for each individual, 5 blood samples taken. Y-axis shows inferred attack rate minus the true attack rate. Red violin plot show the inferred posterior distribution of these attack rate residuals in years where no blood sample was taken, whereas blue violin plots show the attack rate residuals in years where a blood sample was taken. Green violin plot shows the empirical prior minus the mean true attack rate across all times. Plot subtitles show assumed beta prior parameters.

be well suited to analysis under prior version 3 where the aim is to infer an individual's lifetime infection history, but not necessarily population-level attack rates.

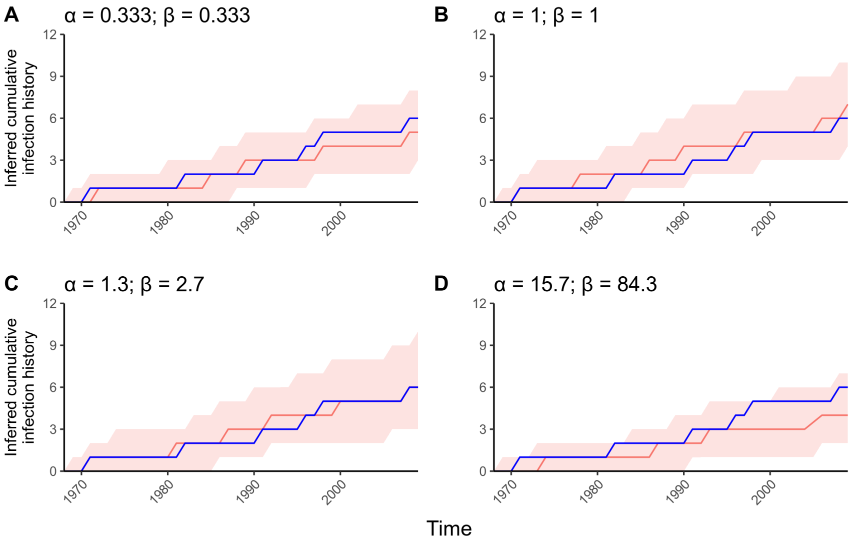


**Fig 15. Simulation-recovery of one individual's infection history using prior version 2 with various strength priors, sparse data.** Simulation with 200 individuals, 9 viruses tested for each individual, one blood sample taken. Y-axis shows the cumulative number of infections for this individual over time. Red line and shaded region shows posterior median and 95% credible intervals. Blue line shows the true cumulative number of infections over time.

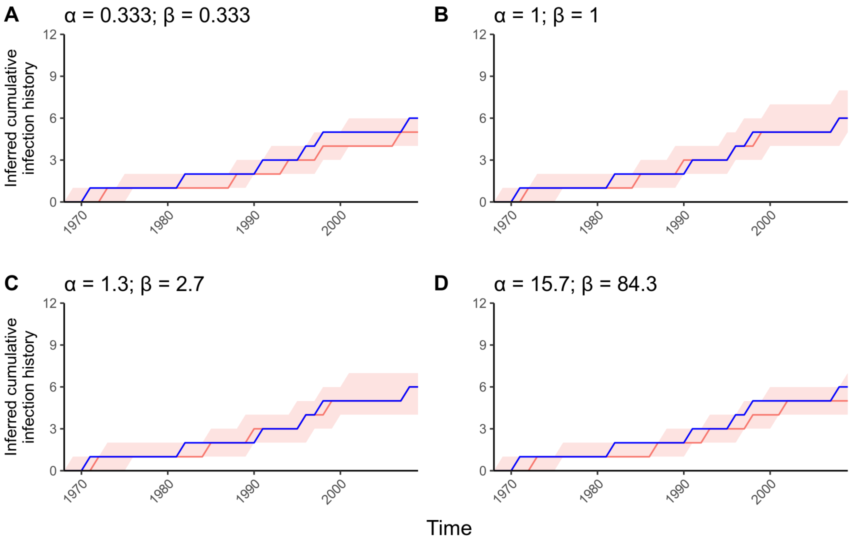


**Fig 16. Simulation-recovery of one individual's infection history using prior version 2 with various strength priors, full data.** Simulation with 200 individuals, 41 viruses tested for each individual, one blood sample taken. Y-axis shows the cumulative number of infections for this individual over time. Red line and shaded region shows posterior median and 95% credible intervals. Blue line shows the true cumulative number of infections over time.

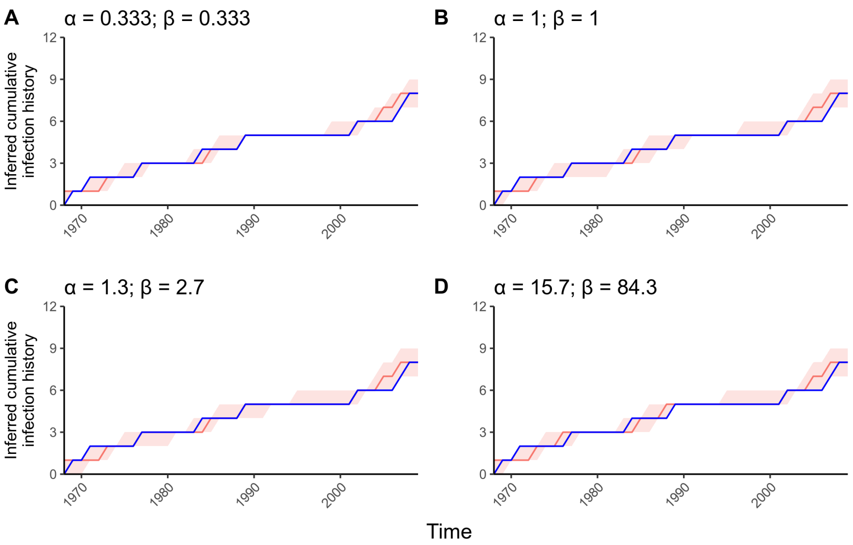


**Fig 17. Simulation-recovery of one individual's infection history using prior version 2 with various strength priors, rich data.** Simulation with 200 individuals, 41 viruses tested for each individual, 5 blood samples taken. Y-axis shows the cumulative number of infections for this individual over time. Red line and shaded region shows posterior median and 95% credible intervals. Blue line shows the true cumulative number of infections over time.

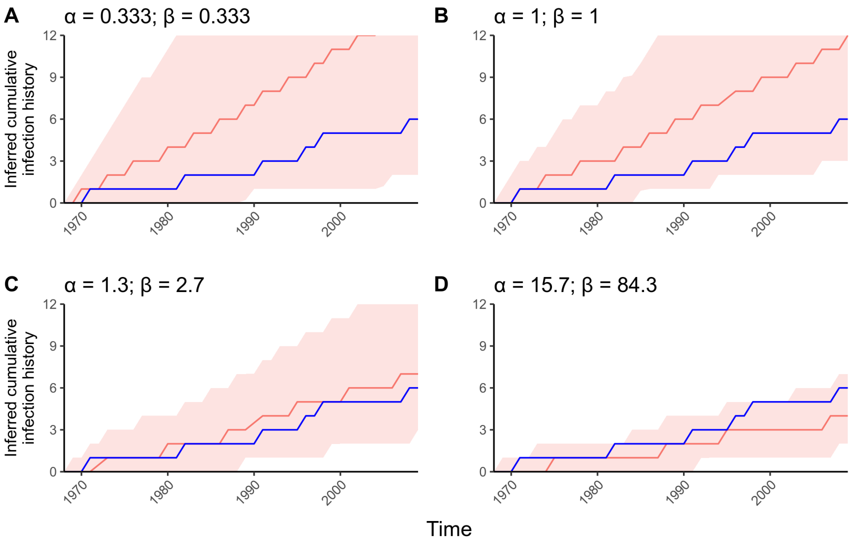


**Fig 18. Simulation-recovery of one individual's infection history using prior version 3 with various strength priors, sparse data.** Simulation with 200 individuals, 9 viruses tested for each individual, one blood sample taken. Y-axis shows the cumulative number of infections for this individual over time. Red line and shaded region shows posterior median and 95% credible intervals. Blue line shows the true cumulative number of infections over time.

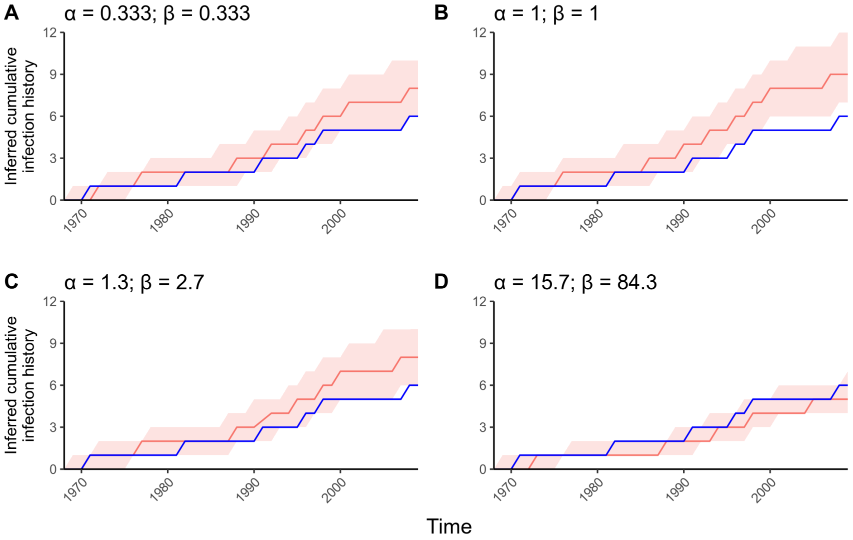


**Fig 19. Simulation-recovery of one individual's infection history using prior version 3 with various strength priors, full data.** Simulation with 200 individuals, 41 viruses tested for each individual, one blood sample taken. Y-axis shows the cumulative number of infections for this individual over time. Red line and shaded region shows posterior median and 95% credible intervals. Blue line shows the true cumulative number of infections over time.

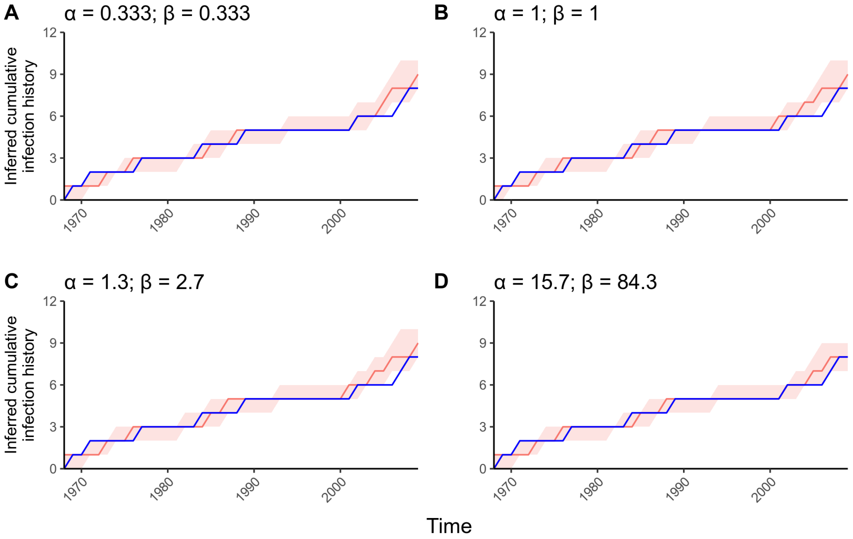


**Fig 20. Simulation-recovery of one individual's infection history using prior version 3 with various strength priors, rich data.** Simulation with 200 individuals, 41 viruses tested for each individual, 5 blood samples taken. Y-axis shows the cumulative number of infections for this individual over time. Red line and shaded region shows posterior median and 95% credible intervals. Blue line shows the true cumulative number of infections over time.

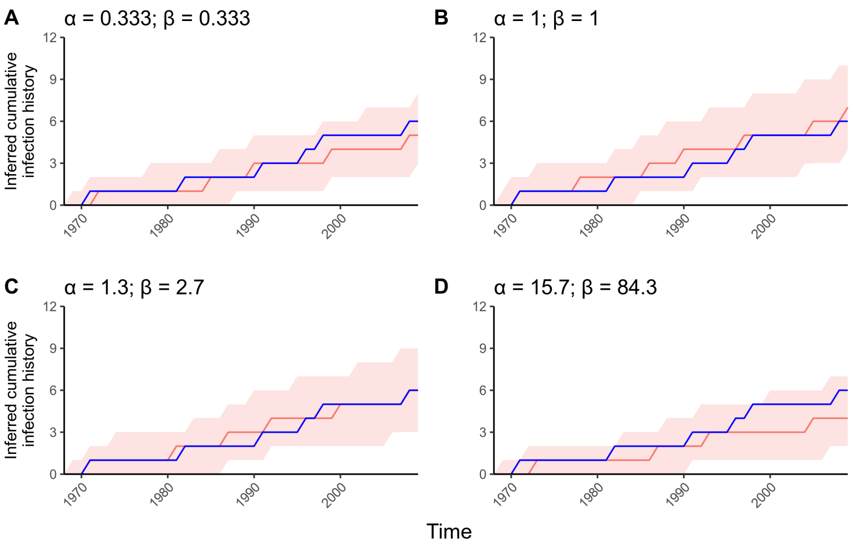


**Fig 21. Simulation-recovery of one individual's infection history using prior version 4 with various strength priors, sparse data.** Simulation with 200 individuals, 9 viruses tested for each individual, one blood sample taken. Y-axis shows the cumulative number of infections for this individual over time. Red line and shaded region shows posterior median and 95% credible intervals. Blue line shows the true cumulative number of infections over time.

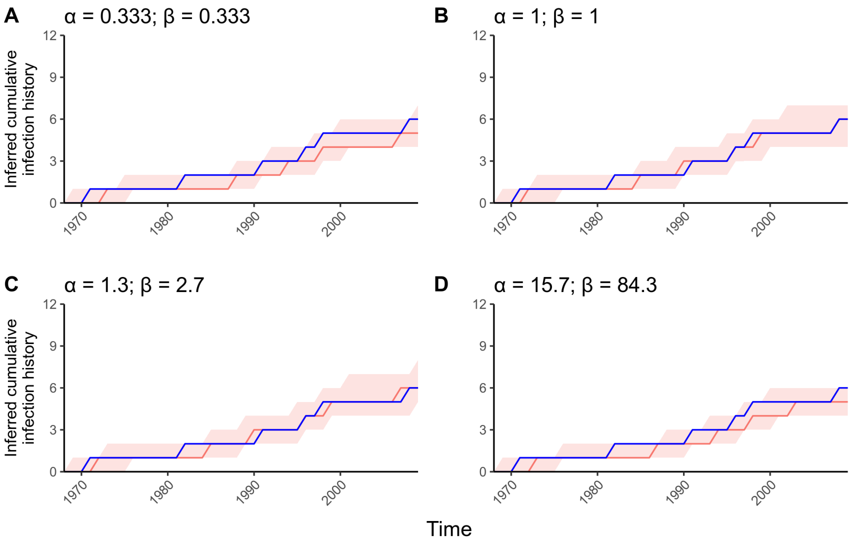


**Fig 22. Simulation-recovery of one individual's infection history using prior version 4 with various strength priors, full data.** Simulation with 200 individuals, 41 viruses tested for each individual, one blood sample taken. Y-axis shows the cumulative number of infections for this individual over time. Red line and shaded region shows posterior median and 95% credible intervals. Blue line shows the true cumulative number of infections over time.

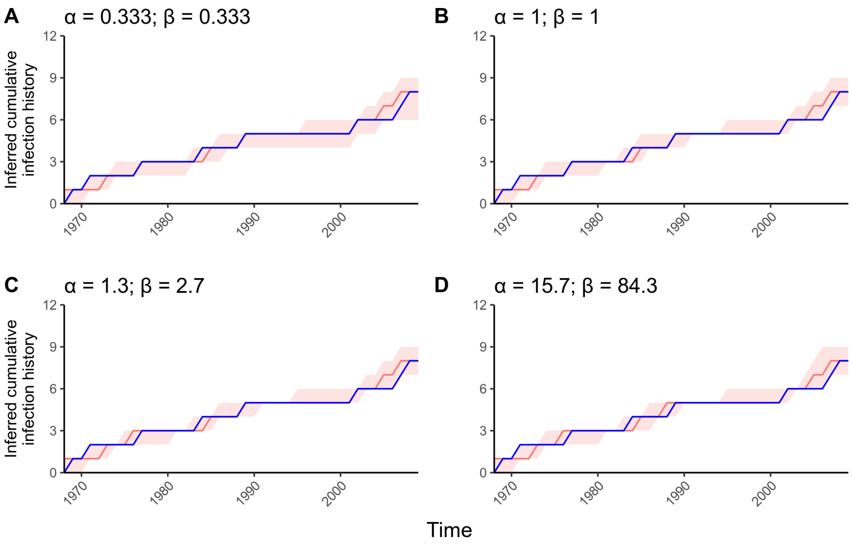


**Fig 23. Simulation-recovery of one individual's infection history using prior version 4 with various strength priors, rich data.** Simulation with 200 individuals, 41 viruses tested for each individual, 5 blood samples taken. Y-axis shows the cumulative number of infections for this individual over time. Red line and shaded region shows posterior median and 95% credible intervals. Blue line shows the true cumulative number of infections over time.

### 5 Appendix

#### 5.1 Proposal algorithm under prior version 1, hyper-prior on probability of infection

In `serosolver`'s MCMC algorithm, all $\phi$ are treated as unknown parameters under this prior and therefore sampled alongside $\theta$. Proposals for $\boldsymbol{Z}$ are made through a random scan across $n$ and $m$, with a proposed transition of flipping the binary entry for $Z_{i,j}$ ie. $Z'_{i,j} = 1$ if $Z_{i,j} = 0$, and $Z'_{i,j} = 0$ if $Z_{i,j} = 1$. The sampling algorithm for $\boldsymbol{Z}$ under the hyper-prior model (section 3.1) is as follows, with italicised terms referring to arguments in *mcmc_pars* of ***serosolver::run_MCMC***:

1. With probability *hist_switch_prob*:
   (a) Select a time point $j$
   (b) Select another time point between 1 and *move_size* time points away from $j$ with uniform probability, where $j \neq l$
   (c) Select *year_swap_propn**$n$ individuals
   (d) For each of these selected individuals, swap the values of $Z_{i,j}$ and $Z_{il}$, adhering to restrictions of birth times (ie. individuals cannot be infected before they are born)
   (e) Increase/decrease $\phi_j$ and $\phi_l$ proportional to the number of infections gained/lost
   (f) Let $\boldsymbol{Z}$ and $\boldsymbol{\phi}$ represent the infection history matrix and probability of infection terms before this proposal step, and $\boldsymbol{Z'}$ and $\boldsymbol{\phi'}$ represent these terms after the proposal step. Set $\boldsymbol{Z} = \boldsymbol{Z'}$ and $\boldsymbol{\phi} = \boldsymbol{\phi'}$ with the following acceptance ratio:

$$A((\boldsymbol{Z'}, \boldsymbol{\phi'}), (\boldsymbol{Z}, \boldsymbol{\phi})) = \min(1, \frac{P(\boldsymbol{Z'}, \theta, \boldsymbol{\phi'}|\boldsymbol{Y})}{P(\boldsymbol{Z}, \theta, \boldsymbol{\phi}|\boldsymbol{Y})}) \tag{58}$$

2. Otherwise:
   (a) Select a random proportion *hist_sample_prob* of $n$ individuals
   (b) For each individual, $i$, propose a new infection history $\boldsymbol{Z'_i}$ by performing one of the following steps:
      i. Perform a "flip" step with probability *1 - swap_propn*:
         A. Select *inf_propn**$m_i$ time points, where $m_i$ is the number of time points that individual $i$ could be infected
         B. Perform a binary flip on each of these times, $Z'_{i,j} = 1 - Z_{i,j}$
      ii. Otherwise, perform a "swap" step:
         A. Select a location $j$
         B. Select a location, $l$, 0 to *move_size* time steps away with equal probability
         C. Set $Z_{i,l} = Z_{i,j}$ and $Z_{i,j} = Z_{i,l}$
   (c) For each sampled individual, independently accept or reject the proposed new infection state with the acceptance ratio:

$$A((\boldsymbol{Z'_i}, \boldsymbol{\phi'}), (\boldsymbol{Z_i}, \boldsymbol{\phi})) = \min(1, \frac{P(\boldsymbol{Z'_i}, \theta, \boldsymbol{\phi'}|\boldsymbol{Y_i})}{P(\boldsymbol{Z_i}, \theta, \boldsymbol{\phi}|\boldsymbol{Y})}) \tag{59}$$

If *hist_opt* is set to 1 by the user, then step 2 above is automatically tuned, whereby *hist_sample_prob* is increased or decreased to achieve a desired acceptance rate (usually between 0.25 and 0.4). It is also possible to manually tune *move_size*, *swap_propn*, *year_swap_propn* and *hist_switch_prob* to improve the acceptance rate of the other proposal steps, though `serosolver` does not currently do this automatically. The acceptance rate of steps 1 and 2 above are printed at regular intervals during the MCMC procedure, which the user may use to tweak these inputs. Further automated tuning remains a direction for further development of the package.

#### 5.2 Proposal algorithm under prior version 2, beta prior on per-time probability of infection

The proposal algorithm for prior version 2 is similar to that of prior version 1, but rather than performing a "flip" step, infection history entries are proposed in a Gibbs-like fashion conditional on the infection status of all other individuals at that time point.

1. With probability *hist_switch_prob*:
    (a) Select a time point $j$
    (b) Select another time point between 1 and *move_size* time points away from $j$ with uniform probability, where $j \neq l$
    (c) Select *year_swap_propn**$n$ individuals and filter for individuals that were alive during both time points
    (d) For each of these selected individuals, swap the values of $Z_{i,j}$ and $Z_{i,l}$
    (e) Let $\boldsymbol{Z}$ and $\boldsymbol{\phi}$ represent the infection history matrix and probability of infection terms before this proposal step, and $\boldsymbol{Z'}$ and $\boldsymbol{\phi'}$ represent these terms after the proposal step. Set $\boldsymbol{Z} = \boldsymbol{Z'}$ and $\boldsymbol{\phi} = \boldsymbol{\phi'}$ with the following acceptance ratio:

$$A(\boldsymbol{Z'}, \boldsymbol{Z}) = \min(1, \frac{\prod_i f(\boldsymbol{Y_i}|\boldsymbol{Z'_i}, \theta)P(\boldsymbol{Z'_i})}{\prod_i f(\boldsymbol{Y_i}|\boldsymbol{Z_i}, \theta)P(\boldsymbol{Z_i})}) \tag{60}$$

2. Otherwise:
    (a) Select a random proportion *hist_sample_prob* of $n$ individuals
    (b) For each individual, $i$, propose a new infection history $\boldsymbol{Z'_i}$ by performing one of the following steps:
        i. Sample new values for $\boldsymbol{Z_i}$ with probability *1 - swap_propn* as follows:
            A. Select *inf_propn**$m_i$ time points, where $m_i$ is the number of time points that individual $i$ could be infected. For each time point, $j$:
            B. Calculate the number of infected individuals less the selected individual $k_j = (\sum_x Z_{x,j}) - Z_{i,j}$
            C. Calculate the number of individuals that could be infected during time $j$, $n_j$
            D. Set $Z_{i,j} = 1$ with probability $\frac{k_j+\alpha}{n_j+\alpha+\beta}$, and $Z_{i,j} = 0$ otherwise

E. Accept the proposed move with the acceptance ratio, noting that by sampling directly from the prior $P(\boldsymbol{Z})$ that this cancels out in the Metropolis ratio:

$$A(\boldsymbol{Z_i'}, \boldsymbol{Z_i}) = \min(1, \frac{f(\mathbf{Y_i}|\boldsymbol{Z_i'}, \theta)}{f(\mathbf{Y_i}|\boldsymbol{Z_i}, \theta)}) \tag{61}$$

ii. Otherwise, perform a "swap" step:

A. Select a location $j$

B. Select a location, $l$, 0 to *move_size* time steps away with equal probability

C. If $Z_{i,l} \neq Z_{i,j}$, set $Z_{i,l} = Z_{i,j}$ and $Z_{i,j} = Z_{i,l}$ with the acceptance ratio:

$$A(\boldsymbol{Z_i'}, \boldsymbol{Z_i}) = \min(1, \frac{f(\mathbf{Y_i}|\boldsymbol{Z_i'}, \theta)P(\boldsymbol{Z_i'})}{f(\mathbf{Y_i}|\boldsymbol{Z_i}, \theta)})P(\boldsymbol{Z_i}) \tag{62}$$

As in prior version 1, if *hist_opt* is set to 1 by the user then *hist_sample_prob* is tuned to achieve a user-specified acceptance rate. It is also possible to manually tune *move_size* and *swap_propn*.

##### 5.3 Proposal algorithm under prior version 3, beta prior on per-individual infection probability

To improve mixing, we extended the logic in Section 3.3.2 to generate a proposal algorithm to add/remove an arbitrary number of infections with each proposal. The full proposal algorithm under prior version 3 is:

1. Select a random proportion *hist_sample_prob* of $n$ individuals

2. For each individual, $i$, propose a new infection history $\boldsymbol{Z_i}$ by performing one of the following steps:

   (a) Sample new values for $\boldsymbol{Z_i'}$ with probability *1 - swap_propn* as follows:

      i. Select $\boldsymbol{z} = \textit{inf\_propn} * m_i$ random time points, where $m_i$ is the number of time points that individual $i$ could be infected. Remove these from the infection history vector, $Z_{i,-\boldsymbol{z}}$

      ii. Count the length of $\boldsymbol{Z_{i,-z}}$ and the number of 1s in $\sum \boldsymbol{Z_{i,-z}}$ (this gives $m$ and $k$ as in the beta-Bernoulli respectively)

      iii. Iterate through 1 to $\boldsymbol{z}$ with index $j$:

         A. Calculate $p = \frac{\alpha+k}{\alpha+\beta+m}$

         B. Add a 1 at location $Z_{i,j}$ with probability $p$, add a 0 otherwise

         C. If a 1 was added, $k = k + 1$.

         D. $m = m + 1$

         E. Go to the next location in $\boldsymbol{z}$

3. Accept the proposed move with the acceptance ratio, noting that the proposal probability and prior $P(\boldsymbol{Z_i})$ cancel out:

$$A(\boldsymbol{Z_i'}, \boldsymbol{Z_i}) = \min(1, \frac{f(\mathbf{Y_i}|\boldsymbol{Z_i'}, \theta)}{f(\mathbf{Y_i}|\boldsymbol{Z_i}, \theta)}) \tag{63}$$

#### 5.4 Proposal algorithm under prior version 4, beta prior on overall probability of infection

The proposal algorithm for prior version 4 is identical to prior version 3, with only three changes:

1. In the step 2(a)(i)B, $k_j$ is replaced with $k_{-i,j} = (\sum_x \sum_y Z_{x,y}) - Z_{i,j}$
2. In step 2(a)(i)C, $n_j$ is replaced by $nm$
3. $P(Z)$ is the prior as described in Section 3.4

### References


[1] Lamnisos, Demetris, Griffin, Jim E. and Steel, Mark F.J. Adaptive Monte Carlo for Bayesian Variable Selection in Regression Models Journal of Computational and Graphical Statistics, 2013;22(3):729-748 DOI 10.1080/10618600.2012.694756

[2] George, Edward I. and McCulloch, Robert E. Approaches for Bayesian Variable Selection Statistica Sinica, 1997;7:339-373 DOI 10.2307/24306083

[3] O'Hara, R. B. and Sillanpää, M. J. A review of Bayesian variable selection methods: what, how and which Bayesian Analysis, 2009;4(1):85-117 DOI 10.1214/09-BA403

[4] Kerman, Jouni Neutral noninformative and informative conjugate beta and gamma prior distributions Electronic Journal of Statistics, 2011;5:1450-1470 DOI 10.1214/11-EJS648

[5] Griffiths, Thomas L. and Ghahramani, Zoubin The Indian Buffet Process: An Introduction and Review Journal of Machine Learning Research, 2011;12:1185-1224

[6] Butler, Ken and Stephens, Michael A. The Distribution of a Sum of Independent Binomial Random Variables Methodol Comput Appl Probab (2017) 19:557–571 DOI 10.1007/s11009-016-9533-4

[7] Liu, Boxiang and Quartermous, Thomas Approximating the Sum of Independent Non-Identical Binomial Random Variables The R Journal Vol. 10/1, July 2018 ISSN 2073-4859
