## Supplementary Material 2 for "Serosolver: an open source tool to infer epidemiological and immunological dynamics from serological data"

Supplementary Material 2: additional antibody kinetics

### Additional immunological mechanisms included in serosolver

#### Non-linear waning

As an alternative to the persistent long term and transient short term model described above, `serosolver` includes an option to have non-linear waning of the antibody response. Here, the long term boost ($\mu_1$) and the long-term cross-reactive response ($\sigma_1$) are both set to 0 and the single parameter that describes waning $\omega$ is replaced with the piece wise linear function,

$$\omega = \omega_1 t_m + \mathbb{1}_{(t_m > t_{change})} \omega_2 t_m \tag{1}$$

where $t_m = k - m$ is the time since infection with strain $m$, $\omega_2 = -\omega_1 \kappa$ and $\kappa \in (0, 1)$. The parameter $t_{change}$ indicates the time at which the slope of the antibody response changes following infection.

#### Additional boosting assumptions

`Serosolver` also includes optional mechanisms that account for titre-dependent and strain-dependent boosting. The strain-specific antibody boosting mechanism is included to allow for different levels of boosting for distinct strains or between clusters. In place of a global long-term boosting parameter, $\mu_1$, individual strain level or group level long-term boosting parameters can be passed into the model such that $\mu_1$ is replaced by $\mu_c$, where $c$ is the index of the boosting parameter for the infecting strain. The hierarchical structure of $\mu_c$ is flexible: each infecting strain may have a unique, independent $\mu_c$; different infection strains may have shared or separate $\mu_c$, for example within a single antigenic cluster; all $\mu_c$ may be estimated independently or drawn from a common distribution. In the latter case, the following term is added to the model:

$$\mu_c \sim \mathcal{N}(\bar{\mu_l}, \sigma^2_{\mu_l}) \tag{2}$$

where $\bar{\mu_l}$ is the mean long term boost, $\sigma_{\mu_l}$ is the standard deviation, both parameters to be estimated or fixed.

The titre-dependent boosting mechanism accounts for potential antibody ceiling effects, whereby antibody boosting is decreased at higher initial titres [1, 2]. For each infection, $m$, the predicted log antibody titre in Equation 2 in the main text is multiplied by $b(m)$,

$$b(m) = \begin{cases} \max\{0, 1 - \gamma x^m_{im}\} & \text{if } \; x^m_{im} \leq x_{change} \; ; \\ \max\{0, 1 - \gamma x_{change}\} & \text{else} \end{cases} \tag{3}$$

where $\gamma$ is a fitted parameter and $x_{change}$ is the titre-dependent boosting threshold. Initial titres above $x_{change}$ do not experience any further boosting suppression.

### Modifying serosolver's code to incorporate additional antibody kinetics models

Modifying source code can be challenging; however, it is our hope that `serosolver` will be extended and branched to test novel biological hypotheses. We have therefore structured the code with the hope that new mechanisms and assumptions can be added without major structural changes to the code base for a user with intermediate R and C++ programming experience. Table 1 outlines the locations in the source code that must be changed to modify the model called by `create_posterior_func`.

All of the plotting code, simulation code (eg. `simulate_data`), and create_posterior_func, ultimately calls titre_data_fast (L28 of src/infection_model_fast.cpp). Making the changes in Table 1 will therefore also update any post-processing, simulation code and the posterior function. Users should include additional model parameters in the parameter vector theta. For example, in example_par_tab, the long-term boosting parameter "mu" is described. This parameter is automatically located and then extracted at L78 of infection_model_fast.cpp. Note that the cross-reactivity matrices for long and short term antibody boosting, antigenic_map_long and antigenic_map_short are pre-calculated using create_cross_reactivity_vector. For example, at L172 and L337 in simulate_data.R, and L393 and L488 in posteriors.R. The rest of the arguments to titre_data_fast pass the current infection history matrix and vectors to control the titre data indexing.

Changes need to be made to titre_data_fast in 3 places: (i) extracting model parameters from L78; (ii) selecting the model version to use at L113; (iii) choosing the correct model to solve based on the options using an ifelse statement from L138. For computational speed reasons, the model solving code of titre_data_fast is directly integrated into the infection history proposal function for prior versions 2 and 4, inf_hist_prop_prior_v2_and_v4 at L163 of proposal.cpp. This code must therefore also be changed in the same way as for titre_data_fast. The equivalent 3 locations in proposal.cpp are: (i) L302; (ii) L339; (iii) L523.

### References


1. Jacobson RM, Grill DE, Oberg AL, Tosh PK, Ovsyannikova IG, Poland GA. Profiles of influenza A/H1N1 vaccine response using hemagglutination-inhibition titers. Human Vaccines & Immunotherapeutics. 2015;11(4):961–969. doi:10.1080/21645515.2015.1011990.

2. Petrie JG, Ohmit SE, Johnson E, Cross RT, Monto AS. Efficacy studies of influenza vaccines: effect of end points used and characteristics of vaccine failures. J Infect Dis. 2011;203(9):1309–1315. doi:10.1093/infdis/jir015.

**Table 1.** Summary of functions and files that must be changed to incorporate new antibody kinetics parameters/models.

| Purpose | Object/function | File | Note |
|---|---|---|---|
| Add new row to input parameter data frame for any new model parameters. Model parameters should be given (i) a name; (ii) a value; (iii) whether the parameter is fixed (1) or estimated (0); (iv) lower bound; (v) upper bound; (vi) lower bound for random starting value; (vii) upper bound for random starting value; (viii) parameter type (1 for kinetics parameter, 0 for a model option flag) | `par_tab` | Input object, specified in a file called `par_tab.csv` | NA |
| Add header for new antibody kinetics functions | NA | `src/boosting_functions_fast.h` | New functions added to `src/boosting_functions_fast.cpp` must first be defined here |
| Add new antibody kinetics model code | NA | `src/boosting_functions_fast.cpp` | Note that arguments to this function should take exactly the parameters needed for the model. The other function arguments (infection times, indices etc) should be taken from `titre_data_fast_individual_base` |
| Add calls to new antibody kinetics code | `titre_data_fast` and `inf_hist_prop_prior_v2_and_v4` | `src/infection_model_fast.cpp` and `src/proposal.cpp` | Setup for the function calls should be added to L78 and L113 for `titre_data_fast`, and L302 and L339 for `inf_hist_prop_prior_v2_and_v4` |
