## Supplementary Material 3 for "Serosolver: an open source tool to infer epidemiological and immunological dynamics from serological data"

Serosolver: paper case study 1


### Serosolver: paper case study 1

### Overview

This vignette provides all of the analyses for case study 1 in the accompanying package and paper. Briefly, the first case study aims to 1) reconstruct the unobserved infection dynamics from measured titres collected several months apart, 2) examine these infection dynamics stratified by available demographic variables, such as vaccination status and age, and 3) estimate biological parameters shaping the short-term antibody response.All of the functions used here are well documented and have many tunable arguments, and we therefore encourage users to refer to the helps files.

### Setup

#### Installation and requirements

`serosolver` may be installed from github using the `devtools` package. There are a number of additional packages that we need for this analysis.

```
# Required to run serosolver
#devtools::install_github("seroanalytics/serosolver")
library(serosolver)
library(plyr)
library(data.table)

## Required for this analysis
library(reshape2)
library(foreach)
library(doParallel)
library(bayesplot)
library(coda)
library(ggplot2)
library(viridis)

# set up cluster
set.seed(1234)
cl <- makeCluster(5)

## Note that this vignette was generated on a Windows machine,
## and the setup for parallelisation is different on a Linux or Mac:

if(Sys.info()[["sysname"]]=="Darwin" | Sys.info()[["sysname"]]=="Linux"){
  library(doMC)
  library(doRNG)
  registerDoMC(cores=5)
}else{
  registerDoParallel(cl)
}
```

#### Assumptions

In this analysis, serological samples were taken between 2009 and 2011 and therefore all time variables are relative to this time period. We are interested in inferring infections and attack rates at a quarterly resolution, and therefore set `resolution` to 4. Our primary outcome of interest is to infer unbiased attack rates, and we therefore use the version of the code with a Beta prior on per-time attack rates, `prior_version=2`. We set these parameters at the start of the analysis. Additionally, we assume that the samples are tested against the same virus, therefore, we set all antigenic coordinates (more information below) to those of the first time point.

```
filename <- "case_study_1"
resolution <- 4 ## set to 4 for quarterly resolution
sample_years <- 2009:2012

serosolver::describe_proposals()
#> Which version to use in run_MCMC? The following text describes the proposal step for updating infection histories.
#> Version 1: Beta prior on per time attack rates. Explicit FOI on each epoch using probability of infection term. Proposal performs N `flip` proposals at random locations in an individual's infection history, switching 1->0 or 0->1. Otherwise, swaps the contents of two random locations
#> Version 2: Beta prior on per time attack rates. Gibbs sampling of infection histories as in Indian Buffet Process papers, integrating out each probability of infection term.
#> Version 3: Beta prior on probability of infection for an individual, assuming independence between individuals. Samples from a beta binomial with alpha and beta specified by the par_tab input. Proposes nInfs moves at a time for add/remove, or when swapping, swaps locations up to moveSize time steps away
#> Version 4: Beta prior on probability of any infection. Gibbs sampling of infection histories using total number of infections across all times and all individuals as the prior
prior_version <- 2
```

#### Preparing the data

The data used in this analysis are haemagglutination inhibition (HI) titres against A/H1N1pdm09 that began circulating in 2009. The raw data have been pre-processed to both convert them into a form usable for `serosolver` and to separate the data into vaccinated and unvaccinated data sets. Details about the pre-processing can be found in the scripts folder of the `serosolver` package.

```
## Read in titre data
# unvaccinated
input_dat_path <- system.file("extdata", "HKdata_h1n1_unvac.csv", package = "serosolver")
input_dat <- read.csv(file = input_dat_path, header = TRUE)
# vaccinated
# input_dat_path2 <- system.file("extdata", "HKdata_h1n1_vac.csv", package = "serosolver")
# input_dat_vac <- read.csv(file = input_dat_path2, header = TRUE)

indivs <- unique(input_dat$individual) #all individuals

# Subset data for indivs
titre_dat <- input_dat[input_dat$individual %in% indivs,c("individual","virus","titre","samples","DOB")]
titre_dat$individual <- match(titre_dat$individual, indivs)

titre_dat <- unique(titre_dat)
titre_dat <- plyr::ddply(titre_dat,.(individual,virus,samples),function(x) cbind(x,"run"=1:nrow(x),"group"=1))
print(head(titre_dat))
#>   individual virus titre samples  DOB run group
#> 1          1  8037     0    8039 8036   1     1
#> 2          1  8037     0    8040 8036   1     1
#> 3          1  8037     7    8044 8036   1     1
#> 4          1  8037     7    8047 8036   1     1
#> 5          2  8037     0    8039 8036   1     1
#> 6          2  8037     5    8041 8036   1     1
```

NOTE: vaccinated and unvaccinated data must be run separately.

Given that this analysis uses titres from a single virus it is necessary to define an antigenic map with matching antigenic distance for each time point. We use coordinates based on the antigenic map created by Fonville et al. Generating the antigenic map involves fitting a smoothing spline through provided coordinates to give a representative virus for each time point (in this case, each quarter) that an individual could be infected. This process also inputs antigenic coordinates for time points that we do not have a measured virus. We then overwrite the coordinates at every time point with those of the first time point.

```
## Read in raw coordinates
antigenic_coords_path <- system.file("extdata", "fonville_map_approx.csv", package = "serosolver")
antigenic_coords <- read.csv(antigenic_coords_path, stringsAsFactors=FALSE)

## Convert to form expected by serosolver
antigenic_map <- generate_antigenic_map(antigenic_coords, resolution)

## Restrict entries to years of interest. Entries in antigenic_map determine
## the times that individual can be infected ie. the dimensions of the infection
## history matrix.
antigenic_map <- antigenic_map[antigenic_map$inf_years>=(sample_years[1]*resolution+1) & antigenic_map$inf_years<=sample_years[4]*resolution,]

## Change all coordinates to the same as the first pair of coordinates (because we assume the virus is the same at every time point)
antigenic_map[2:nrow(antigenic_map),c('x_coord','y_coord')] <- antigenic_map[1,c('x_coord','y_coord')]
print(head(antigenic_map))
#>      x_coord  y_coord inf_years
#> 166 37.69244 8.233511      8037
#> 167 37.69244 8.233511      8038
#> 168 37.69244 8.233511      8039
#> 169 37.69244 8.233511      8040
#> 170 37.69244 8.233511      8041
#> 171 37.69244 8.233511      8042

strain_isolation_times <- unique(antigenic_map$inf_years)
```

NOTE: `generate_antigenic_map` expects the provided file `fonville_map_approx.csv`. Users should refer to `generate_antigenic_map_flexible` for more generic antigenic map generation.

Finally, we must specify the `par_tab` data frame, which controls which parameters are included in the model, which are fixed, and their uniform prior ranges. Given that we are integrating out the probability of infection terms under prior version 2, we must remove these parameters from `par_tab`. Furthermore, given that we are interested in short-term dynamics with relatively sparse data, we remove parameters relating to the long-term antibody kinetics phase to avoid identifiability issues. We set alpha and beta to 1.

```
par_tab_path <- system.file("extdata", "par_tab_base.csv", package = "serosolver")
par_tab <- read.csv(par_tab_path, stringsAsFactors=FALSE)

## Set parameters for beta and alpha to 1
par_tab[par_tab$names %in% c("alpha","beta"),"values"] <- c(1,1)
## Maximum recordable log titre in these data is 9
par_tab[par_tab$names == "MAX_TITRE","values"] <- 9

## Remove phi parameters, as these are integrated out under prior version 2
par_tab <- par_tab[par_tab$names != "phi",]

## Fix all long term parameters to 0
par_tab[par_tab$names %in% c("mu","tau","sigma1","sigma2"),"fixed"] <- 1 #mu, tau, sigma1, and sigma2 are fixed
par_tab[par_tab$names %in% c("mu","tau","sigma1","sigma2"),"values"] <- 0 # set these values to 0
```

#### Summary

- Choose resolution, attack rate priors and reference time for “the present”
- Read in titre data and convert into a useable form for `serosolver`
- Generate antigenic map for the exposing virus
- Generate parameter control table for MCMC

### Running the MCMC

We are now ready to fit our model. We will fit multiple chains in parallel, though the below analysis could easily be replicated by running chains sequentially. Starting conditions for the MCMC chain must be generated that return a finite likelihood. The user may modify many of the MCMC control parameters, though the defaults are fine for most purposes.

Changing the number of iterations and the length of the adaptive period are often desirable. More crucially, the amount of chain thinning should be specified to ensure that users are not saving a large number of MCMC iterations (as this will rapidly fill disk space!). Thinning should be set such that at least 1000 iterations are saved (i.e., `iterations`/`thin` and `thin_hist`). Users are encouraged to pay extra attention to `thin_hist`, which dictates the thinning of the infection history chain, and can generate a very large file if left unchecked.

```
## Distinct filename for each chain
no_chains <- 5
filenames <- paste0(filename, "_",1:no_chains)
chain_path <- sub("par_tab_base.csv","",par_tab_path)
chain_path_real <- paste0(chain_path, "cs1_real/")
chain_path_sim <- paste0(chain_path, "cs1_sim/")

## Create the posterior solving function that will be used in the MCMC framework 
par_tab[par_tab$names == "mu_short","lower_bound"] <- 1
model_func <- create_posterior_func(par_tab=par_tab,
                            titre_dat=titre_dat,
                            antigenic_map=antigenic_map,
                            version=prior_version) # function in posteriors.R
#> Creating posterior solving function...
#> 
  
## Generate results in parallel
res <- foreach(x = filenames, .packages = c('serosolver','data.table','plyr')) %dopar% {
  ## Not all random starting conditions return finite likelihood, so for each chain generate random
  ## conditions until we get one with a finite likelihood
  start_prob <- -Inf
  while(!is.finite(start_prob)){
    ## Generating starting antibody kinetics parameters
    start_tab <- generate_start_tab(par_tab)
    
    ## Generate starting infection history
    start_inf <- setup_infection_histories_new_2(titre_dat, strain_isolation_times, space=3,titre_cutoff=4)
    start_prob <- sum(model_func(start_tab$values, start_inf)[[1]])
  }
  
  res <- run_MCMC(par_tab = start_tab, 
                  titre_dat = titre_dat,
                  antigenic_map = antigenic_map,
                  start_inf_hist = start_inf, 
                  mcmc_pars = c("iterations"=500000,"adaptive_period"=100000,
                                "thin"=1000,"thin_hist"=1000,
                                "save_block"=1000,"inf_propn"=1, 
                                "hist_sample_prob"=1,"hist_switch_prob"=0.8,
                                "year_swap_propn"=1),
                  filename = paste0(chain_path_real,x), 
                  CREATE_POSTERIOR_FUNC = create_posterior_func, 
                  version = prior_version)
}
```

### Post-run analyses

Once the MCMC chains are run, `serosolver` provides a number of simple functions to generate standard outputs and MCMC diagnostics. The saved MCMC chains are compatible with the `coda` and `bayesplot` packages, and users are encouraged to use these. First, read in the MCMC chains. The below function distinguishes between posterior samples for the infection history matrix and for the process parameters. The function searches for all files with the filenames generated by `run_MCMC` in the specified directory, and returns data structures with these concatenated and separated in a list.

```
## Read in the MCMC chains
# Note that `thin` here is in addition to any thinning done during the fitting
# Chain length values in load function need to be consistent with MCMC run
all_chains <- load_mcmc_chains(location=chain_path_real,thin=1,burnin=100000,
                             par_tab=par_tab,unfixed=FALSE,convert_mcmc=TRUE)
#> Chains detected:     5Highest MCMC sample interations: 
#> Chains detected: 
#> /tmp/Rtmpe5gKkm/temp_libpath6e556ded39fc/serosolver/extdata/cs1_real//case_study_1_1_infection_histories.csv
#> /tmp/Rtmpe5gKkm/temp_libpath6e556ded39fc/serosolver/extdata/cs1_real//case_study_1_2_infection_histories.csv
#> /tmp/Rtmpe5gKkm/temp_libpath6e556ded39fc/serosolver/extdata/cs1_real//case_study_1_3_infection_histories.csv
#> /tmp/Rtmpe5gKkm/temp_libpath6e556ded39fc/serosolver/extdata/cs1_real//case_study_1_4_infection_histories.csv
#> /tmp/Rtmpe5gKkm/temp_libpath6e556ded39fc/serosolver/extdata/cs1_real//case_study_1_5_infection_histories.csv
#> [[1]]
#> [1] 77636
#> 
#> [[2]]
#> [1] 77662
#> 
#> [[3]]
#> [1] 77610
#> 
#> [[4]]
#> [1] 77681
#> 
#> [[5]]
#> [1] 77747
## Alternative, load the included MCMC chains rather than re-running
## load(cs1_chains_real)
## all_chains <- cs1_chains_real

print(summary(all_chains))
#>                   Length Class      Mode   
#> theta_chain       65130  mcmc       numeric
#> inf_chain             5  data.table list   
#> theta_list_chains     5  -none-     list   
#> inf_list_chains       5  -none-     list
```

Chains should then be checked for the usual MCMC diagnostics: \(\hat{R}\) and effective sample size. First, looking at the antibody kinetics process parameters:

```
## Get the MCMC chains as a list
list_chains <- all_chains$theta_list_chains
## Look at diagnostics for the free parameters
list_chains1 <- lapply(list_chains, function(x) x[,c("mu_short","wane","error",
                                                     "total_infections",
                                                     "lnlike","prior_prob")])

## Gelman-Rubin diagnostics to assess between-chain convergence for each parameter
print(gelman.diag(as.mcmc.list(list_chains1)))
#> Potential scale reduction factors:
#> 
#>                  Point est. Upper C.I.
#> mu_short              1.001       1.01
#> wane                  1.002       1.01
#> error                 0.999       1.00
#> total_infections      1.003       1.01
#> lnlike                1.005       1.02
#> prior_prob            1.012       1.03
#> 
#> Multivariate psrf
#> 
#> 1.01
gelman.plot(as.mcmc.list(list_chains1))
```

```
## Effective sample size for each parameter
print(effectiveSize(as.mcmc.list(list_chains1)))
#>         mu_short             wane            error total_infections 
#>         1443.389         1559.938         2216.161         1670.971 
#>           lnlike       prior_prob 
#>         1043.881         1046.147

## Posterior estimates for each parameter
print(summary(as.mcmc.list(list_chains1)))
#> 
#> Iterations = 1:501
#> Thinning interval = 1 
#> Number of chains = 5 
#> Sample size per chain = 501 
#> 
#> 1. Empirical mean and standard deviation for each variable,
#>    plus standard error of the mean:
#> 
#>                        Mean        SD  Naive SE Time-series SE
#> mu_short          4.030e+00  0.098375 1.966e-03      0.0026774
#> wane              2.822e-02  0.004378 8.747e-05      0.0001122
#> error             7.885e-01  0.027680 5.531e-04      0.0005930
#> total_infections  1.550e+02  2.381665 4.759e-02      0.0620149
#> lnlike           -1.218e+03 10.026094 2.003e-01      0.3131179
#> prior_prob       -6.472e+02 11.108016 2.219e-01      0.3557723
#> 
#> 2. Quantiles for each variable:
#> 
#>                        2.5%        25%        50%        75%      97.5%
#> mu_short             3.8451  3.966e+00  4.026e+00  4.094e+00  4.227e+00
#> wane                 0.0196  2.547e-02  2.828e-02  3.084e-02  3.706e-02
#> error                0.7382  7.696e-01  7.870e-01  8.064e-01  8.468e-01
#> total_infections   150.0000  1.540e+02  1.550e+02  1.570e+02  1.590e+02
#> lnlike           -1238.1329 -1.225e+03 -1.218e+03 -1.212e+03 -1.199e+03
#> prior_prob        -667.0101 -6.547e+02 -6.479e+02 -6.403e+02 -6.237e+02


## Plot the MCMC trace using the `bayesplot` package
color_scheme_set("viridis")
p_theta_trace <- mcmc_trace(list_chains1)
print(p_theta_trace)
```

and at the infection histories:

```
## Extract infection history chain
inf_chain <- all_chains$inf_chain

## Look at inferred attack rates
p_ar <- plot_attack_rates(inf_chain, titre_dat, strain_isolation_times, pad_chain=FALSE,
                          plot_den = TRUE,prior_pars=list(prior_version=prior_version, 
                                                          alpha=par_tab[par_tab$names=="alpha","values"],
                                                          beta=par_tab[par_tab$names=="beta","values"])) 
print(p_ar)
```

```
## Calculate convergence diagnostics and summary statistics on infection histories
## Important to scale all infection estimates by number alive from titre_dat
n_alive <- get_n_alive_group(titre_dat, strain_isolation_times,melt=TRUE)

## This function generates a number of MCMC outputs
ps_infhist <- plot_posteriors_infhist(inf_chain=inf_chain, 
                                      years=strain_isolation_times, 
                                      samples = 100,  # Needs to be smaller than length of sampled chain 
                                      n_alive=n_alive)
#> Calculating by time summaries...
#> Done
#> Calculating by individual summaries...
#> Done


## Posterior mean, median, 95% credible intervals and effective sample size
## on per time attack rates
print(head(ps_infhist[["estimates"]]$by_year))
#> Empty data.table (0 rows and 7 cols): j,group,mean,median,lower_quantile,upper_quantile...

## Posterior mean, median, 95% credible intervals and effective sample size
## on per individual total number of infections
print(head(ps_infhist[["estimates"]]$by_indiv))
#>    i     mean median lower_quantile upper_quantile effective_size
#> 1: 1 2.000000      2              2              2          0.000
#> 2: 2 1.996806      2              2              2       2505.000
#> 3: 3 2.000000      2              2              2          0.000
#> 4: 4 1.969261      2              1              2       1954.678
#> 5: 6 1.000798      1              1              1       2505.000
#> 6: 7 1.913373      2              1              2       2505.000

## Check for agreement between inferred cumulative infection histories 
## for some individuals
p_indiv_inf_hists <- generate_cumulative_inf_plots(inf_chain, indivs=1:9, pad_chain=FALSE,
                                                   nsamp = 100, # Needs to be smaller than length of sampled chain 
                                                   strain_isolation_times = strain_isolation_times,
                                                   number_col=3)
print(p_indiv_inf_hists[[1]])
```

```
## Posterior probability that infections occured at given times per individual
print(p_indiv_inf_hists[[2]])
```

Users may also easily check the inferred antibody landscapes at the time each sample was taken. Black dots show observations, shaded regions and black line show 95%, 50% credible intervals and posterior median.

```
## get_titre_predictions expects only a single MCMC chain, so
## subset for only one chain
chain <- as.data.frame(all_chains$theta_chain)
chain1 <- chain[chain$chain_no == 1,]
inf_chain1 <- inf_chain[inf_chain$chain_no == 1,]

titre_preds <- get_titre_predictions(chain = chain1, 
                                     infection_histories = inf_chain1, 
                                     titre_dat = titre_dat, 
                                     individuals = unique(titre_dat$individual),
                                     nsamp = 100, # Needs to be smaller than length of sampled chain 
                                     antigenic_map = antigenic_map, 
                                     par_tab = par_tab,expand_titredat=FALSE)
#> Creating model solving function...
#> 
to_use <- titre_preds$predictions
print(head(to_use))
#>   individual virus titre samples  DOB run group    lower lower_50   median
#> 1          1  8037     0    8039 8036   1     1 0.000000 0.000000 0.000000
#> 2          1  8037     0    8040 8036   1     1 0.000000 0.000000 0.000000
#> 3          1  8037     7    8044 8036   1     1 7.340083 7.532602 7.683442
#> 4          1  8037     7    8047 8036   1     1 6.719864 6.878461 6.986778
#> 5          2  8037     0    8039 8036   1     1 0.000000 0.000000 0.000000
#> 6          2  8037     5    8041 8036   1     1 3.782653 3.907915 3.976667
#>   upper_50    upper      max
#> 1 0.000000 0.000000 0.000000
#> 2 0.000000 0.000000 0.000000
#> 3 7.769259 8.139377 7.904277
#> 4 7.098510 7.341082 7.524872
#> 5 0.000000 0.000000 0.000000
#> 6 4.035984 7.903366 4.015372

## Using ggplot
titre_pred_p <- ggplot(to_use[to_use$individual %in% c(1:8,13),])+
  geom_ribbon(aes(x=samples,ymin=lower, ymax=upper),fill="gray90")+
  geom_ribbon(aes(x=samples,ymin=lower_50, ymax=upper_50),fill="gray70")+
  geom_line(aes(x=samples, y=median))+
  geom_point(aes(x=samples, y=titre))+
  coord_cartesian(ylim=c(0,8))+
  ylab("log titre") +
  xlab("Time of virus circulation") +
  theme_classic() +
  facet_wrap(~individual)
titre_pred_p
```

### Simulation recovery

We finish the vignette by presenting a simulation-recovery experiment to test the ability of the framework to recover known infection histories and antibody kinetics parameters using simulated data that matches the real dataset.

#### Extract attack rates from fits

We simulate infection histories and antibody titre data based on the “real” parameters inferred from fitting the model above. First, we extract the maximum posterior probability antibody kinetics parameters and attack rates.

```
## Read in MCMC chains from fitting
all_chains <- load_mcmc_chains(location=chain_path_real,thin=1,burnin=100000,
                               par_tab=par_tab,unfixed=FALSE,convert_mcmc=FALSE)
#> Chains detected:     5Highest MCMC sample interations: 
#> Chains detected: 
#> /tmp/Rtmpe5gKkm/temp_libpath6e556ded39fc/serosolver/extdata/cs1_real//case_study_1_1_infection_histories.csv
#> /tmp/Rtmpe5gKkm/temp_libpath6e556ded39fc/serosolver/extdata/cs1_real//case_study_1_2_infection_histories.csv
#> /tmp/Rtmpe5gKkm/temp_libpath6e556ded39fc/serosolver/extdata/cs1_real//case_study_1_3_infection_histories.csv
#> /tmp/Rtmpe5gKkm/temp_libpath6e556ded39fc/serosolver/extdata/cs1_real//case_study_1_4_infection_histories.csv
#> /tmp/Rtmpe5gKkm/temp_libpath6e556ded39fc/serosolver/extdata/cs1_real//case_study_1_5_infection_histories.csv
#> [[1]]
#> [1] 77636
#> 
#> [[2]]
#> [1] 77662
#> 
#> [[3]]
#> [1] 77610
#> 
#> [[4]]
#> [1] 77681
#> 
#> [[5]]
#> [1] 77747

## Alternative, load the included MCMC chains rather than re-running
## load(cs1_chains_real_b)
## all_chains <- cs1_chains_real_b

## Find samples that were in both theta and inf hist chains
chain <- all_chains$theta_chain
inf_chain <- all_chains$inf_chain
intersect_samps <- intersect(unique(inf_chain$sampno), unique(chain$sampno))
chain <- chain[chain$sampno %in% intersect_samps,]

## Find the parameter values that gave the highest posterior probability
which_mle <- chain[which.max(chain$lnlike),c("sampno","chain_no")]
mle_theta_pars <- chain[chain$sampno == which_mle$sampno & chain$chain_no == which_mle$chain_no,]

## Store total infections to compare later
mle_total_infs <- mle_theta_pars[,"total_infections"]
mle_theta_pars <- mle_theta_pars[,par_tab$names]
mle_inf_hist <- inf_chain[inf_chain$sampno == which_mle$sampno & inf_chain$chain_no == which_mle$chain_no,]

## Generate full infection history matrix using provided function
mle_inf_hist <- expand_summary_inf_chain(mle_inf_hist[,c("sampno","j","i","x")])
## Find number of infections per year from this infection history
no_infs <- colSums(mle_inf_hist[,3:ncol(mle_inf_hist)])

## If missing time points in simulated attack rates
if(length(no_infs) < length(strain_isolation_times)){
  diff_lengths <- length(strain_isolation_times) - length(no_infs)
  no_infs <- c(no_infs, rep(0, diff_lengths))
}

## Find attack rate per year
n_alive <- get_n_alive(titre_dat, strain_isolation_times)
attack_rates <- no_infs/n_alive
```

Functions are provided to simulate antibody titre data under a given serosurvey design. The antibody kinetics parameters and attack rates estimated above are used to simulate titres from the model. The `simulate_data` function is well documented, and users should refer to the help file to customise the simulated serosurvey design.

```
set.seed(1234)

sim_par_tab <- par_tab
sim_par_tab$values <- as.numeric(mle_theta_pars)
sim_par_tab[sim_par_tab$names %in% c("alpha","beta"),"values"] <- c(1/3,1/3)
sim_par_tab[sim_par_tab$names == "wane","values"] <- 1
sim_par_tab[sim_par_tab$names == "wane","values"] <- par_tab[par_tab$names == "wane","values"]/resolution
sim_par_tab[sim_par_tab$names %in% c("mu","tau","sigma1","sigma2"),"fixed"] <- 1 
sim_par_tab[sim_par_tab$names %in% c("mu","tau","sigma1","sigma2"),"values"] <- 0 
sim_par_tab[sim_par_tab$names == "MAX_TITRE","values"] <- 9 

sampling_times <- seq(2009*resolution + 1, 2012*resolution, by=1)

age_min <- 6*resolution
age_max <- 6*resolution
n_indiv <- length(unique(titre_dat$individual))

dat <- simulate_data(par_tab = sim_par_tab, 
                     n_indiv = n_indiv,
                     buckets = resolution,
                     strain_isolation_times = strain_isolation_times,
                     sampling_times = sampling_times, 
                     nsamps = 4, 
                     antigenic_map = antigenic_map, 
                     age_min = age_min,
                     age_max = age_max,
                     attack_rates=attack_rates,
                     repeats = 1)
#> Simulating data


## Inspect simulated antibody titre data and infection histories
sim_titre_dat <- dat[["data"]]
sim_infection_histories <- dat[["infection_histories"]]

## Store total infections to compare later
actual_total_infections <- sum(sim_infection_histories)

plot_data(sim_titre_dat, sim_infection_histories, strain_isolation_times,n_samps = 5)
```

```
## Use titres only against same viruses tested in real data
viruses <- unique(titre_dat$virus)
sim_titre_dat <- sim_titre_dat[sim_titre_dat$virus %in% viruses, ]
sim_ages <- dat[["ages"]]
sim_titre_dat <- merge(sim_titre_dat, sim_ages)
sim_ar <- dat[["attack_rates"]]
```

#### Simulation fitting

Once these simulated data have been generated, the work flow becomes exactly the same as with the real data above.

```
filename <- "case_study_1_sim"

## Distinct filename for each chain
no_chains <- 5
filenames <- paste0(filename, "_",1:no_chains)

## Create the posterior solving function that will be used in the MCMC framework 
model_func <- create_posterior_func(par_tab=sim_par_tab,
                                titre_dat=sim_titre_dat,
                                antigenic_map=antigenic_map,
                                version=2) # function in posteriors.R
#> Creating posterior solving function...
#> 

## Generate results in parallel
res <- foreach(x = filenames, .packages = c('serosolver','data.table','plyr')) %dopar% {
  ## Not all random starting conditions return finite likelihood, so for each chain generate random
  ## conditions until we get one with a finite likelihood
  start_prob <- -Inf
  while(!is.finite(start_prob)){
    ## Generate starting values for theta
    start_tab <- generate_start_tab(par_tab)
    ## Generate starting infection history
    start_inf <- setup_infection_histories_new_2(sim_titre_dat, strain_isolation_times, space=3,titre_cutoff=4)
    start_prob <- sum(model_func(start_tab$values, start_inf)[[1]])
  }
  
  res <- run_MCMC(par_tab = start_tab, 
                  titre_dat = sim_titre_dat,
                  antigenic_map = antigenic_map,
                  start_inf_hist = start_inf, 
                  mcmc_pars = c("iterations"=500000,"adaptive_period"=100000,"thin"=1000,
                                "thin_hist"=1000,"save_block"=1000,
                                "hist_switch_prob"=0.8, "hist_sample_prob"=1),
                  filename = paste0(chain_path_sim,x), 
                  CREATE_POSTERIOR_FUNC = create_posterior_func, 
                  version = prior_version)
}
```

#### Simulation analysis

MCMC chains should be checked for convergence under the usual diagnostics. We also compare the inferred posterior distributions to the known true parameter values. We see that convergence and between-chain agreement is good and that the model recovers reasonably unbiased estimates for some parameters. However, under this sampling strategy the model slightly underestimates the amount of long term antibody boosting elicited by a single infection and overestimates the total number of infections. This is driven by the contribution of the attack rate prior relative to the contribution of the likelihood (the data). Increasing the number of measured titres (for example, measure titres against 40 viruses rather than 9) or using a more informative attack rate prior would help reduce this bias.

```
## Read in the MCMC chains
## Note that `thin` here is in addition to any thinning done during the fitting
sim_all_chains <- load_mcmc_chains(location=chain_path_sim,thin=1,burnin=100000,
                                   par_tab=par_tab,unfixed=FALSE,convert_mcmc=TRUE)
#> Chains detected:     5Highest MCMC sample interations: 
#> Chains detected: 
#> /tmp/Rtmpe5gKkm/temp_libpath6e556ded39fc/serosolver/extdata/cs1_sim//case_study_1_sim_1_infection_histories.csv
#> /tmp/Rtmpe5gKkm/temp_libpath6e556ded39fc/serosolver/extdata/cs1_sim//case_study_1_sim_2_infection_histories.csv
#> /tmp/Rtmpe5gKkm/temp_libpath6e556ded39fc/serosolver/extdata/cs1_sim//case_study_1_sim_3_infection_histories.csv
#> /tmp/Rtmpe5gKkm/temp_libpath6e556ded39fc/serosolver/extdata/cs1_sim//case_study_1_sim_4_infection_histories.csv
#> /tmp/Rtmpe5gKkm/temp_libpath6e556ded39fc/serosolver/extdata/cs1_sim//case_study_1_sim_5_infection_histories.csv
#> [[1]]
#> [1] 71411
#> 
#> [[2]]
#> [1] 71652
#> 
#> [[3]]
#> [1] 71613
#> 
#> [[4]]
#> [1] 71506
#> 
#> [[5]]
#> [1] 71460

## Alternative, load the included MCMC chains rather than re-running
## load(cs1_chains_sim)
## sim_all_chains <- cs1_chains_sim

theta_chain <- sim_all_chains$theta_chain
## Get the MCMC chains as a list
list_chains <- sim_all_chains$theta_list_chains
## Look at diagnostics for the free parameters
list_chains1 <- lapply(list_chains, function(x) x[,c("mu_short","wane","error",
                                                     "total_infections",
                                                     "lnlike","prior_prob")])

## Gelman-Rubin diagnostics and effective sample size
print(gelman.diag(as.mcmc.list(list_chains1)))
#> Potential scale reduction factors:
#> 
#>                  Point est. Upper C.I.
#> mu_short              0.999       1.00
#> wane                  1.001       1.00
#> error                 1.004       1.01
#> total_infections      1.007       1.02
#> lnlike                1.018       1.05
#> prior_prob            1.018       1.05
#> 
#> Multivariate psrf
#> 
#> 1.02
print(effectiveSize(as.mcmc.list(list_chains1)))
#>         mu_short             wane            error total_infections 
#>        2243.4232        1995.0457        2328.1566        1053.4466 
#>           lnlike       prior_prob 
#>         826.3437         737.4148

melted_theta_chain <- reshape2::melt(as.data.frame(theta_chain), id.vars=c("sampno","chain_no"))
estimated_pars <- c(sim_par_tab[sim_par_tab$fixed == 0,"names"],"total_infections")
melted_theta_chain <- melted_theta_chain[melted_theta_chain$variable %in% estimated_pars,]
colnames(melted_theta_chain)[3] <- "names"

add_row <- data.frame("total_infections",actual_total_infections,0,0.1,0,10000,0,0,1)
colnames(add_row) <- colnames(sim_par_tab)
sim_par_tab1 <- rbind(sim_par_tab, add_row)

ggplot(melted_theta_chain) + 
  geom_density(aes(x=value,fill=as.factor(chain_no)),alpha=0.5) +
  geom_vline(data=sim_par_tab1[sim_par_tab1$fixed == 0,],aes(xintercept=values),linetype="dashed") +
  facet_wrap(~names,scales="free") + 
  theme_classic() +
  theme(legend.position="bottom")
```

Recovery of known attack rates is also reasonably accurate, though the constraint of the posterior distibution is quite low for many years where identifiability is poor. Again, more titre data or more individuals would improve inferential power. One particularly reassuring plot is the comparison of known individual cumulative infection histories (the cumulative sum of infections over time for an individual) against the estimated posterior distribution of cumulative infection histories. We see that the 95% credible intervals capture the true cumulative infection histories in almost all cases.

```
## Extract infection history chain
inf_chain <- sim_all_chains$inf_chain

## Look at inferred attack rates
p_ar <- plot_attack_rates(inf_chain, sim_titre_dat, strain_isolation_times, pad_chain=FALSE,
                            plot_den = TRUE,prior_pars=list(prior_version=prior_version, 
                                                          alpha=par_tab[par_tab$names=="alpha","values"],
                                                          beta=par_tab[par_tab$names=="beta","values"]))  +
  geom_point(data=sim_ar,aes(x=year,y=AR),col="purple")

print(p_ar)
```

```
## Calculate convergence diagnostics and summary statistics on infection histories
## Important to scale all infection estimates by number alive from titre_dat
sim_n_alive <- get_n_alive_group(sim_titre_dat, strain_isolation_times,melt=TRUE)

## This function generates a number of MCMC outputs
ps_infhist <- plot_posteriors_infhist(inf_chain=inf_chain, 
                                      years=strain_isolation_times, 
                                      n_alive=sim_n_alive)
#> Calculating by time summaries...
#> Done
#> Calculating by individual summaries...
#> Done

## Check for agreement between inferred cumulative infection histories 
## for some individuals
p_indiv_inf_hists <- generate_cumulative_inf_plots(inf_chain,indivs=c(12,21,39,41,44,66,71,76,79),pad_chain=FALSE,
                                                   real_inf_hist=sim_infection_histories,
                                                  strain_isolation_times = strain_isolation_times,
                                                  number_col=3)
print(p_indiv_inf_hists[[1]])
```

```
## Posterior probability that infections occured at given times per individual
print(p_indiv_inf_hists[[2]])
```
