## Supplementary Material 4 for "Serosolver: an open source tool to infer epidemiological and immunological dynamics from serological data"

Serosolver: paper case study 2


### Serosolver: paper case study 2

### Overview

This vignette provides all of the analysis for case study 2 in the accompanying package and paper. Briefly, the aim is to infer antibody kinetics and historical attack rates using cross-sectional haemagglutination inhibition titre data on a panel of recent and historical A/H3N2 strains. All of the functions used here are well documented and have many tunable arguments, and we therefore encourage users to refer to the helps files.

### Setup

#### Installation and requirements

`serosolver` may be installed from github using the `devtools` package. There are a number of additional packages that we need for this analysis.

```
# Required to run serosolver
devtools::install_github("seroanalytics/serosolver")
#> 
#>   
   checking for file ‘/tmp/Rtmpe5gKkm/remotes6e553f4d9b65/seroanalytics-serosolver-1d3bb37/DESCRIPTION’ ...
  
✔  checking for file ‘/tmp/Rtmpe5gKkm/remotes6e553f4d9b65/seroanalytics-serosolver-1d3bb37/DESCRIPTION’
#> 
  
─  preparing ‘serosolver’:
#> 
  
   checking DESCRIPTION meta-information ...
  
✔  checking DESCRIPTION meta-information
#> ─  cleaning src
#> 
  
─  checking for LF line-endings in source and make files and shell scripts
#> 
  
─  checking for empty or unneeded directories
#> 
  
─  looking to see if a ‘data/datalist’ file should be added
#> 
  
─  building ‘serosolver_1.0.0.tar.gz’
#> 
  
   
#> 
library(serosolver)
library(plyr)
library(data.table)

## Required for this analysis
library(reshape2)
library(foreach)
library(doParallel)
library(bayesplot)
library(coda)
library(ggplot2)
library(viridis)

# set up cluster
set.seed(1234)
cl <- makeCluster(5)
registerDoParallel(cl)

## Note that this vignette was generated on a Windows machine,
## and the setup for parallelisation is different on a Linux machine
## for Linux machine:
#  library(doMC)
#  library(doRNG)
#  registerDoMC(cores=5)
```

#### Assumptions

In this analysis, all serological samples were taken in 2009 and therefore all time variables are relative to this year. We are interested in inferring infections and attack rates at an annual resolution, and therefore set `resolution` to 1. Our primary outcome of interest is to infer unbiased historical attack rates, and we therefore use the version of the code with a Beta prior on per-time attack rates, `prior_version=2`. Furthermore, we have found that in the situation where the number of possible infections to infer is large but the amount of data is relatively sparse, identifiability is poor when using reference priors (eg. uniform Beta or Jeffrey’s prior). We instead opted to use a weakly informative prior for annual attack rates with mode = 0.15 but with high variance, corresponding to prior observations of annual influenza attack rates XX REF XX. We set these parameters at the start of the analysis.

```
filename <- "case_study_2"
resolution <- 1 ## eg. this would be set to 12 for monthly resolution
sample_year <- 2009

serosolver::describe_proposals()
#> Which version to use in run_MCMC? The following text describes the proposal step for updating infection histories.
#> Version 1: Beta prior on per time attack rates. Explicit FOI on each epoch using probability of infection term. Proposal performs N `flip` proposals at random locations in an individual's infection history, switching 1->0 or 0->1. Otherwise, swaps the contents of two random locations
#> Version 2: Beta prior on per time attack rates. Gibbs sampling of infection histories as in Indian Buffet Process papers, integrating out each probability of infection term.
#> Version 3: Beta prior on probability of infection for an individual, assuming independence between individuals. Samples from a beta binomial with alpha and beta specified by the par_tab input. Proposes nInfs moves at a time for add/remove, or when swapping, swaps locations up to moveSize time steps away
#> Version 4: Beta prior on probability of any infection. Gibbs sampling of infection histories using total number of infections across all times and all individuals as the prior
prior_version <- 2
```

#### Preparing the data

The data used in this analysis are haemagglutination inhibition (HI) titres against a number of A/H3N2 that have circulated since 1968. The raw data are in a wide format, providing the highest two-fold dilution of serum at which haemagglutination is inhibited. The first step of the analysis is therefore to clean the titre data and convert the data frame to the long format, as described in the quickstart vignette.

```
## Read in data
raw_dat_path <- system.file("extdata", "Fluscape_HI_data.csv", package = "serosolver")
raw_dat <- read.csv(file = raw_dat_path, stringsAsFactors = FALSE)
print(head(raw_dat))
#>   Age HI.H3N2.1968 HI.H3N2.1975 HI.H3N2.1979 HI.H3N2.1989 HI.H3N2.1995
#> 1  75           80           40           40           80          160
#> 2  35           20           80          160           40           80
#> 3  71           80           40           20           20           40
#> 4  65           80           40           40           20           40
#> 5  64          160           80           40           10           40
#> 6  33           40           20          160           80           80
#>   HI.H3N2.2002 HI.H3N2.2003 HI.H3N2.2005 HI.H3N2.2008
#> 1          160           40           80           40
#> 2           20           10            0            0
#> 3           80           20           10            0
#> 4           20            0            0            0
#> 5           40            0           20           20
#> 6          160           40           40           20

## Add indexing column for each individual
raw_dat$individual <- 1:nrow(raw_dat)

## Convert data to long format
melted_dat <- reshape2::melt(raw_dat, id.vars=c("individual","Age"),stringsAsFactors=FALSE)

## Modify column names to meet serosolver's expectations
colnames(melted_dat) <- c("individual","DOB","virus","titre")
melted_dat$virus <- as.character(melted_dat$virus)

## Extract circulation years for each virus code, which will be used 
## by serosolver as the circulation time
melted_dat$virus <- as.numeric(sapply(melted_dat$virus, function(x) strsplit(x,split = "HI.H3N2.")[[1]][2]))

## Clean and log transform the data
melted_dat <- melted_dat[complete.cases(melted_dat),]
melted_dat[melted_dat$titre == 0,"titre"] <- 5
melted_dat$titre <- log2(melted_dat$titre/5)

## Convert ages to DOB
melted_dat$DOB <- sample_year - melted_dat$DOB

## All samples taken at the same time
melted_dat$samples <- sample_year

## Add column for titre repeats, enumerating for each measurement for the same virus/sample/individual
melted_dat <- plyr::ddply(melted_dat,.(individual,virus,samples),function(x) cbind(x,"run"=1:nrow(x),"group"=1))

## Rename to data expected by serosolver
titre_dat <- melted_dat
print(head(titre_dat))
#>   individual  DOB virus titre samples run group
#> 1          1 1934  1968     4    2009   1     1
#> 2          1 1934  1975     3    2009   1     1
#> 3          1 1934  1979     3    2009   1     1
#> 4          1 1934  1989     4    2009   1     1
#> 5          1 1934  1995     5    2009   1     1
#> 6          1 1934  2002     5    2009   1     1
```

Given that this analysis uses titres from multiple, antigenically related viruses, it is necessary to define an antigenic map describing the antigenic distance between all of the viruses here. We use coordinates based on the antigenic map created by Fonville et al. Generating the antigenic map involves fitting a smoothing spline through provided coordinates to give a representative virus for each time point (in this case, each year) that an individual could be infected. This process also inputs antigenic coordinates for time points that we do not have a measured virus.

```
## Read in raw coordinates
antigenic_coords_path <- system.file("extdata", "fonville_map_approx.csv", package = "serosolver")
antigenic_coords <- read.csv(file = antigenic_coords_path, stringsAsFactors=FALSE)
print(head(antigenic_coords))
#>   Strain    X    Y
#> 1   HK68  1.8  2.4
#> 2   EN72  2.7  4.9
#> 3   VI75  7.6  6.3
#> 4   TX77  7.9  8.8
#> 5   BK79  9.6 11.0
#> 6   SI87 15.0  6.8

## Convert to form expected by serosolver
antigenic_map <- generate_antigenic_map(antigenic_coords, resolution)
print(head(antigenic_map))
#>      x_coord   y_coord inf_years
#> 1 -0.3036008 0.2322615      1968
#> 2  0.6175154 1.4365363      1969
#> 3  1.5386317 2.6408110      1970
#> 4  2.4597479 3.8217916      1971
#> 5  3.3808642 4.7622756      1972
#> 6  4.3019805 5.4353631      1973

## More flexible version of the above function
virus_key <- c(
    "HK68" = 1968, "EN72" = 1972, "VI75" = 1975, "TX77" = 1977, "BK79" = 1979, "SI87" = 1987, "BE89" = 1989, "BJ89" = 1989,
    "BE92" = 1992, "WU95" = 1995, "SY97" = 1997, "FU02" = 2002, "CA04" = 2004, "WI05" = 2005, "PE06" = 2006
  )
antigenic_coords$Strain <- virus_key[antigenic_coords$Strain]
antigenic_map <- generate_antigenic_map_flexible(antigenic_coords)

## Restrict entries to years of interest. Entries in antigenic_map determine
## the times that individual can be infected ie. the dimensions of the infection
## history matrix.
antigenic_map <- antigenic_map[antigenic_map$inf_years >= 1968 & antigenic_map$inf_years <= sample_year,]
strain_isolation_times <- unique(antigenic_map$inf_years)
```

NOTE: `generate_antigenic_map` expects the provided file `fonville_map_approx.csv`. Users should refer to `generate_antigenic_map_flexible` for more generic antigenic map generation.

Finally, we must specify the `par_tab` data frame, which controls which parameters are included in the model, which are fixed, and their uniform prior ranges. Given that we are integrating out the probability of infection terms under prior version 2, we must remove these parameters from `par_tab`. Furthermore, given that we are interested in long-term dynamics with relatively sparse data, we remove parameters relating to the short-term antibody kinetics phase to avoid identifiability issues. We set alpha and beta of the Beta prior to give a mode of 0.15 assuming that our prior belief has the equivalent weighting to 4 observed individuals.

```
par_tab_path <- system.file("extdata", "par_tab_base.csv", package = "serosolver")
par_tab <- read.csv(file = par_tab_path, stringsAsFactors=FALSE)

## Set parameters for Beta prior on infection histories
beta_pars <- find_beta_prior_mode(0.15,4)
par_tab[par_tab$names == "alpha","values"] <- beta_pars$alpha
par_tab[par_tab$names == "beta","values"] <- beta_pars$beta
## Maximum recordable log titre in these data is 8
par_tab[par_tab$names == "MAX_TITRE","values"] <- 8

## Remove phi parameters, as these are integrated out under prior version 2
par_tab <- par_tab[par_tab$names != "phi",]

## Fix all short term parameters to 0
par_tab[par_tab$names %in% c("mu_short","sigma2","wane"),"fixed"] <- 1 # mu_short, waning and sigma2 are fixed
par_tab[par_tab$names %in% c("mu_short","sigma2","wane"),"values"] <- 0 # set these values to 0
```

#### Summary

### Running the MCMC

We are now ready to fit our model. We will fit multiple chains in parallel, though the below analysis could easily be replicated by running chains sequentially. Starting conditions for the MCMC chain must be generated that return a finite likelihood. The user may modify many of the MCMC control parameters, though the defaults are fine for most purposes. We have made some minor tweaks in this case study to improve convergence on infection history estimates. Step sizes for parameters in `par_tab` are tuned automatically, and some automated tuning of the infection history proposals takes place for prior version 3. However, for other attack rate priors, it is necessary for the user to do some manual tuning of a) the number of individuals sampled at each step `hist_sample_prob`; b) the number of time points sampled at each step `inf_propn`; c) the frequency of individual infection history swapping steps (ie. for an individual, choose two time points and swap their contents)`swap_propn`; d) proportion of infection history sampling steps which should be the alternative swapping step, where the contents of infection histories at two time points are swapped `hist_switch_prob`; e) proportion of infection histories to swap with each alternative swapping step `year_swap_propn`. For example, in this case study, we attack rates are likely to be highly correlated in adjacent years (as we have limited data to distinguish between infections in years close in time), and we therefore increase the frequency of the alternative infection history swapping step with `year_swap_propn`.

Changing the number of iterations and the length of the adaptive period are often desirable. More crucially, the amount of chain thinning should be specified to ensure that users are not saving a large number of MCMC iterations (as this will rapidly fill disk space!). Thinning should be set such that at least 1000 iterations are saved (ie. `iterations`/`thin` and `thin_hist`). Users are encouraged to pay extra attention to `thin_hist`, which dictates the thinning of the infection history chain, and can generate a very large file if left unchecked.

```
## Distinct filename for each chain
no_chains <- 5
filenames <- paste0(filename, "_",1:no_chains)
chain_path <- sub("par_tab_base.csv","",par_tab_path)
chain_path_real <- paste0(chain_path, "cs2_real/")
chain_path_sim <- paste0(chain_path, "cs2_sim/")

## Create the posterior solving function that will be used in the MCMC framework 
model_func <- create_posterior_func(par_tab=par_tab,
                                titre_dat=titre_dat,
                                antigenic_map=antigenic_map,
                                version=prior_version) # function in posteriors.R
#> Creating posterior solving function...
#> 

## Generate results in parallel
res <- foreach(x = filenames, .packages = c('serosolver','data.table','plyr')) %dopar% {
  ## Not all random starting conditions return finite likelihood, so for each chain generate random
  ## conditions until we get one with a finite likelihood
  start_prob <- -Inf
  while(!is.finite(start_prob)){
    ## Generating starting antibody kinetics parameters
    start_tab <- generate_start_tab(par_tab)
    
    ## Generate starting infection history
    start_inf <- setup_infection_histories_new_2(titre_dat, strain_isolation_times, space=3,titre_cutoff=4)
    start_prob <- sum(model_func(start_tab$values, start_inf)[[1]])
  }
  
  
  res <- run_MCMC(par_tab = start_tab, 
                  titre_dat = titre_dat,
                  antigenic_map = antigenic_map,
                  start_inf_hist = start_inf, 
                  mcmc_pars = c("iterations"=500000,"adaptive_period"=100000,"thin"=1000,
                                "thin_hist"=1000,"save_block"=1000,
                                "inf_propn"=1, "hist_sample_prob"=1,
                                "hist_switch_prob"=0.8, "year_swap_propn"=1),
                  filename = paste0(chain_path_real,x), 
                  CREATE_POSTERIOR_FUNC = create_posterior_func, 
                  version = prior_version)
}
```

### Post-run analyses

Once the MCMC chains are run, `serosolver` provides a number of simple functions to generate standard outputs and MCMC diagnostics. The saved MCMC chains are compatible with the `coda` and `bayesplot` packages, and users are encouraged to use these. First, read in the MCMC chains. The below function distinguishes between posterior samples for the infection history matrix and for the process parameters. The function searches for all files with the filenames generated by `run_MCMC` in the specified directory, and returns data structures with these concatenated and seperated in a list.

```
## Read in the MCMC chains
## Note that `thin` here is in addition to any thinning done during the fitting
all_chains <- load_mcmc_chains(location=chain_path_real,thin=1,burnin=100000,
                               par_tab=par_tab,unfixed=FALSE,convert_mcmc=TRUE)
#> Chains detected:     5Highest MCMC sample interations: 
#> Chains detected: 
#> /tmp/Rtmpe5gKkm/temp_libpath6e556ded39fc/serosolver/extdata/cs2_real//case_study_2_1_infection_histories.csv
#> /tmp/Rtmpe5gKkm/temp_libpath6e556ded39fc/serosolver/extdata/cs2_real//case_study_2_2_infection_histories.csv
#> /tmp/Rtmpe5gKkm/temp_libpath6e556ded39fc/serosolver/extdata/cs2_real//case_study_2_3_infection_histories.csv
#> /tmp/Rtmpe5gKkm/temp_libpath6e556ded39fc/serosolver/extdata/cs2_real//case_study_2_4_infection_histories.csv
#> /tmp/Rtmpe5gKkm/temp_libpath6e556ded39fc/serosolver/extdata/cs2_real//case_study_2_5_infection_histories.csv
#> [[1]]
#> [1] 636633
#> 
#> [[2]]
#> [1] 645305
#> 
#> [[3]]
#> [1] 653492
#> 
#> [[4]]
#> [1] 639808
#> 
#> [[5]]
#> [1] 646868

## Alternative, load the included MCMC chains rather than re-running
## load(cs2_chains_real)
## all_chains <- cs2_chains_real

print(summary(all_chains))
#>                   Length Class      Mode   
#> theta_chain       65130  mcmc       numeric
#> inf_chain             5  data.table list   
#> theta_list_chains     5  -none-     list   
#> inf_list_chains       5  -none-     list
```

Chains should then be checked for the usual MCMC diagnostics: \(\hat{R}\) and effective sample size. First, looking at the antibody kinetics process parameters:

```
## Get the MCMC chains as a list
list_chains <- all_chains$theta_list_chains
## Look at diagnostics for the free parameters
list_chains1 <- lapply(list_chains, function(x) x[,c("mu","sigma1","error",
                                                     "tau","total_infections",
                                                     "lnlike","prior_prob")])

## Gelman-Rubin diagnostics to assess between-chain convergence for each parameter
print(gelman.diag(as.mcmc.list(list_chains1)))
#> Potential scale reduction factors:
#> 
#>                  Point est. Upper C.I.
#> mu                     1.03       1.08
#> sigma1                 1.01       1.02
#> error                  1.00       1.00
#> tau                    1.01       1.03
#> total_infections       1.02       1.05
#> lnlike                 1.03       1.08
#> prior_prob             1.03       1.08
#> 
#> Multivariate psrf
#> 
#> 1.03
gelman.plot(as.mcmc.list(list_chains1))
```

```
## Effective sample size for each parameter
print(effectiveSize(as.mcmc.list(list_chains1)))
#>               mu           sigma1            error              tau 
#>         375.6031         658.9961        2335.9568        1100.0764 
#> total_infections           lnlike       prior_prob 
#>         549.9216         551.4273         553.4807

## Posterior estimates for each parameter
print(summary(as.mcmc.list(list_chains1)))
#> 
#> Iterations = 1:501
#> Thinning interval = 1 
#> Number of chains = 5 
#> Sample size per chain = 501 
#> 
#> 1. Empirical mean and standard deviation for each variable,
#>    plus standard error of the mean:
#> 
#>                        Mean        SD  Naive SE Time-series SE
#> mu                2.222e+00 1.409e-01 2.814e-03      0.0075051
#> sigma1            1.044e-01 4.362e-03 8.715e-05      0.0001721
#> error             1.162e+00 3.779e-02 7.551e-04      0.0007875
#> tau               3.064e-02 5.099e-03 1.019e-04      0.0001627
#> total_infections  1.286e+03 8.798e+01 1.758e+00      3.7372707
#> lnlike           -3.987e+03 1.483e+02 2.963e+00      6.3895087
#> prior_prob       -2.078e+03 1.399e+02 2.794e+00      5.9953367
#> 
#> 2. Quantiles for each variable:
#> 
#>                        2.5%        25%        50%        75%      97.5%
#> mu                1.943e+00  2.127e+00  2.226e+00  2.316e+00  2.490e+00
#> sigma1            9.589e-02  1.014e-01  1.047e-01  1.074e-01  1.123e-01
#> error             1.090e+00  1.137e+00  1.162e+00  1.186e+00  1.242e+00
#> tau               2.053e-02  2.722e-02  3.056e-02  3.411e-02  4.052e-02
#> total_infections  1.125e+03  1.225e+03  1.283e+03  1.344e+03  1.468e+03
#> lnlike           -4.291e+03 -4.090e+03 -3.982e+03 -3.885e+03 -3.714e+03
#> prior_prob       -2.357e+03 -2.174e+03 -2.074e+03 -1.981e+03 -1.815e+03


## Plot the MCMC trace using the `bayesplot` package
color_scheme_set("viridis")
p_theta_trace <- mcmc_trace(list_chains1)
print(p_theta_trace)
```

and at the infection histories:

```
## Extract infection history chain
inf_chain <- all_chains$inf_chain

## Look at inferred attack rates
p_ar <- plot_attack_rates(inf_chain, titre_dat, strain_isolation_times, pad_chain=FALSE,
                          plot_den = TRUE,prior_pars=list(prior_version=prior_version, 
                                                          alpha=par_tab[par_tab$names=="alpha","values"],
                                                          beta=par_tab[par_tab$names=="beta","values"])) 
print(p_ar)
#> Warning in regularize.values(x, y, ties, missing(ties)): collapsing to
#> unique 'x' values

#> Warning in regularize.values(x, y, ties, missing(ties)): collapsing to
#> unique 'x' values

#> Warning in regularize.values(x, y, ties, missing(ties)): collapsing to
#> unique 'x' values

#> Warning in regularize.values(x, y, ties, missing(ties)): collapsing to
#> unique 'x' values
```

```
## Calculate convergence diagnostics and summary statistics on infection histories
## Important to scale all infection estimates by number alive from titre_dat
n_alive <- get_n_alive_group(titre_dat, strain_isolation_times,melt=TRUE)

## This function generates a number of MCMC outputs
ps_infhist <- plot_posteriors_infhist(inf_chain=inf_chain, 
                                      years=strain_isolation_times, 
                                      n_alive=n_alive)
#> Calculating by time summaries...
#> Done
#> Calculating by individual summaries...
#> Done


## Posterior mean, median, 95% credible intervals and effective sample size
## on per time attack rates
print(head(ps_infhist[["estimates"]]$by_year))
#> Empty data.table (0 rows and 7 cols): j,group,mean,median,lower_quantile,upper_quantile...

## Posterior mean, median, 95% credible intervals and effective sample size
## on per individual total number of infections
print(head(ps_infhist[["estimates"]]$by_indiv))
#>    i      mean median lower_quantile upper_quantile effective_size
#> 1: 1 12.407585     12             10             16       1525.336
#> 2: 2  7.752495      8              6             10       2054.784
#> 3: 3  9.282635      9              7             12       1608.925
#> 4: 4  8.418762      8              6             11       1637.303
#> 5: 5  9.674651     10              7             13       1873.818
#> 6: 6  9.214371      9              7             12       1756.417

## Check convergence of infection history summary statistics
## MCMC trace plots of attack rates
print(ps_infhist[["by_time_trace"]][[1]])
```

```
## MCMC trace plots of total number of infections per individual
print(ps_infhist[["by_indiv_trace"]][[1]])
```

```
## Distribution of total number of infections
print(ps_infhist[["indiv_infections"]])
```

```
## Check for agreement between inferred cumulative infection histories 
## for some individuals
p_indiv_inf_hists <- generate_cumulative_inf_plots(inf_chain,indivs=1:9,pad_chain=FALSE,
                                                  strain_isolation_times = strain_isolation_times,
                                                  number_col=3)
print(p_indiv_inf_hists[[1]])
```

```
## Posterior probability that infections occured at given times per individual
print(p_indiv_inf_hists[[2]])
```

Mixing can sometimes be very poor for per-time attack rates when adjacent times are highly correlated. This is often the case when the amount of data relatively poor. A cruder time resolution (eg. per two years) may be advisable, and mixing may benefit from increasing the `hist_switch_prob` and `hist_sample_prob` parameters in the `mcmc_pars` list in `run_MCMC`. `hist_sample_prob` determines how frequently the MCMC sampler uses a proposal step that swaps the a proportion `hist_switch_prob` of individual’s infection states between two time points.

Users may also easily check the inferred antibody landscapes at the time each sample was taken. Black dots show observations, shaded regions and black line show 95%, 50% credible intervals and posterior median.

```
## get_titre_predictions expects only a single MCMC chain, so
## subset for only one chain
chain <- as.data.frame(all_chains$theta_chain)
chain1 <- chain[chain$chain_no == 1,]
inf_chain1 <- inf_chain[inf_chain$chain_no == 1,]

titre_preds <- get_titre_predictions(chain = chain1, 
                                     infection_histories = inf_chain1, 
                                     titre_dat = titre_dat, 
                                     individuals = unique(titre_dat$individual),
                                     antigenic_map = antigenic_map, 
                                     par_tab = par_tab,expand_titredat=FALSE)
#> Creating model solving function...
#> 
to_use <- titre_preds$predictions
print(head(to_use))
#>   individual  DOB virus titre samples run group    lower lower_50   median
#> 1          1 1934  1968     4    2009   1     1 2.962815 3.731254 4.148012
#> 2          1 1934  1975     3    2009   1     1 1.853615 2.609832 3.157147
#> 3          1 1934  1979     3    2009   1     1 2.373041 3.167113 3.776016
#> 4          1 1934  1989     4    2009   1     1 2.442537 3.257086 3.785972
#> 5          1 1934  1995     5    2009   1     1 3.575836 4.541814 4.926535
#> 6          1 1934  2002     5    2009   1     1 4.218560 4.881464 5.365499
#>   upper_50    upper      max
#> 1 4.659237 5.526025 5.254333
#> 2 3.633447 4.697035 4.396225
#> 3 4.259451 5.198850 3.416123
#> 4 4.448317 5.629520 4.769613
#> 5 5.201654 6.400920 4.815301
#> 6 5.730106 6.431701 4.524252

## Using ggplot
titre_pred_p <- ggplot(to_use[to_use$individual %in% 1:9,])+
  geom_ribbon(aes(x=virus,ymin=lower, ymax=upper),fill="gray90")+
  geom_ribbon(aes(x=virus,ymin=lower_50, ymax=upper_50),fill="gray70")+
  geom_line(aes(x=virus, y=median))+
  geom_point(aes(x=virus, y=titre))+
  coord_cartesian(ylim=c(0,8))+
  ylab("log titre") +
  xlab("Time of virus circulation") +
  theme_classic() +
  facet_wrap(~individual)
titre_pred_p
```

#### Further analyses

Figures in the main text can be readily generated from the MCMC output from above. The source code to generate these figures has been hidden, but can be found in the original .Rmd file for this vignette.

First, we are interested in calculating the number of infections experienced by individuals over time as a function of their age. We see that individuals are infected less frequently as they become older.

Given the sparsity of data here, the default attack rate plot is difficult to interpret. Below is an alternative visualisation of the attack rate, with the 95% and 50% credible intervals shown in red, the posterior median shown in black and the posterior maximum likelihood estimate shown as a dashed green line.

```
## Find samples that were in both theta and inf hist chains
chain <- as.data.frame(all_chains$theta_chain)
intersect_samps <- intersect(unique(inf_chain$sampno), unique(chain$sampno))
chain <- chain[chain$sampno %in% intersect_samps,]

## Find the parameter values that gave the highest posterior probability
which_mle <- chain[which.max(chain$lnlike),c("sampno","chain_no")]

## Take subset of chain for computational speed, as do not need all samples
samps <- unique(inf_chain[,c("sampno","chain_no")])
n_samps <- sample(1:nrow(samps), 100)
samps <- samps[n_samps,]
samps <- rbind(samps, which_mle) ## Plus MLE estimate
## Append the MLE estimate, note that this is max(sampno)

## Create new index variables for simplicity
samps$sampno1 <- 1:nrow(samps)
samps$chain_no1 <- 1

## Inner join to return only our subset of samples
## Reformat sampno and chain_no identifiers so that code
## sees samples as coming from one chain
inf_chain <- merge(inf_chain, samps, by=c("sampno","chain_no"))
inf_chain <- inf_chain[,c("sampno1","chain_no1","i","j","x")]
colnames(inf_chain)[1:2] <- c("sampno","chain_no")
inf_chain <- pad_inf_chain(inf_chain)
## Rename columns to be more informative

## Column names expected by code below
colnames(inf_chain) <- c("sampno","chain_no","individual","year","infected")

## Data on which strains belong to which cluster
cluster_path <- system.file("extdata", "fonville_clusters.csv", package = "serosolver")
clusters <- read.csv(file = cluster_path, stringsAsFactors=FALSE)
clusters <- clusters[clusters$year <= sample_year,]

## j=1 corresponds to the year 1968
inf_chain$year <- inf_chain$year + 1967

## Merge cluster data and infection history data
inf_chain <- merge(inf_chain, clusters[,c("year","cluster1")],by="year")

## Calculate ages and age groups of all individuals
titre_dat$age <- max(strain_isolation_times) - titre_dat$DOB
titre_dat$age_group <- cut(titre_dat$age,breaks=c(0,20,100),include.lowest=TRUE)
ages <- unique(titre_dat[,c("individual","age_group","DOB","age")])

## Merge infection histories with individual data
inf_chain<- merge(inf_chain, data.table(ages), by=c("individual"))

## Alive status for each individual for each time,
## only interested in individuals that were alive 
## when a virus circulated
inf_chain$alive <- inf_chain$DOB <= inf_chain$year
inf_chain <- inf_chain[inf_chain$alive,]
```

Finally, inferring individual infection histories allows us to investigate age-specific patterns of incidence. Here, we show the proportion of individuals that were infected at least once within a single antigenic cluster, finding that clusters that circulate for longer tend to infect a far higher proportion of the population. Furthermore, we see that a far higher proportion of the younger age group is infected in more recent years.

### Simulation recovery

We finish the vignette by presenting a simulation-recovery experiment to test the ability of the framework to recover known infection histories and antibody kinetics parameters using simulated data that matches the real dataset.

#### Extract MLE parameters from fits

We simulate infection histories and antibody titre data based on the “real” parameters inferred from fitting the model above. First, we extract the maximum posterior probability antibody kinetics parameters and attack rates.

```
## Read in MCMC chains from fitting
all_chains <- load_mcmc_chains(location=chain_path_real,thin=1,burnin=100000,
                               par_tab=par_tab,unfixed=FALSE,convert_mcmc=FALSE)
#> Chains detected:     5Highest MCMC sample interations: 
#> Chains detected: 
#> /tmp/Rtmpe5gKkm/temp_libpath6e556ded39fc/serosolver/extdata/cs2_real//case_study_2_1_infection_histories.csv
#> /tmp/Rtmpe5gKkm/temp_libpath6e556ded39fc/serosolver/extdata/cs2_real//case_study_2_2_infection_histories.csv
#> /tmp/Rtmpe5gKkm/temp_libpath6e556ded39fc/serosolver/extdata/cs2_real//case_study_2_3_infection_histories.csv
#> /tmp/Rtmpe5gKkm/temp_libpath6e556ded39fc/serosolver/extdata/cs2_real//case_study_2_4_infection_histories.csv
#> /tmp/Rtmpe5gKkm/temp_libpath6e556ded39fc/serosolver/extdata/cs2_real//case_study_2_5_infection_histories.csv
#> [[1]]
#> [1] 636633
#> 
#> [[2]]
#> [1] 645305
#> 
#> [[3]]
#> [1] 653492
#> 
#> [[4]]
#> [1] 639808
#> 
#> [[5]]
#> [1] 646868

## Alternative, load the included MCMC chains rather than re-running
## load(cs2_chains_real_b)
## all_chains <- cs2_chains_real_b

## Find samples that were in both theta and inf hist chains
chain <- all_chains$theta_chain
inf_chain <- all_chains$inf_chain
intersect_samps <- intersect(unique(inf_chain$sampno), unique(chain$sampno))
chain <- chain[chain$sampno %in% intersect_samps,]

## Find the parameter values that gave the highest posterior probability
which_mle <- chain[which.max(chain$lnlike),c("sampno","chain_no")]
mle_theta_pars <- chain[chain$sampno == which_mle$sampno & chain$chain_no == which_mle$chain_no,]

## Store total infections to compare later
mle_total_infs <- mle_theta_pars[,"total_infections"]
mle_theta_pars <- mle_theta_pars[,par_tab$names]
mle_inf_hist <- inf_chain[inf_chain$sampno == which_mle$sampno & inf_chain$chain_no == which_mle$chain_no,]

## Generate full infection history matrix using provided function
mle_inf_hist <- expand_summary_inf_chain(mle_inf_hist[,c("sampno","j","i","x")])
## Find number of infections per year from this infection history
no_infs <- colSums(mle_inf_hist[,3:ncol(mle_inf_hist)])

## If missing time points in simulated attack rates
if(length(no_infs) < length(strain_isolation_times)){
  diff_lengths <- length(strain_isolation_times) - length(no_infs)
  no_infs <- c(no_infs, rep(0, diff_lengths))
}

## Find attack rate per year
n_alive <- get_n_alive(titre_dat, strain_isolation_times)
attack_rates <- no_infs/n_alive
```

Functions are provided to simulate antibody titre data under a given serosurvey design. The antibody kinetics parameters and attack rates estimated above are used to simulate titres from the model. The `simulate_data` function is well documented, and users should refer to the help file to customise the simulated serosurvey design.

```
set.seed(1234)

sim_par_tab <- par_tab
sim_par_tab$values <- as.numeric(mle_theta_pars)
sim_par_tab[sim_par_tab$names %in% c("alpha","beta"),"values"] <- c(1/3,1/3)

age_min <- 2009 - max(titre_dat$DOB)
age_max <- 2009 - min(titre_dat$DOB)
n_indiv <- length(unique(titre_dat$individual))
dat <- simulate_data(par_tab=sim_par_tab,
                     n_indiv=n_indiv, 
                     buckets=resolution, 
                     strain_isolation_times=strain_isolation_times,
                     sampling_times=2009, 
                     nsamps=1, 
                     antigenic_map=antigenic_map, 
                     age_min=age_min,
                     age_max=age_max,
                     attack_rates=attack_rates,
                     repeats=1)
#> Simulating data


## Inspect simulated antibody titre data and infection histories
sim_titre_dat <- dat[["data"]]
sim_infection_histories <- dat[["infection_histories"]]

## Store total infections to compare later
actual_total_infections <- sum(sim_infection_histories)

plot_data(sim_titre_dat, sim_infection_histories, strain_isolation_times,n_samps = 5)
```

```
## Use titres only against same viruses tested in real data
viruses <- unique(titre_dat$virus)
sim_titre_dat <- sim_titre_dat[sim_titre_dat$virus %in% viruses, ]
sim_ages <- dat[["ages"]]
sim_titre_dat <- merge(sim_titre_dat, sim_ages)
sim_ar <- dat[["attack_rates"]]
```

#### Simulation fitting

Once these simulated data have been generated, the work flow becomes exactly the same as with the real data above.

```
filename <- "case_study_2_sim"

## Distinct filename for each chain
no_chains <- 5
filenames <- paste0(filename, "_",1:no_chains)

## Create the posterior solving function that will be used in the MCMC framework 
model_func <- create_posterior_func(par_tab=sim_par_tab,
                                titre_dat=sim_titre_dat,
                                antigenic_map=antigenic_map,
                                version=prior_version) # function in posteriors.R
#> Creating posterior solving function...
#> 

## Generate results in parallel
res <- foreach(x = filenames) %dopar% {
  ## Not all random starting conditions return finite likelihood, so for each chain generate random
  ## conditions until we get one with a finite likelihood
  start_prob <- -Inf
  while(!is.finite(start_prob)){
    ## Generate starting values for theta
    start_tab <- generate_start_tab(par_tab)
    ## Generate starting infection history
    start_inf <- setup_infection_histories_new_2(sim_titre_dat, strain_isolation_times, space=3,titre_cutoff=4)
    start_prob <- sum(model_func(start_tab$values, start_inf)[[1]])
  }
  
  res <- run_MCMC(par_tab = start_tab, 
                  titre_dat = sim_titre_dat,
                  antigenic_map = antigenic_map,
                  start_inf_hist = start_inf, 
                  mcmc_pars = c("iterations"=500000,"adaptive_period"=100000,"thin"=1000,
                                "thin_hist"=1000,"save_block"=1000,
                                "hist_switch_prob"=0.8, "hist_sample_prob"=1),
                  filename = paste0(chain_path_sim,x), 
                  CREATE_POSTERIOR_FUNC = create_posterior_func, 
                  version = prior_version)
}
```

#### Simulation analysis

MCMC chains should be checked for convergence under the usual diagnostics. We also compare the inferred posterior distributions to the known true parameter values. We see that convergence and between-chain agreement is good and that the model recovers reasonably unbiased estimates for some parameters. However, under this sampling strategy the model slightly underestimates the amount of long term antibody boosting elicited by a single infection and overestimates the total number of infections. This is driven by the contribution of the attack rate prior relative to the contribution of the likelihood (the data). Increasing the number of measured titres (for example, measure titres against 40 viruses rather than 9) or using a more informative attack rate prior would help reduce this bias.

```
## Read in the MCMC chains
## Note that `thin` here is in addition to any thinning done during the fitting
sim_all_chains <- load_mcmc_chains(location=chain_path_sim,thin=1,burnin=100000,
                               par_tab=par_tab,unfixed=FALSE,convert_mcmc=TRUE)
#> Chains detected:     5Highest MCMC sample interations: 
#> Chains detected: 
#> /tmp/Rtmpe5gKkm/temp_libpath6e556ded39fc/serosolver/extdata/cs2_sim//case_study_2_sim_1_infection_histories.csv
#> /tmp/Rtmpe5gKkm/temp_libpath6e556ded39fc/serosolver/extdata/cs2_sim//case_study_2_sim_2_infection_histories.csv
#> /tmp/Rtmpe5gKkm/temp_libpath6e556ded39fc/serosolver/extdata/cs2_sim//case_study_2_sim_3_infection_histories.csv
#> /tmp/Rtmpe5gKkm/temp_libpath6e556ded39fc/serosolver/extdata/cs2_sim//case_study_2_sim_4_infection_histories.csv
#> /tmp/Rtmpe5gKkm/temp_libpath6e556ded39fc/serosolver/extdata/cs2_sim//case_study_2_sim_5_infection_histories.csv
#> [[1]]
#> [1] 668936
#> 
#> [[2]]
#> [1] 673801
#> 
#> [[3]]
#> [1] 671807
#> 
#> [[4]]
#> [1] 674990
#> 
#> [[5]]
#> [1] 674232

## Alternative, load the included MCMC chains rather than re-running
## load(cs2_chains_sim)
## sim_all_chains <- cs2_chains_sim

theta_chain <- sim_all_chains$theta_chain
## Get the MCMC chains as a list
list_chains <- sim_all_chains$theta_list_chains
## Look at diagnostics for the free parameters
list_chains1 <- lapply(list_chains, function(x) x[,c("mu","sigma1","error",
                                                     "tau","total_infections",
                                                     "lnlike","prior_prob")])

## Gelman-Rubin diagnostics and effective sample size
print(gelman.diag(as.mcmc.list(list_chains1)))
#> Potential scale reduction factors:
#> 
#>                  Point est. Upper C.I.
#> mu                    1.016       1.04
#> sigma1                0.999       1.00
#> error                 1.000       1.00
#> tau                   1.021       1.06
#> total_infections      1.008       1.02
#> lnlike                1.010       1.02
#> prior_prob            1.009       1.02
#> 
#> Multivariate psrf
#> 
#> 1.02
print(effectiveSize(as.mcmc.list(list_chains1)))
#>               mu           sigma1            error              tau 
#>         580.2099        1569.4044        2785.2981        1129.4754 
#> total_infections           lnlike       prior_prob 
#>         666.2466         639.4231         672.1412

melted_theta_chain <- reshape2::melt(as.data.frame(theta_chain), id.vars=c("sampno","chain_no"))
estimated_pars <- c(sim_par_tab[sim_par_tab$fixed == 0,"names"],"total_infections")
melted_theta_chain <- melted_theta_chain[melted_theta_chain$variable %in% estimated_pars,]
colnames(melted_theta_chain)[3] <- "names"

add_row <- data.frame("total_infections",actual_total_infections,0,0.1,0,10000,0,0,1)
colnames(add_row) <- colnames(sim_par_tab)
sim_par_tab1 <- rbind(sim_par_tab, add_row)

ggplot(melted_theta_chain) + 
  geom_density(aes(x=value,fill=as.factor(chain_no)),alpha=0.5) +
  geom_vline(data=sim_par_tab1[sim_par_tab1$fixed == 0,],aes(xintercept=values),linetype="dashed") +
  facet_wrap(~names,scales="free") + 
  theme_classic() +
  theme(legend.position="bottom")
```

Recovery of known attack rates is also reasonably accurate, though the constraint of the posterior distibution is quite low for many years where identifiability is poor. Again, more titre data or more individuals would improve inferential power. One particularly reassuring plot is the comparison of known individual cumulative infection histories (the cumulative sum of infections over time for an individual) against the estimated posterior distribution of cumulative infection histories. We see that the 95% credible intervals capture the true cumulative infection histories in almost all cases.

```
## Extract infection history chain
inf_chain <- sim_all_chains$inf_chain

## Look at inferred attack rates
p_ar <- plot_attack_rates(inf_chain, sim_titre_dat, strain_isolation_times, pad_chain=FALSE,
                            plot_den = TRUE,prior_pars=list(prior_version=prior_version, 
                                                          alpha=par_tab[par_tab$names=="alpha","values"],
                                                          beta=par_tab[par_tab$names=="beta","values"]))  +
  geom_point(data=sim_ar,aes(x=year,y=AR),col="purple")
print(p_ar)
```

```
## Calculate convergence diagnostics and summary statistics on infection histories
## Important to scale all infection estimates by number alive from titre_dat
sim_n_alive <- get_n_alive_group(sim_titre_dat, strain_isolation_times,melt=TRUE)

## This function generates a number of MCMC outputs
ps_infhist <- plot_posteriors_infhist(inf_chain=inf_chain, 
                                      years=strain_isolation_times, 
                                      n_alive=sim_n_alive)
#> Calculating by time summaries...
#> Done
#> Calculating by individual summaries...
#> Done

## Check convergence of infection history summary statistics
## MCMC trace plots of attack rates
print(ps_infhist[["by_time_trace"]][[1]])
```

```
## MCMC trace plots of total number of infections per individual
print(ps_infhist[["by_indiv_trace"]][[1]])
```

```
## Check for agreement between inferred cumulative infection histories 
## for some individuals
p_indiv_inf_hists <- generate_cumulative_inf_plots(inf_chain,indivs=1:9,pad_chain=FALSE,
                                                   real_inf_hist=sim_infection_histories,
                                                  strain_isolation_times = strain_isolation_times,
                                                  number_col=3)
print(p_indiv_inf_hists[[1]])
```

```
## Posterior probability that infections occured at given times per individual
print(p_indiv_inf_hists[[2]])
```
